## Supporting Information for "Adapting genetic algorithms for artificial evolution of visual patterns under selection from wild predators"

***SI1: Additional methodological information for case study field trials***

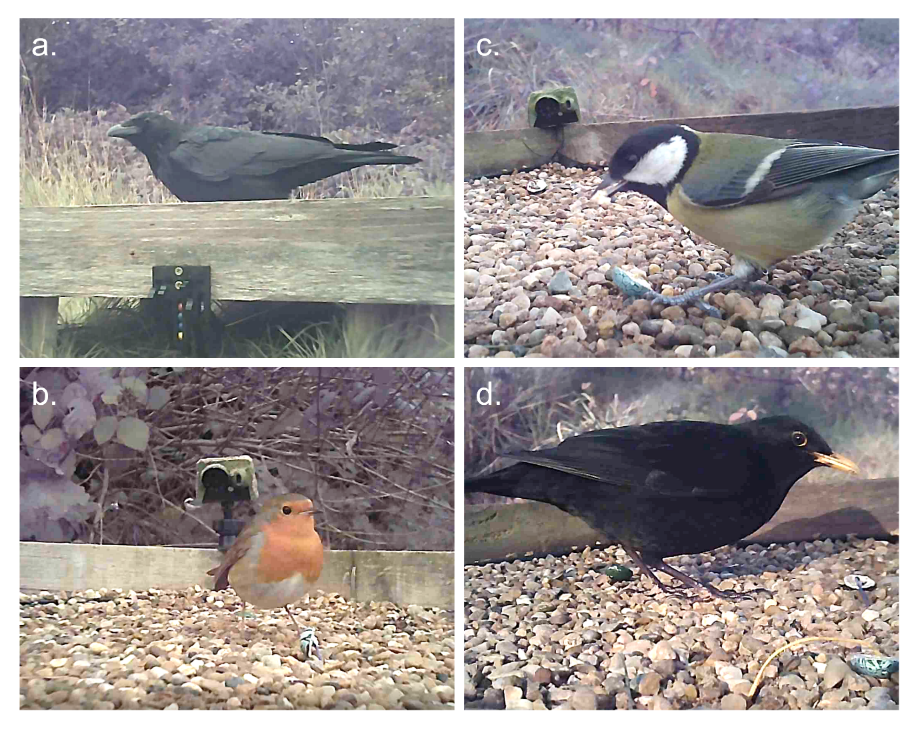

**S1 Fig. Evidence of predation from a crow (*C. corone*) in run 1 (a), and a robin (*E. rubecula*), great tit (*P. major*) and blackbird (*T. merula*) in run 2 (b-d).** Still frames extracted from footage captured by camera traps.

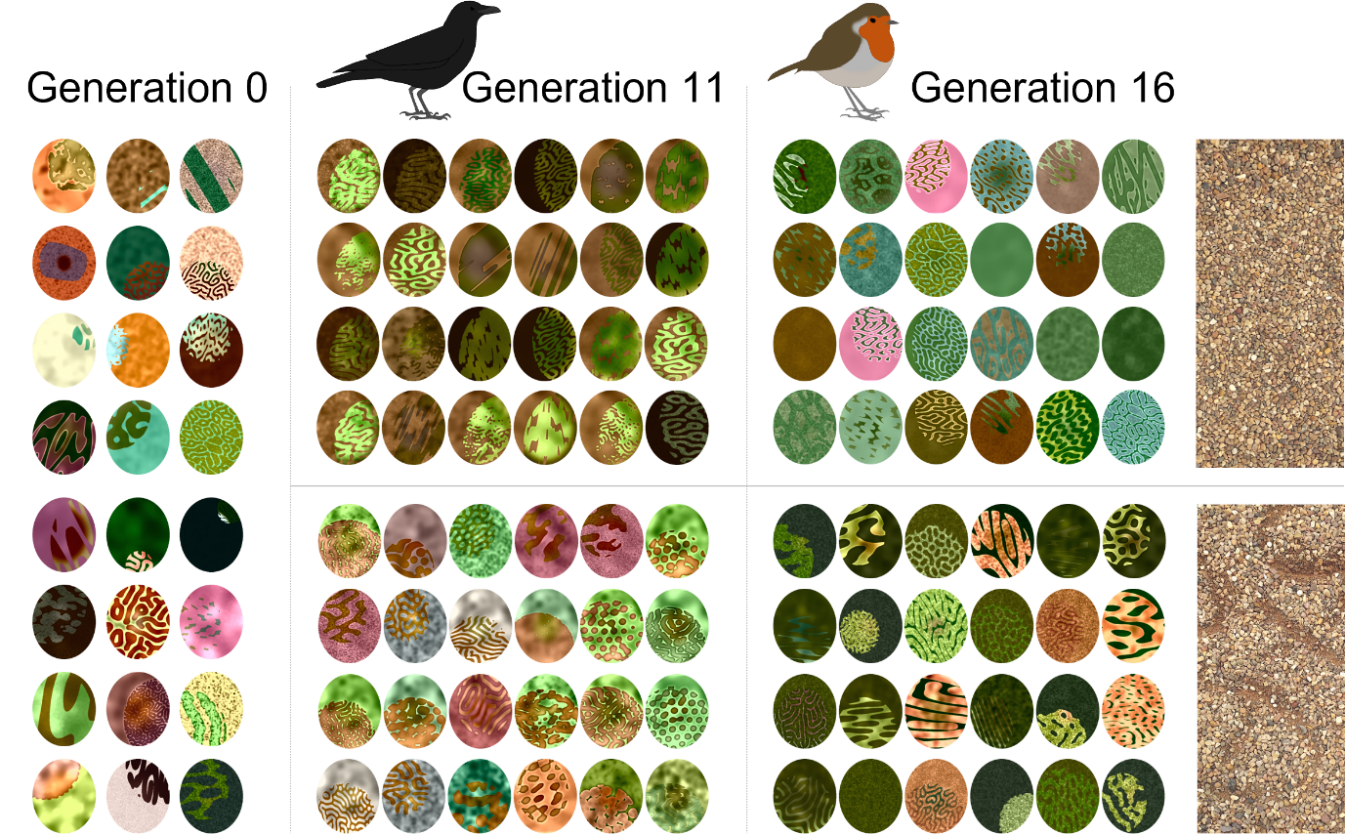

**S2 Fig. Prey patterns in the field trials: generation 0 (left) and the final generation in runs 1 (middle) and 2 (right) for populations shown on smooth (top) and furrowed backgrounds (bottom).**

*Modifications of CamoEvo toolbox for field trials*

Several changes to the CamoEvo pattern generation tools were implemented to facilitate the use of these patterns to make physical targets:

1) A custom oval target shape was created, which was wider than the target size, so that the printed pattern could more easily be transferred to the physical target and cover it completely.
2) An altered version of the base animal pattern generator was used, changing the shading such that it originated from the centre to the edge, to allow for countershading (S3 Fig).
3) Other modifications were designed to automate the task of assigning each item to a tray and position, so that each pattern could more easily and reliably be related to its fate in the field. Each target was randomly assigned to one of 6 groups of 4, corresponding to each tray. Then for each group, the targets were randomly assigned a colour (red, green, blue or yellow), indicating the colour they would correspond to in the mechanical timing gate, and thus their position (from left to right) in the tray. Each target pattern was given a code indicating the population name, repeat number, generation and colour (e.g. Tr1.1.0.R = Treatment1, repeat 1, generation 0, red). The targets were then scaled and positioned on a printable A4 sheet, first by grouping with another custom script and then rescaling and printing with INKSCAPE (INKSCAPE, https://inkscape.org/). A custom script also pseudo-randomly assigned positions in the trays for each item, to facilitate setting up in the field. According to this script, each tray was divided into a 7x7 square grid, and prey position was determined in accordance with the following rules: no prey were placed in the squares along the edges of the tray (as these areas could be more difficult to view), and prey could not be placed in adjacent squares in the grid, to reduce clumping.
4) After each generation was complete, the order of attack, representing the fitness of these targets, was manually entered into a table on ImageJ and re-assigned to the original target names, allowing the fitness values to be more easily fed back into ImageGA.

All custom pattern generation, print sheet and data entry scripts used here can be found in the GitHub repository: <https://github.com/GeorgeHancock471/CamoWild_Repository_CE.v1.2>

Scripts for print sheets and data generation have been further modified for ease of use in CamoEvo V2.0 (see CamoPrint): <https://github.com/GeorgeHancock471/CamoEvo-v2.0-2022_Plugins>

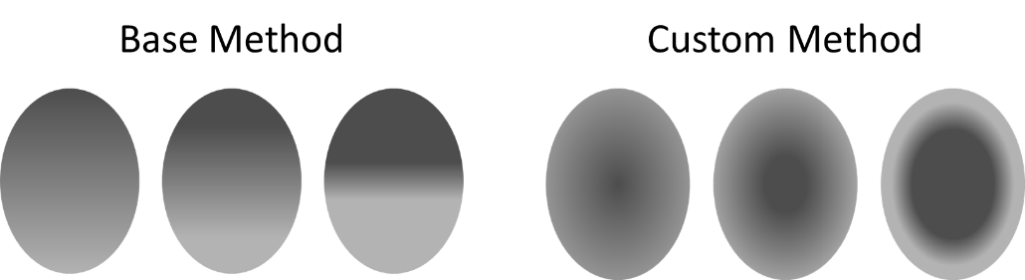

**S3 Fig. Altered target shading method, showing the base method (top-down) on the left and the custom method (centre-edge) on the right.**

*Further details on image analyses to quantify pattern properties*

All prey patterns were printed on waterslide paper and transferred to sheets of white tack to be photographed, capturing their colours as shown to predators in the experiment. Photographs were taken outdoors in diffuse lighting conditions, with a SONY α6000 camera converted to full spectrum and a custom-built 52mm lens. Images were taken in RAW format, with the aperture set to f8, ISO 400 and white balance set to cloudy. Two filters were used, the camera’s own visible light pass filter and a Baader U-Venus filter (transmitting 320-380nm; Baader Planetarium, Germany), to photograph the patterns in both the visible light and UV ranges. All photographs included a scale bar and pair of grey standards made of Zenith Polymer sintered PTFE sheet, reflecting 93 and 7% of light at all wavelengths between 200 and 700nm.

Analyses were carried out using the Multispectral Imaging (MICA) and QCPA plugins [1,2] for ImageJ [3]. Throughout, target populations in the first and second runs were analysed separately, as they were targeted by predators with different visual systems: violet-sensitive crows and ultraviolet-sensitive robins respectively [4]. Normalised multispectral images were converted to cone catches for the appropriate avian visual model (the peafowl *Pavo cristatus*, representing violet-sensitive species, or the bluetit, *Cyanistes caeruleus*, representing UV-sensitive species). Prey patterns were then processed following the QCPA framework: regions of interest (ROIs) were drawn around the prey patterns using the ellipse tool in ImageJ, then each of these underwent spatial acuity modelling with a Gaussian filter and RNL ranked filtering for edge reconstruction. For acuity correction, images were scaled to be viewed at a distance of 53cm, the width of the trays, as a proxy for the maximum distance at which prey could be seen, and different spatial acuity values were applied for prey patterns from the different sets of experiments: 30 cycles/degree (cpd) for prey viewed by crows, based on similar measures for other corvids, rooks (*Corvus frugilegus*) and magpies (*Pica pica*) [5,6], and 6 cpd [6,7] for prey viewed primarily by a robin. To measure pattern, a granularity analysis based on Fast Fourier bandpass filters was run on the double cone catch layer, representing the avian luminance channel, of each processed prey selection. This technique measures pattern energy at each frequency band, following methods previously used in the study of animal patterns (e.g. [8,9]). Frequency bands started from 2 pixels, moving in √2 increments, until reaching the size closest to the target size in images scaled by the QCPA framework after acuity correction (362 pixels for prey in run 1, 64 pixels for run 2). Images were also converted to the Receptor Noise-Limited (RNL) chromaticity space, to obtain the coordinates of the average colour for each prey pattern in the RNL space. For all image analyses, conservative estimates of Weber fractions were used, following default values in the QCPA toolbox and previous work: 0.05 for chromatic variables and 0.1 for luminance ([9,10].

To compare the prey patterns to the backgrounds they were viewed against, and so assess camouflage efficacy, the gravel trays were also photographed with the same equipment and in the same conditions as the prey. Gravel colour and brightness varied greatly between dry and wet conditions, so trays with a smooth surface were photographed both dry and wet (N=5 for each condition). Images were scaled to 25 pixels/mm, a rectangular selection corresponding to a 15x15cm area was selected in the centre of each image, and these selections were processed and analysed as described above.

Several metrics of difference between prey patterns and their backgrounds were then computed. Mean chromatic difference (ΔS) was taken as the Euclidean distance between the average prey colour and background colour in the RNL space. Achromatic, or luminance, difference (ΔL) was calculated as the natural logarithm of the ratio between the mean double cone catch values of the prey and tray backgrounds, divided by a Weber fraction of 0.1, matching other outputs of the QCPA framework [11]. A measure of pattern difference was obtained by comparing curves describing the pattern energy measured at each frequency band by the granularity analyses, for each prey and the tray backgrounds, using the “Pattern and Luminance Difference Calculator” tool in the MICA toolbox plugins.

As the targets are patterned, and the background itself is somewhat heterogeneous, comparing mean colours provides a relatively crude measure of colour-matching. To improve on this, we also calculated a second measure of colour difference, which accounts for the diversity of colours in the targets and backgrounds. After undergoing acuity control and RNL filtering in the QCPA framework, target and background images were converted into colour maps in the RNL space, using the “Create RNL maps from ROIs” tool in the QCPA toolbox [2]. For the background, a single combined map was generated from the damp and dry background images described above, using the “Combine RNL maps” tool, and each target was compared to that map. A custom script (written by GRAH) then calculated the colour difference (in ΔS) between each target colour and the perceptually closest background colour; if a target colour exists in the background, ΔS is equal to 0, and increasing values of ΔS indicate larger differences in colour. To reduce the influence of very rare colours, a 99% threshold was applied, so that colours present in the target or background at the smallest 1% of frequencies were ignored. For each target, all pairwise colour differences thus calculated were then averaged, weighted by the frequency at which the colours appeared in the target. The resulting ΔS value is hereafter referred to as the weighted average colour difference.

In addition, target edge disruption was measured, using the “GabRat” method, previously shown to be a strong predictor of target detectability [12]. This technique uses Gabor filters to quantify the intensity of perceived edges running orthogonal to the outline of a target, relative to its real edges; higher GabRat values indicate higher levels of disruption to edge perception, facilitating disruptive camouflage. As we could not quantify edge disruption of each target in the exact location it was viewed in by predators in the field, we instead used the sRGB images of prey patterns produced by the CamoEvo toolbox, converted these to human LAB space, and calculated GabRat values of the target outline in the L channel using tools in the MICA toolbox (with angles = 6, sigma = 3.0, gamma =1.0 and freq = 2.0), against a plain grey background, matching the combined average L values of targets in the final generations of each experimental run.

Camouflage metrics were then analysed using separate linear models for each field run, with generation and background type (smooth or furrowed) as interacting fixed effects. Model assumptions were checked using diagnostic plots, and dependent variables were transformed as needed to fit these assumptions. Full models were simplified using stepwise model simplification with ANOVAs. To allow for non-linear trends in camouflage metrics across generations, generation was initially fitted as a second-degree polynomial in all models, and the value of including this polynomial term in final models was tested using ANOVAs; final model results only include the polynomial term if it was significant. See full analysis code in supplementary material.

***SI 2: Additional information for screen-based search tasks***

Screen-based search tasks with human volunteers were used to replicate the field trials, and more easily test the impact of changing the prey pattern colour space.

*Methods*

*Task design using CamoEvo*

Each participant viewed a single population of targets over 16 generations, shown on images of the same backgrounds as in the field, either smooth or furrowed. All parameters were selected to replicate the field experiment as closely as possible. Target shapes and patterns were modified using CamoEvo’s alpha and shader layers to mimic the same 3D appearance and match the exact shape of the physical prey items in the field. Additionally, a drop shadow script was added to the background images, by darkening a feathered area behind the target, of the same, but enlarged, shape (S4 Fig; see code in GitHub repository).

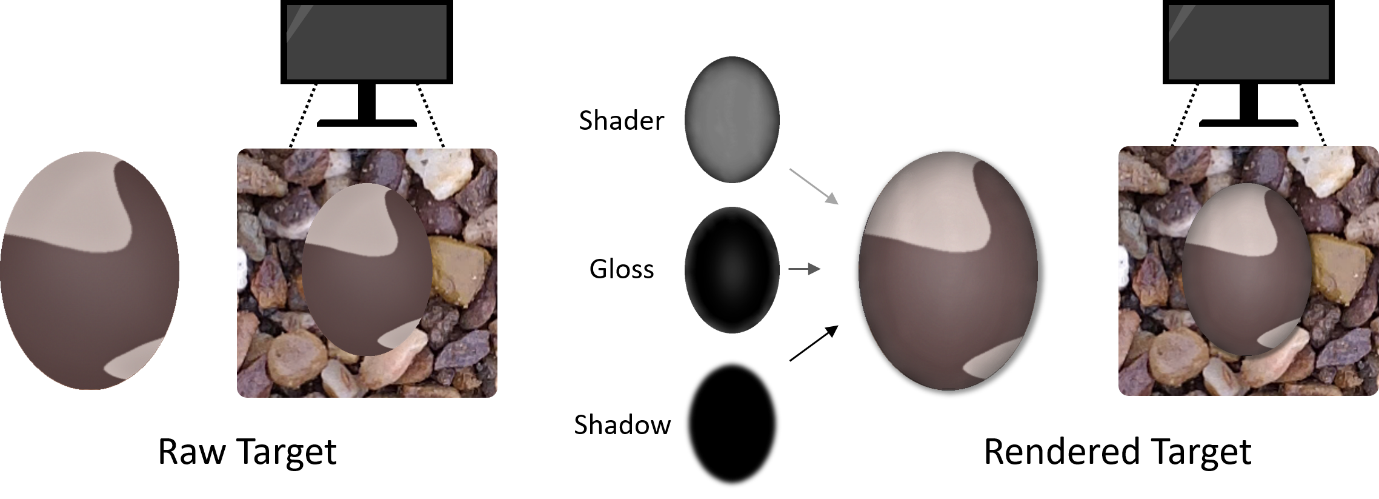

**S4 Fig. Target image modification for screen-based replicate experiments.** Unmodified target (left), and target as viewed by volunteers (right), after shadow and gloss layers have been applied to the target and a drop shadow has been applied to the background image.

Background images were based on photographs of experimental trays used in the field, taken outside in diffuse lighting conditions with a high-resolution smartphone (A202, ASUS, Taiwan). A scale bar and reflectance standards reflecting 93 and 7% of light at all wavelengths between 350 and 700nm were included in the photographs to normalise the images to ambient light levels. Images were taken in non-linear JPG format, but a linearisation model was built for this camera, using a photograph of a colour chart (ColorChecker® Classic, X-Rite, USA) and methods in the MICA toolbox [1]. Images were linearised and normalised, then converted to human CIE XYZ space using custom-built code (written by JT), and back into sRGB images using the “Make Presentation Image” tool in the MICA toolbox (with power set to 0.41 and a maximum value of 0.6). The resulting RGB images were then darkened, by subtracting 50 from the RGB stacks using the “Math” tool in ImageJ. These settings and manipulations were selected to provide realistic background images that closely resembled the trays as seen in the field. To match the field experiment, with variable weather conditions, the tray was photographed dry and damp, with both the smooth and furrowed background treatment. RGB images made from these four photographs were each cropped to 3000x3000 pixels and rotated 0, 90 and 180°, providing six background images (three dry and three damp) per background treatment.

For every generation of prey, participants viewed these six images in a random order, each containing four prey to find. Prey were scaled to 133 pixels, keeping their size relative to the tray image consistent with the field, and positioned pseudo-randomly, following the same rules as in the field. Targets in the “full colour space” populations occupied the same colour space as prey in the field (henceforth the full colour space); for those in the “narrow colour space” populations, a narrower colour space was selected, corresponding to ±2 standard deviations around the mean colour of the background images used in the search task (luminance range 11.1515 to 60.4960, A range 1.0739 to 11.2990, B range -1.3652 to 26.3401; S5 Fig). Participants were given a maximum of 15 seconds per slide to click on all the targets as they detected them. When a participant correctly located a target, it was surrounded by a green circle; missed targets were circled in red after the slide had timed out. Survival of targets into the next generation and pattern evolution were governed by the same rules as in the field, based on the order in which targets were clicked on. A total of 12 volunteers took part, each searching for a single population of prey across 16 generations yielding three replicates for every colour space and background type combination (S5 Fig)

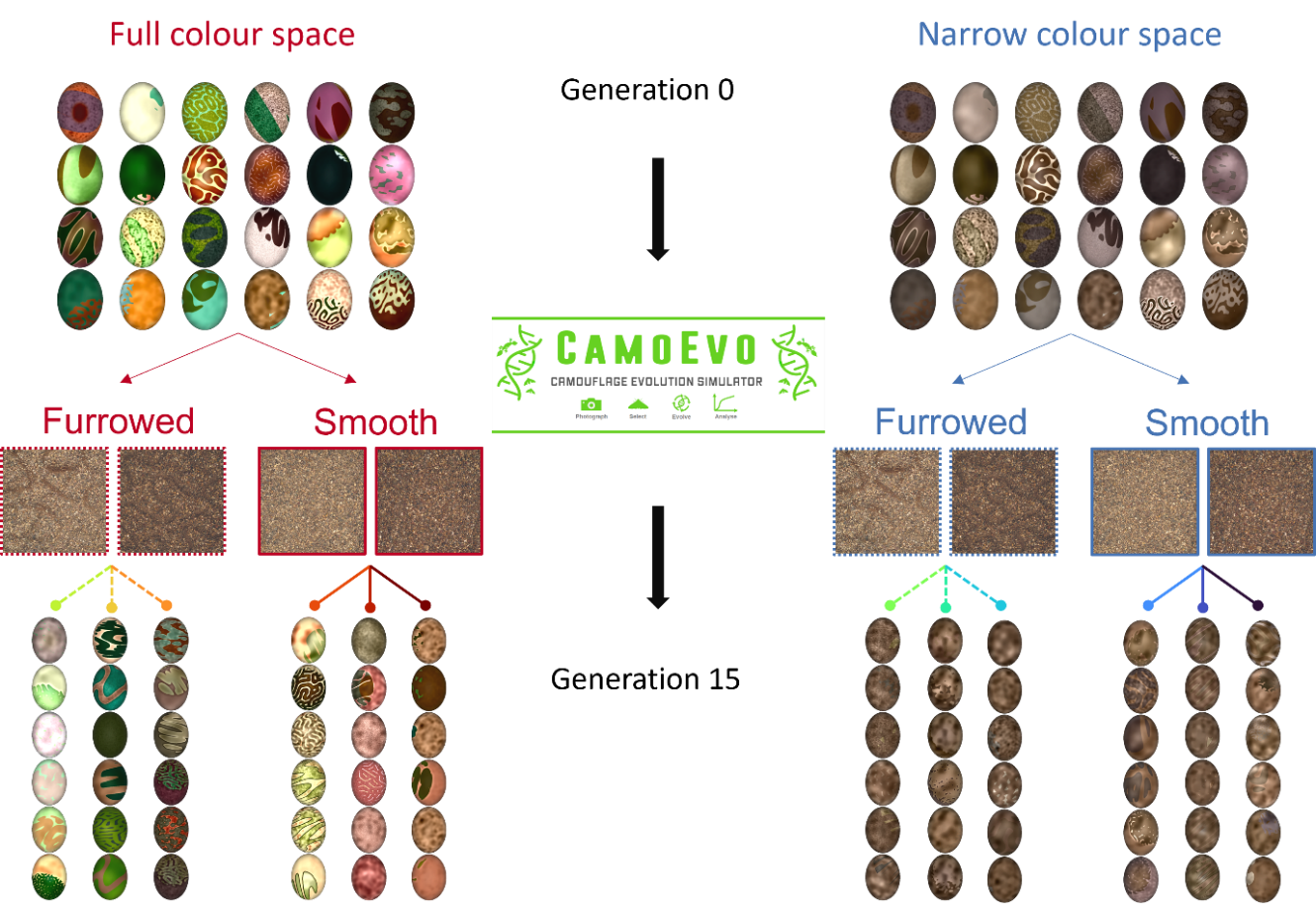

**S5 Fig. Schematic of the screen-based search tasks, showing the targets in generation 0 and 15, for populations using the full and narrow colour spaces.** Populations from both colour spaces were shown against either smooth or furrowed backgrounds, with three replicates per combination. Each replicate was completed by a single participant. For generation 15, each column of targets represents the six most successful targets (slowest to be found) for each replicate population.

*Prey pattern analysis*

The CamoEvo toolbox records the mean values in LAB space of the targets and the backgrounds they are shown against, and target GabRat in every channel, for every generation [13]. Differences in luminance between targets and their backgrounds were calculated as ΔL, and differences in colour as the Euclidean distance between the mean target and background colour in the LAB space. Edge disruption was taken as GabRat in the L channel.

In addition to the metrics computed as standard by the CamoEvo toolbox, we calculated a measure of pattern difference between the target patterns and backgrounds shown on screen, matching that used for the field trials. To do that, sRGB images of the targets and background images shown on screen were converted first to CIE XYZ, then LAB space in ImageJ, using the same custom-built tools as above, for the original manipulation of tray photographs. Granularity analyses were run on the L channel images, following the same methods as for the field prey, with the largest frequency band set to correspond to the size of the prey relative to the background in the computer-based trials (128 pixels). To match the field trials, background analyses were restricted to the smooth background only, and the results obtained for the damp and dry tray were averaged to produce a single overall background curve. The difference between the granularity curves obtained for the targets and this average background were once again calculated using the “Pattern and Luminance Difference Calculator” tool in the MICA toolbox. Finally, images of targets and smooth backgrounds in the CIE XYZ space were used to make colour maps and calculate the weighted average colour difference between all target and background colours (in ΔS), using the same tools and parameters as for the field experiments.

To account for replicate populations, camouflage metrics for targets in the screen-based tasks were analysed with linear mixed effects models, using the package ‘lme4’ [14]. Populations from each colour space were analysed separately, with initial models including generation, and background type as interacting fixed effects. Population ID was included as a random effect with a random slope for generation, following best practice for fitting random effects [15], but models with only a random intercept were also evaluated. After model simplification, final models with and without a random slope for population were compared using Akaike’s Information Criterion (AIC [16]), and the model with lowest AIC was preferred (see supporting code). For capture time, only the first targets to be clicked on in each slide (ie. the lowest-ranked targets) were included. To fit model assumptions, response variables were log- or square-root-transformed as needed (see S1 &S2 Tables). To analyse colour difference as a weighted average measure for the narrow colour space, values were rounded and models were fitted with negative binomial distributions, using the package ‘glmmTMB’ [17]. As for analyses of the field trials, model assumptions were checked with diagnostic plots, and full models were simplified using ANOVAs. Generation was initially fitted as a second-degree polynomial, but the polynomial term is only included in final model results if found significant after ANOVA tests between models with and without the polynomial. To compare experimental populations, selected by volunteers, and control populations, data were combined for each colour space, and analysed as described above, with full models initially testing for an effect of the interaction between generation and selection scenario (control vs experimental) on camouflage metrics (see supporting code).

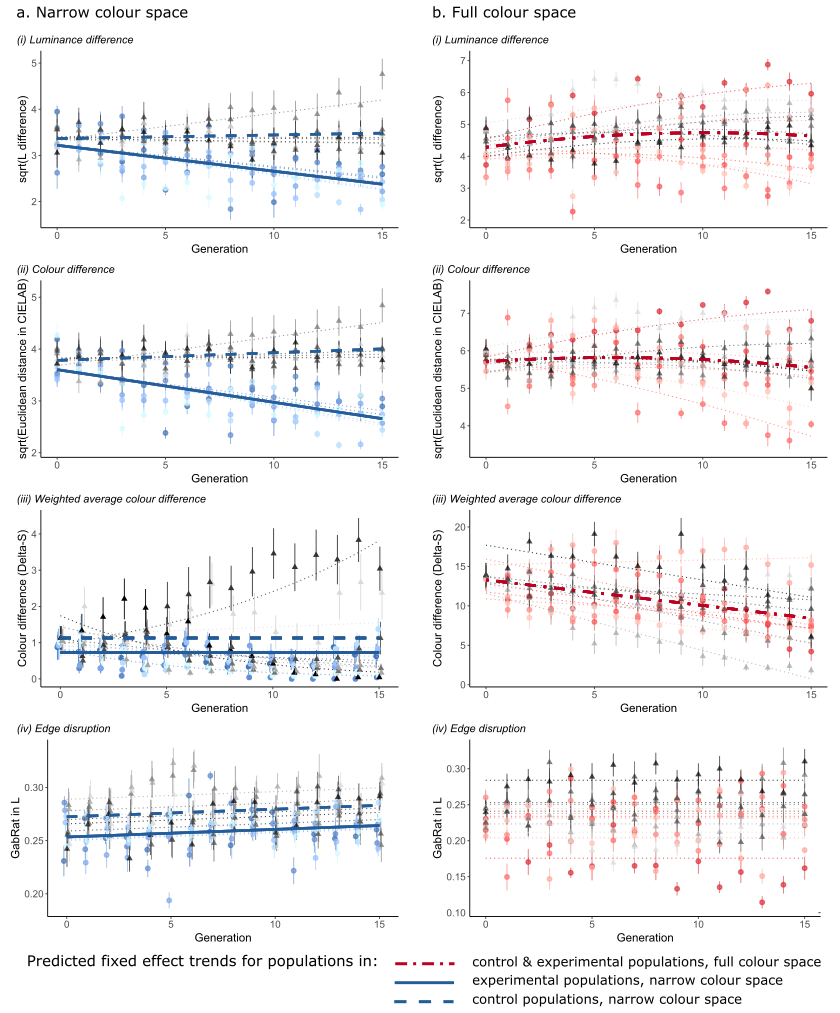

**S6 Fig. Changes in camouflage metrics in the screen-based experimental populations and control replicates, using the narrow (a) and full (b) colour spaces.** Plots show the mean and standard error of each metric per generation, coloured by population, along with predictions from the best statistical models. Coloured circles represent experimental populations, greyscale triangles control runs. Dotted lines represent population-level predictions; thicker lines show overall trends for fixed effects only.

**S1 Table. Model simplification tables for full models testing changes in metrics of camouflage efficacy in the screen-based experiment, for the narrow colour space.** Models include population-level random slopes for generation, except for luminance and colour difference (a-b), edge disruption (e) and capture time (f). Models with a polynomial term for generation are fully simplified, then the usefulness of the 2^nd^ order polynomial term is tested with an ANOVA, comparing final models with and without the polynomial terms. Significant factors are highlighted in italics.

a. Luminance difference (ΔL, square-root-transformed))

| Factor | χ^2^ | df | p |
| --- | --- | --- | --- |
| poly(Generation,2) : Background type | 4.958 | 2 | 0.0838 |
| Background type | 2.270 | 1 | 0.132 |
| *poly(Generation,2)* | *116.590* | *2* | *<0.001* |
| Effect of polynomial for generation | 2.525 | 1 | 0.112 |
| *Generation* | *114.070* | *1* | *<0.001* |

b. Colour difference (Euclidean distance between mean colours in Lab space, square-root-transformed)

| Factor | χ^2^ | df | p |
| --- | --- | --- | --- |
| poly(Generation,2) : Background type | 3.620 | 2 | 0. 164 |
| Background type | 1.568 | 1 | 0. 211 |
| *poly(Generation,2)* | *191.31* | *2* | *<0.001* |
| *Effect of polynomial for generation* | *9.247* | *1* | *0.002* |

c. Colour difference (weighted average colour difference in XYZ space, in ΔS)

| Factor | χ^2^ | df | p |
| --- | --- | --- | --- |
| *poly(Generation,2) : Background type* | *7.434* | *2* | *0.024* |
| *Effect of polynomial for generation* | *7.828* | *2* | *0.020* |

d. Pattern difference

| Factor | χ^2^ | df | p |
| --- | --- | --- | --- |
| poly(Generation,2) : Background type | 0.257 | 2 | 0.880 |
| Background type | 0.538 | 1 | 0.463 |
| *poly(Generation,2)* | *10.471* | *2* | *0.005* |
| Effect of polynomial for generation | 2.005 | 1 | 0.157 |
| *Generation* | *8.466* | *1* | *0.004* |

e. Edge disruption (GabRat in L)

| Factor | χ^2^ | df | p |
| --- | --- | --- | --- |
| poly(Generation,2) : Background type | 2.363 | 2 | 0.307 |
| Background type | 0.020 | 1 | 0.888 |
| *poly(Generation,2)* | *8.556* | *2* | *0.014* |
| Effect of polynomial for generation | 0.514 | 1 | 0.474 |
| *Generation* | *8.043* | *1* | *0.005* |

f. Capture time (log-transformed; lowest-ranked targets only)

| Factor | χ^2^ | df | p |
| --- | --- | --- | --- |
| poly(Generation,2) : Background type | 4.762 | 2 | 0.093 |
| Background type | 3.826 | 1 | 0.050 |
| *poly(Generation,2)* | *6.340* | *2* | *0.042* |
| Effect of polynomial for generation | 0 | 1 | 0.998 |
| *Generation* | *6.340* | *1* | *0.012* |

**S2 Table. Model simplification tables for full models testing changes in metrics of camouflage efficacy in the screen-based experiment, for the full colour space.** Models include population-level random slopes for generation. Models with a polynomial term for generation are fully simplified, then the usefulness of the 2^nd^ order polynomial term is tested with an ANOVA, comparing final models with and without the polynomial terms. Significant factors are highlighted in italics.

a. Luminance difference (ΔL)

| Factor | χ^2^ | df | p |
| --- | --- | --- | --- |
| poly(Generation,2) : Background type | 0.146 | 2 | 0.930 |
| Background type | 0.709 | 1 | 0.400 |
| poly(Generation,2) | 0.801 | 2 | 0.670 |

b. Colour difference (Euclidean distance between mean colours in Lab space)

| Factor | χ^2^ | df | P |
| --- | --- | --- | --- |
| poly(Generation,2) : Background type | 0.143 | 2 | 0.931 |
| Background | 0.250 | 1 | 0.617 |
| *poly(Generation,2)* | *7.255* | *2* | *0.027* |
| *Effect of polynomial for generation* | *5.854* | *1* | *0.016* |

c. Colour difference (weighted average colour difference in XYZ space, in ΔS)

| Factor | χ^2^ | df | p |
| --- | --- | --- | --- |
| *poly(Generation,2) : Background type* | *6.965* | *2* | *0.031* |
| *Effect of polynomial for generation* | *7.760* | *2* | *<0.02*1 |

d. Pattern difference

| Factor | χ^2^ | df | p |
| --- | --- | --- | --- |
| poly(Generation,2) : Background type | 4.454 | 2 | 0.108 |
| Background type | 0.020 | 1 | 0.889 |
| poly(Generation,2) | 2.788 | 2 | 0.248 |

e. Edge disruption (GabRat in L)

| Factor | χ^2^ | df | p |
| --- | --- | --- | --- |
| poly(Generation,2) : Background type | 0.481 | 2 | 0.786 |
| Background type | 0.444 | 1 | 0.505 |
| poly(Generation,2) | 2.030 | 2 | 0.363 |

f. Capture time (log-transformed; lowest-ranked targets only)

| Factor | χ^2^ | df | P |
| --- | --- | --- | --- |
| *poly(Generation,2) : Background type* | *10.832* | *2* | *0.004* |
| *Effect of polynomial for generation* | *15.824* | *2* | *<0.001* |

**S3 Table. Model simplification tables for full models testing changes in metrics of camouflage efficacy between control and experimental populations of the screen experiment, for the narrow colour space.** Models include population-level random slopes for generation, except for models of edge disruption. Models with a polynomial term for generation are fully simplified, then the usefulness of the 2^nd^ order polynomial term is tested with an ANOVA, comparing final models with and without the polynomial terms. Significant factors are highlighted in italics.

a. Luminance difference (ΔL, square-root-transformed))

| Factor | χ^2^ | df | p |
| --- | --- | --- | --- |
| *poly(Generation,2) : Scenario* | *9.650* | *2* | *0.008* |
| Effect of polynomial for generation | 0.587 | 2 | 0.746 |
| *Generation : Scenario* | *9.443* | *1* | *0.002* |

b. Colour difference (Euclidean distance between mean colours in Lab space, square-root-transformed)

| Factor | χ^2^ | df | p |
| --- | --- | --- | --- |
| *poly(Generation,2) : Scenario* | *16.968* | *2* | *<0.001* |
| Effect of polynomial for generation | 2.251 | 2 | 0.325 |
| *Generation : Scenario* | *15.557* | *1* | *<0.001* |

c. Colour difference (weighted average colour difference in XYZ space, in ΔS)

| Factor | χ^2^ | df | p |
| --- | --- | --- | --- |
| poly(Generation,2) : Scenario | 2.214 | 2 | 0.331 |
| poly(Generation,2) | 4.784 | 2 | 0.091 |
| *Scenario* | *9.352* | *1* | *0.002* |

d. Edge disruption (GabRat in L)

| Factor | χ^2^ | df | p |
| --- | --- | --- | --- |
| poly(Generation,2) : Scenario | 0.343 | 2 | 0.842 |
| *poly(Generation,2)* | *7.064* | *2* | *0.030* |
| Effect of polynomial for generation | 0.777 | 1 | 0.378 |
| *Generation* | *6.287* | *1* | *0.012* |
| *Scenario* | *8.338* | *1* | *0.004* |

**S4 Table. Model simplification tables for full models testing changes in metrics of camouflage efficacy between control and experimental populations of the screen experiment, for the full colour space.** Models include population-level random slopes for generation, except for models of edge disruption. Models with a polynomial term for generation are fully simplified, then the usefulness of the 2^nd^ order polynomial term is tested with an ANOVA, comparing final models with and without the polynomial terms. Significant factors are highlighted in italics.

a. Luminance difference (ΔL, square-root-transformed))

| Factor | χ^2^ | df | P |
| --- | --- | --- | --- |
| poly(Generation,2) : Scenario | 0.485 | 2 | 0.785 |
| Scenario | 2.202 | 1 | 0.138 |
| *poly(Generation,2)* | *8.535* | *2* | *0.014* |
| *Effect of polynomial for generation* | *6.994* | *1* | *0.008* |

b. Colour difference (Euclidean distance between mean colours in Lab space, square-root-transformed)

| Factor | χ^2^ | df | P |
| --- | --- | --- | --- |
| poly(Generation,2) : Scenario | 1.741 | 2 | 0.419 |
| Scenario | 0.059 | 1 | 0. 809 |
| *poly(Generation,2)* | *6.346* | *2* | *0.042* |
| *Effect of polynomial for generation* | *6.035* | *1* | *0.014* |

c. Colour difference (weighted average colour difference in XYZ space, in ΔS)

| Factor | χ^2^ | df | p |
| --- | --- | --- | --- |
| poly(Generation,2) : Scenario | 5.577 | 2 | 0.062 |
| Scenario | 0.339 | 1 | 0.560 |
| *poly(Generation,2)* | *14.411* | *2* | *<0.001* |
| Effect of polynomial for generation | 2.173 | 1 | 0.140 |
| *Generation* | *12.238* | *1* | *<0.001* |

d. Edge disruption (GabRat in L)

| Factor | χ^2^ | df | p |
| --- | --- | --- | --- |
| poly(Generation,2) : Scenario | 0.426 | 2 | 0.808 |
| Scenario | 2.487 | 1 | 0.115 |
| poly(Generation,2) | 3.849 | 2 | 0.146 |

***Supporting references***

6.	Martin GR. The sensory ecology of birds. Oxford: Oxford University Press; 2017. 320 p.

16.	Akaike H. Information theory and the maximum likelihood principle. In: Petrov BN, Csàki F, editors. 2nd International Symposium on Information Theory. 1973.

17.	Brooks ME, Kristensen K, van Benthem, Koen J. Magnusson A, Berg CW, Nielsen A, Skaug HJ, et al. glmmTMB balances speed and flexibility among packages for zero-inflated generalized linear mixed modeling. R J. 2017;9(2):378–400.
