## Supporting data and code for "Adapting genetic algorithms for artificial evolution of visual patterns under selection from wild predators": CamoWild_Markdown.html

CamoEvoField\_Markdown


### CamoEvoField\_Markdown

###### E.S.Briolat

###### 27th October 2023

#### Field trials

```
#load data
R1ALL<-read.csv("CamoField_Run1.csv", header=TRUE)
  #data for prey patterns in run1, from both smooth and furrowed populations
R2ALL<-read.csv("CamoField_Run2.csv", header=TRUE)
  #data for prey patterns in run2, from both smooth and furrowed populations

R1ALL_GR<-read.csv("CamoField_GabRat_Run1.csv", header=TRUE)
R2ALL_GR<-read.csv("CamoField_GabRat_Run2.csv", header=TRUE)
  #GabRat data for prey patterns in runs  and 2

#prepare data
R1ALL$Treatment <- relevel(R1ALL$Treatment, "Smooth")
R2ALL$Treatment <- relevel(R2ALL$Treatment, "Smooth")
R1ALL_GR$Treatment<-relevel(R1ALL_GR$Treatment, "Smooth")
R2ALL_GR$Treatment<-relevel(R2ALL_GR$Treatment, "Smooth")

#plotting theme
my_theme<-theme(panel.grid.major = element_blank(), panel.grid.minor = element_blank(),
                panel.background = element_blank(), axis.line = element_line(colour = "black"))
```

##### Field trials - how do camouflage metrics change across generations?

Linear models test the interacting effects of generation and background type (treatment). Generation is initially fitted as a 2nd order polynomial, and the significance of the polynomial term is tested in the final model.

###### Run 1

Colour difference between mean colours (prey against gravel backgrounds):

```
r1mod1<-lm(dS~poly(Generation,2)*Treatment, data=R1ALL)
par(mfrow=c(2,2))
plot(r1mod1)
```

```
hist(resid(r1mod1))

#model simplification
r1mod1b<-lm(dS~poly(Generation,2)+Treatment, data=R1ALL)
anova(r1mod1b, r1mod1)#significant interaction
```

```
## Analysis of Variance Table
## 
## Model 1: dS ~ poly(Generation, 2) + Treatment
## Model 2: dS ~ poly(Generation, 2) * Treatment
##   Res.Df    RSS Df Sum of Sq     F    Pr(>F)    
## 1    572 3277.3                                 
## 2    570 3191.9  2    85.398 7.625 0.0005395 ***
## ---
## Signif. codes:  0 '***' 0.001 '**' 0.01 '*' 0.05 '.' 0.1 ' ' 1
```

```
#test effect of polynomial
r1mod1c<-lm(dS~Generation*Treatment, data=R1ALL)
anova(r1mod1,r1mod1c)#polynomial is useful
```

```
## Analysis of Variance Table
## 
## Model 1: dS ~ poly(Generation, 2) * Treatment
## Model 2: dS ~ Generation * Treatment
##   Res.Df    RSS Df Sum of Sq      F   Pr(>F)   
## 1    570 3191.9                                
## 2    572 3252.8 -2   -60.856 5.4338 0.004596 **
## ---
## Signif. codes:  0 '***' 0.001 '**' 0.01 '*' 0.05 '.' 0.1 ' ' 1
```

```
summary(r1mod1)
```

```
## 
## Call:
## lm(formula = dS ~ poly(Generation, 2) * Treatment, data = R1ALL)
## 
## Residuals:
##     Min      1Q  Median      3Q     Max 
## -5.6813 -1.9386 -0.0916  1.8637  7.6866 
## 
## Coefficients:
##                                        Estimate Std. Error t value Pr(>|t|)    
## (Intercept)                              9.0244     0.1394  64.719  < 2e-16 ***
## poly(Generation, 2)1                    -4.7560     3.3466  -1.421  0.15582    
## poly(Generation, 2)2                     9.9924     3.3466   2.986  0.00295 ** 
## TreatmentFurrowed                        1.8171     0.1972   9.215  < 2e-16 ***
## poly(Generation, 2)1:TreatmentFurrowed  11.2441     4.7328   2.376  0.01784 *  
## poly(Generation, 2)2:TreatmentFurrowed -14.6684     4.7328  -3.099  0.00204 ** 
## ---
## Signif. codes:  0 '***' 0.001 '**' 0.01 '*' 0.05 '.' 0.1 ' ' 1
## 
## Residual standard error: 2.366 on 570 degrees of freedom
## Multiple R-squared:  0.1512, Adjusted R-squared:  0.1438 
## F-statistic: 20.31 on 5 and 570 DF,  p-value: < 2.2e-16
```

```
#plot

#make predictions
pred<-predict(r1mod1, newdata = R1ALL, interval = 'confidence')
R1ALLp<-cbind(R1ALL, pred)

#make plot
rep1col<-ggplot(R1ALLp, aes(Generation, dS, col=Treatment,linetype=Treatment, fill=Treatment))+
  geom_point(alpha=0.3)+
  scale_colour_manual(values=c("black","grey50"))+
  scale_fill_manual(values=c("black","grey50"))+
  facet_grid(.~Treatment)+
  scale_x_continuous(breaks=c(0,5,10,15))+
  geom_ribbon(aes(ymin=lwr,ymax=upr), alpha=0.2, colour=NA)+
  geom_line(aes(Generation, fit), size=1)+
  guides(col=FALSE)+guides(linetype=FALSE)+guides(fill=FALSE)+
  ylab("Colour difference between means (Delta-S)")+
  my_theme+
  theme(axis.title = element_text(size = 16))  +
  theme(axis.text = element_text(size = 14)) +
  theme(strip.background = element_rect( fill="grey90"))+
  theme(strip.text.x = element_blank())
```

Colour difference based on weighted average distance:

```
r1mod1d<-lm(NewMapComp_DeltaS~poly(Generation,2)*Treatment, data=R1ALL)
par(mfrow=c(2,2))
plot(r1mod1d)
```

```
hist(resid(r1mod1d))

#model simplification
r1mod1e<-lm(NewMapComp_DeltaS~poly(Generation,2)+Treatment, data=R1ALL)
anova(r1mod1d, r1mod1e)#significant interaction
```

```
## Analysis of Variance Table
## 
## Model 1: NewMapComp_DeltaS ~ poly(Generation, 2) * Treatment
## Model 2: NewMapComp_DeltaS ~ poly(Generation, 2) + Treatment
##   Res.Df   RSS Df Sum of Sq      F    Pr(>F)    
## 1    570 38996                                  
## 2    572 41380 -2   -2384.3 17.425 4.514e-08 ***
## ---
## Signif. codes:  0 '***' 0.001 '**' 0.01 '*' 0.05 '.' 0.1 ' ' 1
```

```
#test effect of polynomial
r1mod1f<-lm(NewMapComp_DeltaS~Generation*Treatment, data=R1ALL)
anova(r1mod1d,r1mod1f)#polynomial is useful
```

```
## Analysis of Variance Table
## 
## Model 1: NewMapComp_DeltaS ~ poly(Generation, 2) * Treatment
## Model 2: NewMapComp_DeltaS ~ Generation * Treatment
##   Res.Df   RSS Df Sum of Sq      F    Pr(>F)    
## 1    570 38996                                  
## 2    572 40623 -2   -1627.3 11.893 8.704e-06 ***
## ---
## Signif. codes:  0 '***' 0.001 '**' 0.01 '*' 0.05 '.' 0.1 ' ' 1
```

```
summary(r1mod1d)
```

```
## 
## Call:
## lm(formula = NewMapComp_DeltaS ~ poly(Generation, 2) * Treatment, 
##     data = R1ALL)
## 
## Residuals:
##     Min      1Q  Median      3Q     Max 
## -17.089  -5.617  -1.941   5.421  26.370 
## 
## Coefficients:
##                                        Estimate Std. Error t value Pr(>|t|)    
## (Intercept)                              8.4416     0.4874  17.320  < 2e-16 ***
## poly(Generation, 2)1                   -36.7096    11.6973  -3.138 0.001787 ** 
## poly(Generation, 2)2                    28.5475    11.6973   2.441 0.014970 *  
## TreatmentFurrowed                        8.1093     0.6893  11.765  < 2e-16 ***
## poly(Generation, 2)1:TreatmentFurrowed  58.8420    16.5425   3.557 0.000406 ***
## poly(Generation, 2)2:TreatmentFurrowed -77.9405    16.5425  -4.712 3.09e-06 ***
## ---
## Signif. codes:  0 '***' 0.001 '**' 0.01 '*' 0.05 '.' 0.1 ' ' 1
## 
## Residual standard error: 8.271 on 570 degrees of freedom
## Multiple R-squared:  0.2355, Adjusted R-squared:  0.2288 
## F-statistic: 35.13 on 5 and 570 DF,  p-value: < 2.2e-16
```

```
#plot

#make predictions
pred<-predict(r1mod1d, newdata = R1ALL, interval = 'confidence')
R1ALLp2<-cbind(R1ALL, pred)

#make plot
rep1colnew<-ggplot(R1ALLp2, aes(Generation, NewMapComp_DeltaS, col=Treatment,linetype=Treatment, fill=Treatment))+
  geom_point(alpha=0.3)+
  scale_colour_manual(values=c("black","grey50"))+
  scale_fill_manual(values=c("black","grey50"))+
  facet_grid(.~Treatment)+
  scale_x_continuous(breaks=c(0,5,10,15))+
  geom_ribbon(aes(ymin=lwr,ymax=upr), alpha=0.2, colour=NA)+
  geom_line(aes(Generation, fit), size=1)+
  guides(col=FALSE)+guides(linetype=FALSE)+guides(fill=FALSE)+
  ylab("Weighted average colour difference (Delta-S)")+
  my_theme+
  theme(axis.title = element_text(size = 16))  +
  theme(axis.text = element_text(size = 14)) +
  theme(strip.background = element_rect( fill="grey90"))+
  theme(strip.text.x = element_blank())
```

Luminance difference:

```
r1mod2<-lm(dL~poly(Generation,2)*Treatment, data=R1ALL)
par(mfrow=c(2,2))
plot(r1mod2)
```

```
hist(resid(r1mod2))
#best transform is square root
r1mod2<-lm(sqrt(dL)~poly(Generation,2)*Treatment, data=R1ALL)
par(mfrow=c(2,2))
```

```
plot(r1mod2)
```

```
hist(resid(r1mod2))

#model simplification
r1mod2b<-lm(sqrt(dL)~poly(Generation,2)+Treatment, data=R1ALL)
anova(r1mod2, r1mod2b)#significant interaction
```

```
## Analysis of Variance Table
## 
## Model 1: sqrt(dL) ~ poly(Generation, 2) * Treatment
## Model 2: sqrt(dL) ~ poly(Generation, 2) + Treatment
##   Res.Df    RSS Df Sum of Sq      F    Pr(>F)    
## 1    570 394.64                                  
## 2    572 420.95 -2   -26.305 18.997 1.031e-08 ***
## ---
## Signif. codes:  0 '***' 0.001 '**' 0.01 '*' 0.05 '.' 0.1 ' ' 1
```

```
#is polynomial useful?
r1mod2c<-lm(sqrt(dL)~Generation*Treatment, data=R1ALL)
anova(r1mod2,r1mod2c)#yes
```

```
## Analysis of Variance Table
## 
## Model 1: sqrt(dL) ~ poly(Generation, 2) * Treatment
## Model 2: sqrt(dL) ~ Generation * Treatment
##   Res.Df    RSS Df Sum of Sq      F  Pr(>F)   
## 1    570 394.64                               
## 2    572 401.45 -2   -6.8113 4.9189 0.00762 **
## ---
## Signif. codes:  0 '***' 0.001 '**' 0.01 '*' 0.05 '.' 0.1 ' ' 1
```

```
summary(r1mod2)
```

```
## 
## Call:
## lm(formula = sqrt(dL) ~ poly(Generation, 2) * Treatment, data = R1ALL)
## 
## Residuals:
##     Min      1Q  Median      3Q     Max 
## -2.1086 -0.6238 -0.1187  0.7316  1.8378 
## 
## Coefficients:
##                                        Estimate Std. Error t value Pr(>|t|)    
## (Intercept)                             1.91125    0.04903  38.981  < 2e-16 ***
## poly(Generation, 2)1                   -5.45995    1.17674  -4.640 4.33e-06 ***
## poly(Generation, 2)2                    2.36401    1.17674   2.009    0.045 *  
## TreatmentFurrowed                       0.40519    0.06934   5.844 8.60e-09 ***
## poly(Generation, 2)1:TreatmentFurrowed 10.24693    1.66416   6.157 1.40e-09 ***
## poly(Generation, 2)2:TreatmentFurrowed  0.47042    1.66416   0.283    0.778    
## ---
## Signif. codes:  0 '***' 0.001 '**' 0.01 '*' 0.05 '.' 0.1 ' ' 1
## 
## Residual standard error: 0.8321 on 570 degrees of freedom
## Multiple R-squared:  0.1259, Adjusted R-squared:  0.1182 
## F-statistic: 16.41 on 5 and 570 DF,  p-value: 3.792e-15
```

```
#make predictions
pred2<-predict(r1mod2, newdata = R1ALL, interval = 'confidence')
R1ALL$sqrtdL<-sqrt(R1ALL$dL)
R1ALLp2<-cbind(R1ALL, pred2)

rep1lum<-ggplot(R1ALLp2, aes(Generation, sqrtdL, col=Treatment,linetype=Treatment, fill=Treatment))+
  geom_point(alpha=0.3)+
  scale_colour_manual(values=c("black","grey50"))+
  scale_fill_manual(values=c("black","grey50"))+
  facet_grid(.~Treatment)+
  scale_x_continuous(breaks=c(0,5,10,15))+
  geom_ribbon(aes(ymin=lwr,ymax=upr), alpha=0.2, colour=NA)+
  geom_line(aes(Generation, fit), size=1)+
  guides(col=FALSE)+guides(linetype=FALSE)+guides(fill=FALSE)+
  ylab("Luminance difference (sqrt(Delta-L))")+
  my_theme+
  theme(axis.title = element_text(size = 16))  +
  theme(axis.text = element_text(size = 14)) +
  theme(strip.background = element_rect( fill="grey90"))+
  theme(strip.text.x = element_blank())
```

Pattern difference:

```
r1mod3<-lm(pattern_energy_difference~poly(Generation,2)*Treatment, data=R1ALL)
par(mfrow=c(2,2))
plot(r1mod3)
```

```
hist(resid(r1mod3))
#try a log transform
r1mod3<-lm(log(pattern_energy_difference)~poly(Generation,2)*Treatment, data=R1ALL)
par(mfrow=c(2,2))
```

```
plot(r1mod3)
```

```
hist(resid(r1mod3))#better

#model simplification
r1mod3b<-lm(log(pattern_energy_difference)~poly(Generation,2)+Treatment, data=R1ALL)
anova(r1mod3, r1mod3b)#significant interaction
```

```
## Analysis of Variance Table
## 
## Model 1: log(pattern_energy_difference) ~ poly(Generation, 2) * Treatment
## Model 2: log(pattern_energy_difference) ~ poly(Generation, 2) + Treatment
##   Res.Df    RSS Df Sum of Sq      F    Pr(>F)    
## 1    570 174.37                                  
## 2    572 179.91 -2   -5.5412 9.0571 0.0001342 ***
## ---
## Signif. codes:  0 '***' 0.001 '**' 0.01 '*' 0.05 '.' 0.1 ' ' 1
```

```
#is polynomial useful?
r1mod3c<-lm(log(pattern_energy_difference)~Generation*Treatment, data=R1ALL)
anova(r1mod3c, r1mod3)#polynomial not significant
```

```
## Analysis of Variance Table
## 
## Model 1: log(pattern_energy_difference) ~ Generation * Treatment
## Model 2: log(pattern_energy_difference) ~ poly(Generation, 2) * Treatment
##   Res.Df    RSS Df Sum of Sq      F Pr(>F)
## 1    572 175.20                           
## 2    570 174.37  2    0.8315 1.3591 0.2577
```

```
#further simplification
r1mod3d<-lm(log(pattern_energy_difference)~Generation+Treatment, data=R1ALL)
anova(r1mod3c, r1mod3d)#significant interaction
```

```
## Analysis of Variance Table
## 
## Model 1: log(pattern_energy_difference) ~ Generation * Treatment
## Model 2: log(pattern_energy_difference) ~ Generation + Treatment
##   Res.Df    RSS Df Sum of Sq      F    Pr(>F)    
## 1    572 175.20                                  
## 2    573 180.18 -1   -4.9869 16.282 6.201e-05 ***
## ---
## Signif. codes:  0 '***' 0.001 '**' 0.01 '*' 0.05 '.' 0.1 ' ' 1
```

```
summary(r1mod3c)
```

```
## 
## Call:
## lm(formula = log(pattern_energy_difference) ~ Generation * Treatment, 
##     data = R1ALL)
## 
## Residuals:
##     Min      1Q  Median      3Q     Max 
## -1.6556 -0.2709  0.0188  0.2938  1.5123 
## 
## Coefficients:
##                               Estimate Std. Error t value Pr(>|t|)    
## (Intercept)                  -2.826709   0.061345 -46.079  < 2e-16 ***
## Generation                    0.030328   0.009447   3.210   0.0014 ** 
## TreatmentFurrowed             0.026308   0.086754   0.303   0.7618    
## Generation:TreatmentFurrowed -0.053908   0.013360  -4.035  6.2e-05 ***
## ---
## Signif. codes:  0 '***' 0.001 '**' 0.01 '*' 0.05 '.' 0.1 ' ' 1
## 
## Residual standard error: 0.5534 on 572 degrees of freedom
## Multiple R-squared:  0.08165,    Adjusted R-squared:  0.07684 
## F-statistic: 16.95 on 3 and 572 DF,  p-value: 1.464e-10
```

```
#make predictions
pred3<-predict(r1mod3c, newdata = R1ALL, interval = 'confidence')
R1ALL$logpat<-log(R1ALL$pattern_energy_difference)
R1ALLp3<-cbind(R1ALL, pred3)

rep1pat<-ggplot(R1ALLp3, aes(Generation, logpat, col=Treatment,linetype=Treatment, fill=Treatment))+
  geom_point(alpha=0.3)+
  scale_colour_manual(values=c("black","grey50"))+
  scale_fill_manual(values=c("black","grey50"))+
  facet_grid(.~Treatment)+
  scale_x_continuous(breaks=c(0,5,10,15))+
  geom_ribbon(aes(ymin=lwr,ymax=upr), alpha=0.2, colour=NA)+
  geom_line(aes(Generation, fit), size=1)+
  guides(col=FALSE)+guides(linetype=FALSE)+guides(fill=FALSE)+
  ylab("Pattern difference (log(energy difference))")+
  my_theme+
  theme(axis.title = element_text(size = 16))  +
  theme(axis.text = element_text(size = 14)) +
  theme(strip.background = element_rect( fill="grey90"))+
  theme(strip.text.x = element_blank())
```

Edge disruption (GabRat):

```
r1mod4<-lm(GabRat~poly(Generation,2)*Treatment, data=R1ALL_GR)
par(mfrow=c(2,2))
plot(r1mod4)
```

```
hist(resid(r1mod4))
#log transform is better
r1mod4<-lm(log(GabRat)~poly(Generation,2)*Treatment, data=R1ALL_GR)
par(mfrow=c(2,2))
```

```
plot(r1mod4)
```

```
hist(resid(r1mod4))

#model simplification
r1mod4b<-lm(log(GabRat)~poly(Generation,2)+Treatment, data=R1ALL_GR)
anova(r1mod4, r1mod4b)#significant interaction
```

```
## Analysis of Variance Table
## 
## Model 1: log(GabRat) ~ poly(Generation, 2) * Treatment
## Model 2: log(GabRat) ~ poly(Generation, 2) + Treatment
##   Res.Df    RSS Df Sum of Sq      F   Pr(>F)   
## 1    570 82.339                                
## 2    572 83.712 -2   -1.3734 4.7538 0.008964 **
## ---
## Signif. codes:  0 '***' 0.001 '**' 0.01 '*' 0.05 '.' 0.1 ' ' 1
```

```
#test polynomial
r1mod4c<-lm(log(GabRat)~Generation*Treatment, data=R1ALL_GR)
anova(r1mod4,r1mod4c)#polynomial is useful
```

```
## Analysis of Variance Table
## 
## Model 1: log(GabRat) ~ poly(Generation, 2) * Treatment
## Model 2: log(GabRat) ~ Generation * Treatment
##   Res.Df    RSS Df Sum of Sq      F   Pr(>F)   
## 1    570 82.339                                
## 2    572 83.943 -2   -1.6044 5.5533 0.004087 **
## ---
## Signif. codes:  0 '***' 0.001 '**' 0.01 '*' 0.05 '.' 0.1 ' ' 1
```

```
summary(r1mod4)
```

```
## 
## Call:
## lm(formula = log(GabRat) ~ poly(Generation, 2) * Treatment, data = R1ALL_GR)
## 
## Residuals:
##      Min       1Q   Median       3Q      Max 
## -0.77341 -0.30811 -0.03684  0.28586  1.08510 
## 
## Coefficients:
##                                        Estimate Std. Error t value Pr(>|t|)    
## (Intercept)                            -2.21341    0.02240 -98.831  < 2e-16 ***
## poly(Generation, 2)1                    1.30818    0.53750   2.434   0.0152 *  
## poly(Generation, 2)2                    0.12210    0.53750   0.227   0.8204    
## TreatmentFurrowed                       0.25179    0.03167   7.950 1.01e-14 ***
## poly(Generation, 2)1:TreatmentFurrowed  1.35955    0.76014   1.789   0.0742 .  
## poly(Generation, 2)2:TreatmentFurrowed -1.90924    0.76014  -2.512   0.0123 *  
## ---
## Signif. codes:  0 '***' 0.001 '**' 0.01 '*' 0.05 '.' 0.1 ' ' 1
## 
## Residual standard error: 0.3801 on 570 degrees of freedom
## Multiple R-squared:  0.1554, Adjusted R-squared:  0.148 
## F-statistic: 20.97 on 5 and 570 DF,  p-value: < 2.2e-16
```

```
#make predictions
R1ALL_GR$loggr<-log(R1ALL_GR$GabRat)
r1mod4<-lm(loggr~poly(Generation,2)*Treatment, data=R1ALL_GR)

pred<-predict(r1mod4, newdata = R1ALL_GR, interval = 'confidence')
R1ALL_GRp<-cbind(R1ALL_GR, pred)

rep1gr<-ggplot(R1ALL_GRp, aes(Generation, loggr, col=Treatment,linetype=Treatment, fill=Treatment))+
  geom_point(alpha=0.3)+
  scale_colour_manual(values=c("black","grey50"))+
  scale_fill_manual(values=c("black","grey50"))+
  facet_grid(.~Treatment)+
  scale_x_continuous(breaks=c(0,5,10,15))+
  geom_ribbon(aes(ymin=lwr,ymax=upr), alpha=0.2, colour=NA)+
  geom_line(aes(Generation, fit), size=1)+
  guides(col=FALSE)+guides(linetype=FALSE)+guides(fill=FALSE)+
  ylab("Edge disruption (log(GabRat))")+
  my_theme+
  theme(axis.title = element_text(size = 16))  +
  theme(axis.text = element_text(size = 14)) +
  theme(strip.background = element_rect( fill="grey90"))+
  theme(strip.text.x = element_blank())
```

Table 1: Run 1 - mean colour and luminance contrast (prey-background) in first and final generations

| Generation | Treatment | dS | NewMapComp\_DeltaS | dL |
| --- | --- | --- | --- | --- |
| 0 | Smooth | 10.276131 | 12.852083 | 7.068491 |
| 0 | Furrowed | 10.276131 | 12.852083 | 7.068491 |
| 11 | Smooth | 9.319351 | 9.074474 | 3.473943 |
| 11 | Furrowed | 11.246019 | 15.120797 | 7.757217 |

###### Run 2

Colour difference between means:

```
r2mod1<-lm(dS~poly(Generation,2)*Treatment, data=R2ALL)
par(mfrow=c(2,2))
plot(r2mod1)
```

```
hist(resid(r2mod1))

#model simplification
r2mod1b<-lm(dS~poly(Generation,2)+Treatment, data=R2ALL)
anova(r2mod1,r2mod1b)#significant interaction
```

```
## Analysis of Variance Table
## 
## Model 1: dS ~ poly(Generation, 2) * Treatment
## Model 2: dS ~ poly(Generation, 2) + Treatment
##   Res.Df   RSS Df Sum of Sq      F    Pr(>F)    
## 1    810 12214                                  
## 2    812 12538 -2   -324.51 10.761 2.442e-05 ***
## ---
## Signif. codes:  0 '***' 0.001 '**' 0.01 '*' 0.05 '.' 0.1 ' ' 1
```

```
#is polynomial useful?
r2mod1c<-lm(dS~Generation*Treatment, data=R2ALL)
anova(r2mod1,r2mod1c)#yes
```

```
## Analysis of Variance Table
## 
## Model 1: dS ~ poly(Generation, 2) * Treatment
## Model 2: dS ~ Generation * Treatment
##   Res.Df   RSS Df Sum of Sq      F    Pr(>F)    
## 1    810 12214                                  
## 2    812 12509 -2   -295.77 9.8078 6.184e-05 ***
## ---
## Signif. codes:  0 '***' 0.001 '**' 0.01 '*' 0.05 '.' 0.1 ' ' 1
```

```
summary(r2mod1)
```

```
## 
## Call:
## lm(formula = dS ~ poly(Generation, 2) * Treatment, data = R2ALL)
## 
## Residuals:
##     Min      1Q  Median      3Q     Max 
## -8.7691 -3.2445  0.1997  2.8238  9.6717 
## 
## Coefficients:
##                                        Estimate Std. Error t value Pr(>|t|)    
## (Intercept)                             15.0707     0.1922  78.394  < 2e-16 ***
## poly(Generation, 2)1                    -3.2787     5.4915  -0.597 0.550643    
## poly(Generation, 2)2                     1.4522     5.4915   0.264 0.791504    
## TreatmentFurrowed                        0.8539     0.2719   3.141 0.001747 ** 
## poly(Generation, 2)1:TreatmentFurrowed -25.2185     7.7662  -3.247 0.001213 ** 
## poly(Generation, 2)2:TreatmentFurrowed -25.7305     7.7662  -3.313 0.000963 ***
## ---
## Signif. codes:  0 '***' 0.001 '**' 0.01 '*' 0.05 '.' 0.1 ' ' 1
## 
## Residual standard error: 3.883 on 810 degrees of freedom
## Multiple R-squared:  0.06549,    Adjusted R-squared:  0.05972 
## F-statistic: 11.35 on 5 and 810 DF,  p-value: 1.327e-10
```

```
#make predictions
pred<-predict(r2mod1, newdata = R2ALL, interval = 'confidence')
R2ALLp<-cbind(R2ALL, pred)

rep2col<-ggplot(R2ALLp, aes(Generation, dS, col=Treatment,linetype=Treatment, fill=Treatment))+
  geom_point(alpha=0.3)+
  scale_colour_manual(values=c("darkorange3","darkorange"))+
  scale_fill_manual(values=c("darkorange3","darkorange"))+
  facet_grid(.~Treatment)+
  scale_x_continuous(breaks=c(0,5,10,15))+
  geom_ribbon(aes(ymin=lwr,ymax=upr), alpha=0.2, colour=NA)+
  geom_line(aes(Generation, fit), size=1)+
  guides(col=FALSE)+guides(linetype=FALSE)+guides(fill=FALSE)+
  ylab("Colour difference between means (Delta-S)")+
  my_theme+
  theme(axis.title = element_text(size = 16))  +
  theme(axis.text = element_text(size = 14)) +
  theme(strip.background = element_rect( fill="grey90"))+
  theme(strip.text.x = element_blank())
```

Weighted average colour difference with new method:

```
r2mod2<-lm(NewMapComp_DeltaS~poly(Generation,2)*Treatment, data=R2ALL)
par(mfrow=c(2,2))
plot(r2mod2)
```

```
hist(resid(r2mod2))

#model simplification
r2mod2b<-lm(NewMapComp_DeltaS~poly(Generation,2)+Treatment, data=R2ALL)
anova(r2mod2,r2mod2b)#significant interaction
```

```
## Analysis of Variance Table
## 
## Model 1: NewMapComp_DeltaS ~ poly(Generation, 2) * Treatment
## Model 2: NewMapComp_DeltaS ~ poly(Generation, 2) + Treatment
##   Res.Df   RSS Df Sum of Sq      F    Pr(>F)    
## 1    810 71902                                  
## 2    812 74543 -2   -2640.8 14.875 4.527e-07 ***
## ---
## Signif. codes:  0 '***' 0.001 '**' 0.01 '*' 0.05 '.' 0.1 ' ' 1
```

```
#summary(r2mod2)

#is polynomial useful?
r2mod2c<-lm(NewMapComp_DeltaS~Generation*Treatment, data=R2ALL)
anova(r2mod2,r2mod2c)#no
```

```
## Analysis of Variance Table
## 
## Model 1: NewMapComp_DeltaS ~ poly(Generation, 2) * Treatment
## Model 2: NewMapComp_DeltaS ~ Generation * Treatment
##   Res.Df   RSS Df Sum of Sq      F Pr(>F)
## 1    810 71902                           
## 2    812 72131 -2   -228.65 1.2879 0.2764
```

```
#further simplification?
r2mod2d<-lm(NewMapComp_DeltaS~Generation+Treatment, data=R2ALL)
anova(r2mod2d,r2mod2c)#interaction still significant
```

```
## Analysis of Variance Table
## 
## Model 1: NewMapComp_DeltaS ~ Generation + Treatment
## Model 2: NewMapComp_DeltaS ~ Generation * Treatment
##   Res.Df   RSS Df Sum of Sq      F    Pr(>F)    
## 1    813 74624                                  
## 2    812 72131  1    2493.4 28.069 1.509e-07 ***
## ---
## Signif. codes:  0 '***' 0.001 '**' 0.01 '*' 0.05 '.' 0.1 ' ' 1
```

```
summary(r2mod2c)
```

```
## 
## Call:
## lm(formula = NewMapComp_DeltaS ~ Generation * Treatment, data = R2ALL)
## 
## Residuals:
##     Min      1Q  Median      3Q     Max 
## -22.343  -6.767  -0.517   5.418  31.782 
## 
## Coefficients:
##                              Estimate Std. Error t value Pr(>|t|)    
## (Intercept)                  13.33978    0.89349  14.930  < 2e-16 ***
## Generation                    0.65922    0.09525   6.921 9.09e-12 ***
## TreatmentFurrowed             0.81967    1.26358   0.649    0.517    
## Generation:TreatmentFurrowed -0.71363    0.13470  -5.298 1.51e-07 ***
## ---
## Signif. codes:  0 '***' 0.001 '**' 0.01 '*' 0.05 '.' 0.1 ' ' 1
## 
## Residual standard error: 9.425 on 812 degrees of freedom
## Multiple R-squared:  0.1127, Adjusted R-squared:  0.1094 
## F-statistic: 34.38 on 3 and 812 DF,  p-value: < 2.2e-16
```

```
#make predictions
pred<-predict(r2mod2c, newdata = R2ALL, interval = 'confidence')
R2ALLp<-cbind(R2ALL, pred)

rep2colnew<-ggplot(R2ALLp, aes(Generation, NewMapComp_DeltaS, col=Treatment,linetype=Treatment, fill=Treatment))+
  geom_point(alpha=0.3)+
  scale_colour_manual(values=c("darkorange3","darkorange"))+
  scale_fill_manual(values=c("darkorange3","darkorange"))+
  facet_grid(.~Treatment)+
  scale_x_continuous(breaks=c(0,5,10,15))+
  geom_ribbon(aes(ymin=lwr,ymax=upr), alpha=0.2, colour=NA)+
  geom_line(aes(Generation, fit), size=1)+
  guides(col=FALSE)+guides(linetype=FALSE)+guides(fill=FALSE)+
  ylab("Weighted average colour difference (Delta-S)")+
  my_theme+
  theme(axis.title = element_text(size = 16))  +
  theme(axis.text = element_text(size = 14)) +
  theme(strip.background = element_rect( fill="grey90"))+
  theme(strip.text.x = element_blank())
```

Luminance difference:

```
r2mod2<-lm(dL~poly(Generation,2)*Treatment, data=R2ALL)
par(mfrow=c(2,2))
plot(r2mod2)
```

```
hist(resid(r2mod2))

#model simplification
r2mod2b<-lm(dL~poly(Generation,2)+Treatment, data=R2ALL)
anova(r2mod2,r2mod2b)#significant interaction
```

```
## Analysis of Variance Table
## 
## Model 1: dL ~ poly(Generation, 2) * Treatment
## Model 2: dL ~ poly(Generation, 2) + Treatment
##   Res.Df    RSS Df Sum of Sq      F   Pr(>F)   
## 1    810 9499.1                                
## 2    812 9624.1 -2   -124.94 5.3269 0.005031 **
## ---
## Signif. codes:  0 '***' 0.001 '**' 0.01 '*' 0.05 '.' 0.1 ' ' 1
```

```
#is polynomial useful?
r2mod2c<-lm(dL~Generation*Treatment, data=R2ALL)
anova(r2mod2,r2mod2c)#no
```

```
## Analysis of Variance Table
## 
## Model 1: dL ~ poly(Generation, 2) * Treatment
## Model 2: dL ~ Generation * Treatment
##   Res.Df    RSS Df Sum of Sq      F Pr(>F)
## 1    810 9499.1                           
## 2    812 9546.2 -2   -47.035 2.0054 0.1353
```

```
#further simplification?
r2mod2d<-lm(dL~Generation+Treatment, data=R2ALL)
anova(r2mod2d,r2mod2c)#interaction still significant
```

```
## Analysis of Variance Table
## 
## Model 1: dL ~ Generation + Treatment
## Model 2: dL ~ Generation * Treatment
##   Res.Df    RSS Df Sum of Sq      F   Pr(>F)   
## 1    813 9626.9                                
## 2    812 9546.2  1    80.751 6.8687 0.008936 **
## ---
## Signif. codes:  0 '***' 0.001 '**' 0.01 '*' 0.05 '.' 0.1 ' ' 1
```

```
summary(r2mod2c)
```

```
## 
## Call:
## lm(formula = dL ~ Generation * Treatment, data = R2ALL)
## 
## Residuals:
##     Min      1Q  Median      3Q     Max 
## -7.6683 -2.8565 -0.0417  2.6607  9.0180 
## 
## Coefficients:
##                              Estimate Std. Error t value Pr(>|t|)    
## (Intercept)                   6.51000    0.32504  20.028  < 2e-16 ***
## Generation                   -0.16695    0.03465  -4.818 1.73e-06 ***
## TreatmentFurrowed             1.72376    0.45968   3.750 0.000189 ***
## Generation:TreatmentFurrowed -0.12843    0.04900  -2.621 0.008936 ** 
## ---
## Signif. codes:  0 '***' 0.001 '**' 0.01 '*' 0.05 '.' 0.1 ' ' 1
## 
## Residual standard error: 3.429 on 812 degrees of freedom
## Multiple R-squared:  0.1138, Adjusted R-squared:  0.1106 
## F-statistic: 34.77 on 3 and 812 DF,  p-value: < 2.2e-16
```

```
#make predictions
pred<-predict(r2mod2c, newdata = R2ALL, interval = 'confidence')
R2ALLp<-cbind(R2ALL, pred)

rep2lum<-ggplot(R2ALLp, aes(Generation, dL, col=Treatment,linetype=Treatment, fill=Treatment))+
  geom_point(alpha=0.3)+
  scale_colour_manual(values=c("darkorange3","darkorange"))+
  scale_fill_manual(values=c("darkorange3","darkorange"))+
  facet_grid(.~Treatment)+
  scale_x_continuous(breaks=c(0,5,10,15))+
  geom_ribbon(aes(ymin=lwr,ymax=upr), alpha=0.2, colour=NA)+
  geom_line(aes(Generation, fit), size=1)+
  guides(col=FALSE)+guides(linetype=FALSE)+guides(fill=FALSE)+
  ylab("Luminance difference (Delta-L)")+
  my_theme+
  theme(axis.title = element_text(size = 16))  +
  theme(axis.text = element_text(size = 14)) +
  theme(strip.background = element_rect( fill="grey90"))+
  theme(strip.text.x = element_blank())
```

Pattern difference:

```
r2mod3<-lm(pattern_energy_difference~poly(Generation,2)*Treatment, data=R2ALL)
par(mfrow=c(2,2))
plot(r2mod3)
```

```
hist(resid(r2mod3))
#log transform is better
r2mod3<-lm(log(pattern_energy_difference)~poly(Generation,2)*Treatment, data=R2ALL)
par(mfrow=c(2,2))
```

```
plot(r2mod3)
```

```
hist(resid(r2mod3))

#model simplication
r2mod3b<-lm(log(pattern_energy_difference)~poly(Generation,2)+Treatment, data=R2ALL)
anova(r2mod3, r2mod3b)#interaction not significant
```

```
## Analysis of Variance Table
## 
## Model 1: log(pattern_energy_difference) ~ poly(Generation, 2) * Treatment
## Model 2: log(pattern_energy_difference) ~ poly(Generation, 2) + Treatment
##   Res.Df    RSS Df Sum of Sq      F Pr(>F)
## 1    810 269.45                           
## 2    812 270.33 -2  -0.88333 1.3277 0.2657
```

```
r2mod3c<-lm(log(pattern_energy_difference)~poly(Generation,2), data=R2ALL)
anova(r2mod3c, r2mod3b)#no significant difference between backgrounds
```

```
## Analysis of Variance Table
## 
## Model 1: log(pattern_energy_difference) ~ poly(Generation, 2)
## Model 2: log(pattern_energy_difference) ~ poly(Generation, 2) + Treatment
##   Res.Df    RSS Df Sum of Sq      F Pr(>F)
## 1    813 270.84                           
## 2    812 270.33  1   0.50725 1.5236 0.2174
```

```
r2mod3d<-lm(log(pattern_energy_difference)~1, data=R2ALL)
anova(r2mod3c, r2mod3d) #no effect of generation either
```

```
## Analysis of Variance Table
## 
## Model 1: log(pattern_energy_difference) ~ poly(Generation, 2)
## Model 2: log(pattern_energy_difference) ~ 1
##   Res.Df    RSS Df Sum of Sq      F Pr(>F)
## 1    813 270.84                           
## 2    815 271.49 -2  -0.64934 0.9746 0.3778
```

```
#plot data only
rep2pat<-ggplot(R2ALL, aes(Generation, log(pattern_energy_difference), col=Treatment))+
  geom_point(alpha=0.4)+
  scale_colour_manual(values=c("darkorange3","darkorange"))+
  facet_grid(.~Treatment)+
  scale_x_continuous(breaks=c(0,5,10,15))+
  guides(col=FALSE)+guides(linetype=FALSE)+
  ylab("Pattern difference log(energy difference)")+
  my_theme+
  theme(axis.title = element_text(size = 16))  +
  theme(axis.text = element_text(size = 14)) +
  theme(strip.background = element_rect( fill="grey90"))+
  theme(strip.text.x = element_blank())
```

Edge disruption (GabRat):

```
r2mod4<-lm(GabRat~poly(Generation,2)*Treatment, data=R2ALL_GR)
par(mfrow=c(2,2))
plot(r2mod4)
```

```
hist(resid(r2mod4))
#log transform is better
r2mod4<-lm(log(GabRat)~poly(Generation,2)*Treatment, data=R2ALL_GR)
par(mfrow=c(2,2))
```

```
plot(r2mod4)
```

```
hist(resid(r2mod4))

#model simplification
r2mod4b<-lm(log(GabRat)~poly(Generation,2)+Treatment, data=R2ALL_GR)
anova(r2mod4,r2mod4b)#not significant
```

```
## Analysis of Variance Table
## 
## Model 1: log(GabRat) ~ poly(Generation, 2) * Treatment
## Model 2: log(GabRat) ~ poly(Generation, 2) + Treatment
##   Res.Df    RSS Df Sum of Sq      F Pr(>F)
## 1    810 94.158                           
## 2    812 94.617 -2  -0.45929 1.9755 0.1394
```

```
r2mod4c<-lm(log(GabRat)~poly(Generation,2), data=R2ALL_GR)
anova(r2mod4c,r2mod4b)#background type also not significant
```

```
## Analysis of Variance Table
## 
## Model 1: log(GabRat) ~ poly(Generation, 2)
## Model 2: log(GabRat) ~ poly(Generation, 2) + Treatment
##   Res.Df    RSS Df Sum of Sq      F Pr(>F)
## 1    813 94.679                           
## 2    812 94.617  1  0.061849 0.5308 0.4665
```

```
r2mod4d<-lm(log(GabRat)~1, data=R2ALL_GR)
anova(r2mod4d,r2mod4c)#significant effect of generation
```

```
## Analysis of Variance Table
## 
## Model 1: log(GabRat) ~ 1
## Model 2: log(GabRat) ~ poly(Generation, 2)
##   Res.Df     RSS Df Sum of Sq      F    Pr(>F)    
## 1    815 103.213                                  
## 2    813  94.679  2    8.5338 36.639 5.813e-16 ***
## ---
## Signif. codes:  0 '***' 0.001 '**' 0.01 '*' 0.05 '.' 0.1 ' ' 1
```

```
#is polynomial useful?
r2mod4e<-lm(log(GabRat)~Generation, data=R2ALL_GR)
anova(r2mod4c,r2mod4e)#no
```

```
## Analysis of Variance Table
## 
## Model 1: log(GabRat) ~ poly(Generation, 2)
## Model 2: log(GabRat) ~ Generation
##   Res.Df    RSS Df Sum of Sq      F Pr(>F)
## 1    813 94.679                           
## 2    814 94.807 -1  -0.12791 1.0983 0.2949
```

```
#check generation is still significant
anova(r2mod4e,r2mod4d)#yes
```

```
## Analysis of Variance Table
## 
## Model 1: log(GabRat) ~ Generation
## Model 2: log(GabRat) ~ 1
##   Res.Df     RSS Df Sum of Sq      F    Pr(>F)    
## 1    814  94.807                                  
## 2    815 103.213 -1   -8.4059 72.172 < 2.2e-16 ***
## ---
## Signif. codes:  0 '***' 0.001 '**' 0.01 '*' 0.05 '.' 0.1 ' ' 1
```

```
summary(r2mod4e)
```

```
## 
## Call:
## lm(formula = log(GabRat) ~ Generation, data = R2ALL_GR)
## 
## Residuals:
##      Min       1Q   Median       3Q      Max 
## -0.79299 -0.25532 -0.00302  0.24117  1.11863 
## 
## Coefficients:
##              Estimate Std. Error  t value Pr(>|t|)    
## (Intercept) -2.338660   0.022877 -102.228   <2e-16 ***
## Generation   0.020718   0.002439    8.495   <2e-16 ***
## ---
## Signif. codes:  0 '***' 0.001 '**' 0.01 '*' 0.05 '.' 0.1 ' ' 1
## 
## Residual standard error: 0.3413 on 814 degrees of freedom
## Multiple R-squared:  0.08144,    Adjusted R-squared:  0.08031 
## F-statistic: 72.17 on 1 and 814 DF,  p-value: < 2.2e-16
```

```
#make predictions
R2ALL_GR$loggr<-log(R2ALL_GR$GabRat)
pred<-predict(r2mod4e, newdata = R2ALL_GR, interval = 'confidence')
R2GRp<-cbind(R2ALL_GR, pred)

rep2gr<-ggplot(R2GRp, aes(Generation, loggr, col=Treatment,linetype=Treatment, fill=Treatment))+
  geom_point(alpha=0.3)+
  scale_colour_manual(values=c("darkorange3","darkorange"))+
  scale_fill_manual(values=c("darkorange3","darkorange"))+
  facet_grid(.~Treatment)+
  scale_x_continuous(breaks=c(0,5,10,15))+
  geom_ribbon(aes(ymin=lwr,ymax=upr), alpha=0.2, colour=NA)+
  geom_line(aes(Generation, fit), size=1)+
  guides(col=FALSE)+guides(linetype=FALSE)+guides(fill=FALSE)+
  ylab("Edge disruption (log(GabRat))")+
  my_theme+
  theme(axis.title = element_text(size = 16))  +
  theme(axis.text = element_text(size = 14)) +
  theme(strip.background = element_rect( fill="grey90"))+
  theme(strip.text.x = element_blank())
```

Table 2: Run 2 - mean colour and luminance contrast (prey-background) in first and final generations

| Generation | Treatment | dS | NewMapComp\_DeltaS | dL |
| --- | --- | --- | --- | --- |
| 0 | Smooth | 14.65251 | 14.18463 | 6.864370 |
| 0 | Furrowed | 14.65251 | 14.10947 | 6.864370 |
| 16 | Smooth | 14.36618 | 23.11811 | 4.041681 |
| 16 | Furrowed | 13.40555 | 10.07045 | 3.881180 |

###### Plots for field trials

Figure 4: Camouflage metrics across generations, in field trials

#### Screen-based replicates

##### Load and prepare data

Output from CamoEvo, and additional pattern difference and colour distance (weighted average) analyses

Key variables in main dataset are: Bgd (Background - furrowed (Fr) or smooth (Sm)), ColSpace (colour space - narrow or full), Folder/Participant ID (identify individual populations/participants), Generation (integer) + all CamoEvo metrics and gene data.

```
CWSCREEN<-read.csv("CamoWild_ScreenData.csv", header=TRUE)
CWSCREEN_PATTERN<-read.csv("CamoWild_ScreenData_Pattern.csv", header=TRUE)
CWSCREEN_COLDIST<-read.csv("CamoWild_ScreenData_ColDist.csv", header=TRUE)

#colour difference between means - background to target (euclidean distance) 
CWSCREEN$eucdisTB<-sqrt((CWSCREEN$Target_L_Mean-CWSCREEN$BG_L_Mean)^2+(CWSCREEN$Target_A_Mean-CWSCREEN$BG_A_Mean)^2+
                          (CWSCREEN$Target_B_Mean-CWSCREEN$BG_B_Mean)^2)
#luminance difference (difference in L values)
CWSCREEN$LdiffTB<-abs(CWSCREEN$Target_L_Mean-CWSCREEN$BG_L_Mean)

#separate full and narrow colour spaces for subsequent analyses
CWSCREEN_FULL<-CWSCREEN%>%
  filter(ColSpace=="Full")
CWSCREEN_NARROW<-CWSCREEN%>%
  filter(ColSpace=="Narrow")
```

How do metrics of camouflage efficacy change across generations, in populations on different backgrounds & using different colour spaces?

##### Camouflage metrics: luminance difference

```
#start with Generation (2nd order polynomial) and Bgd, interacting, 
#+ random effect of Folder(Population) with random slope for generation

#full colour space
fl1<-lmer(LdiffTB~poly(Generation,2)*Bgd+(Generation|Folder), data=CWSCREEN_FULL)
#summary(fl1)

#test model assumptions
plot(fl1)
```

```
hist(resid(fl1))#not bad, not improved by transforms
```

```
#simplify model
fl1b<-lmer(LdiffTB~poly(Generation,2)+Bgd+(Generation|Folder), data=CWSCREEN_FULL) 
anova(fl1,fl1b)#can remove interaction
```

```
## Data: CWSCREEN_FULL
## Models:
## fl1b: LdiffTB ~ poly(Generation, 2) + Bgd + (Generation | Folder)
## fl1: LdiffTB ~ poly(Generation, 2) * Bgd + (Generation | Folder)
##      Df   AIC   BIC  logLik deviance  Chisq Chi Df Pr(>Chisq)
## fl1b  8 19303 19349 -9643.4    19287                         
## fl1  10 19307 19365 -9643.4    19287 0.1461      2     0.9295
```

```
#summary(fl1b)
fl1c<-lmer(LdiffTB~poly(Generation,2)+(Generation|Folder), data=CWSCREEN_FULL) 
anova(fl1c,fl1b)#remove bgd
```

```
## Data: CWSCREEN_FULL
## Models:
## fl1c: LdiffTB ~ poly(Generation, 2) + (Generation | Folder)
## fl1b: LdiffTB ~ poly(Generation, 2) + Bgd + (Generation | Folder)
##      Df   AIC   BIC  logLik deviance  Chisq Chi Df Pr(>Chisq)
## fl1c  7 19302 19342 -9643.8    19288                         
## fl1b  8 19303 19349 -9643.4    19287 0.7093      1     0.3997
```

```
fl1d<-lmer(LdiffTB~(Generation|Folder), data=CWSCREEN_FULL) 
anova(fl1c,fl1d)#remove generation
```

```
## Data: CWSCREEN_FULL
## Models:
## fl1d: LdiffTB ~ (Generation | Folder)
## fl1c: LdiffTB ~ poly(Generation, 2) + (Generation | Folder)
##      Df   AIC   BIC  logLik deviance  Chisq Chi Df Pr(>Chisq)
## fl1d  5 19298 19327 -9644.2    19288                         
## fl1c  7 19302 19342 -9643.8    19288 0.8012      2     0.6699
```

```
#test significance of random effects
rand(fl1d)
```

```
## ANOVA-like table for random-effects: Single term deletions
## 
## Model:
## LdiffTB ~ (Generation | Folder)
##                                     npar  logLik   AIC    LRT Df Pr(>Chisq)    
## <none>                                 5 -9642.9 19296                         
## Generation in (Generation | Folder)    3 -9714.9 19436 143.98  2  < 2.2e-16 ***
## ---
## Signif. codes:  0 '***' 0.001 '**' 0.01 '*' 0.05 '.' 0.1 ' ' 1
```

```
#without polynomial?
fl1_2<-lmer(LdiffTB~Generation*Bgd+(Generation|Folder), data=CWSCREEN_FULL) 
fl1_2b<-lmer(LdiffTB~Generation+Bgd+(Generation|Folder), data=CWSCREEN_FULL) 
anova(fl1_2b, fl1_2)#remove interaction
```

```
## Data: CWSCREEN_FULL
## Models:
## fl1_2b: LdiffTB ~ Generation + Bgd + (Generation | Folder)
## fl1_2: LdiffTB ~ Generation * Bgd + (Generation | Folder)
##        Df   AIC   BIC  logLik deviance  Chisq Chi Df Pr(>Chisq)
## fl1_2b  7 19302 19342 -9643.8    19288                         
## fl1_2   8 19304 19350 -9643.8    19288 0.0236      1      0.878
```

```
fl1_2c<-lmer(LdiffTB~Generation+(Generation|Folder), data=CWSCREEN_FULL) 
anova(fl1_2b, fl1_2c)#remove bgd
```

```
## Data: CWSCREEN_FULL
## Models:
## fl1_2c: LdiffTB ~ Generation + (Generation | Folder)
## fl1_2b: LdiffTB ~ Generation + Bgd + (Generation | Folder)
##        Df   AIC   BIC  logLik deviance  Chisq Chi Df Pr(>Chisq)
## fl1_2c  6 19300 19335 -9644.2    19288                         
## fl1_2b  7 19302 19342 -9643.8    19288 0.7093      1     0.3997
```

```
fl1_2d<-lmer(LdiffTB~(Generation|Folder), data=CWSCREEN_FULL) 
anova(fl1_2d, fl1_2c)#remove generation - back to earlier model
```

```
## Data: CWSCREEN_FULL
## Models:
## fl1_2d: LdiffTB ~ (Generation | Folder)
## fl1_2c: LdiffTB ~ Generation + (Generation | Folder)
##        Df   AIC   BIC  logLik deviance Chisq Chi Df Pr(>Chisq)
## fl1_2d  5 19298 19327 -9644.2    19288                        
## fl1_2c  6 19300 19335 -9644.2    19288 0.094      1     0.7591
```

```
#model without random slope
fl2<-lmer(LdiffTB~poly(Generation,2)*Bgd+(1|Folder), data=CWSCREEN_FULL)
fl2b<-lmer(LdiffTB~poly(Generation,2)+Bgd+(1|Folder), data=CWSCREEN_FULL)
anova(fl2, fl2b)#remove Bgd:Generation interaction
```

```
## Data: CWSCREEN_FULL
## Models:
## fl2b: LdiffTB ~ poly(Generation, 2) + Bgd + (1 | Folder)
## fl2: LdiffTB ~ poly(Generation, 2) * Bgd + (1 | Folder)
##      Df   AIC   BIC  logLik deviance  Chisq Chi Df Pr(>Chisq)
## fl2b  6 19443 19478 -9715.4    19431                         
## fl2   8 19446 19493 -9715.1    19430 0.7204      2     0.6975
```

```
fl2c<-lmer(LdiffTB~poly(Generation,2)+(1|Folder), data=CWSCREEN_FULL)
anova(fl2c, fl2b)
```

```
## Data: CWSCREEN_FULL
## Models:
## fl2c: LdiffTB ~ poly(Generation, 2) + (1 | Folder)
## fl2b: LdiffTB ~ poly(Generation, 2) + Bgd + (1 | Folder)
##      Df   AIC   BIC  logLik deviance  Chisq Chi Df Pr(>Chisq)
## fl2c  5 19441 19470 -9715.5    19431                         
## fl2b  6 19443 19478 -9715.4    19431 0.0507      1     0.8218
```

```
fl2d<-lmer(LdiffTB~(1|Folder), data=CWSCREEN_FULL)
anova(fl2c, fl2d)#remove - no effects
```

```
## Data: CWSCREEN_FULL
## Models:
## fl2d: LdiffTB ~ (1 | Folder)
## fl2c: LdiffTB ~ poly(Generation, 2) + (1 | Folder)
##      Df   AIC   BIC  logLik deviance  Chisq Chi Df Pr(>Chisq)
## fl2d  3 19440 19457 -9717.0    19434                         
## fl2c  5 19441 19470 -9715.5    19431 3.0999      2     0.2123
```

```
#without polynomial?
fl2_2<-lmer(LdiffTB~Generation*Bgd+(1|Folder), data=CWSCREEN_FULL)
fl2b_2<-lmer(LdiffTB~Generation+Bgd+(1|Folder), data=CWSCREEN_FULL)
anova(fl2_2, fl2b_2)#remove Bgd:Generation interaction
```

```
## Data: CWSCREEN_FULL
## Models:
## fl2b_2: LdiffTB ~ Generation + Bgd + (1 | Folder)
## fl2_2: LdiffTB ~ Generation * Bgd + (1 | Folder)
##        Df   AIC   BIC  logLik deviance  Chisq Chi Df Pr(>Chisq)
## fl2b_2  5 19442 19471 -9715.8    19432                         
## fl2_2   6 19443 19478 -9715.5    19431 0.6051      1     0.4366
```

```
fl2c_2<-lmer(LdiffTB~Generation+(1|Folder), data=CWSCREEN_FULL)
anova(fl2c_2, fl2b_2)
```

```
## Data: CWSCREEN_FULL
## Models:
## fl2c_2: LdiffTB ~ Generation + (1 | Folder)
## fl2b_2: LdiffTB ~ Generation + Bgd + (1 | Folder)
##        Df   AIC   BIC  logLik deviance  Chisq Chi Df Pr(>Chisq)
## fl2c_2  4 19440 19463 -9715.8    19432                         
## fl2b_2  5 19442 19471 -9715.8    19432 0.0507      1     0.8218
```

```
fl2d_2<-lmer(LdiffTB~(1|Folder), data=CWSCREEN_FULL)
anova(fl2c_2, fl2d_2)#remove all
```

```
## Data: CWSCREEN_FULL
## Models:
## fl2d_2: LdiffTB ~ (1 | Folder)
## fl2c_2: LdiffTB ~ Generation + (1 | Folder)
##        Df   AIC   BIC  logLik deviance  Chisq Chi Df Pr(>Chisq)
## fl2d_2  3 19440 19457 -9717.0    19434                         
## fl2c_2  4 19440 19463 -9715.8    19432 2.4358      1     0.1186
```

```
#compare final models with and w/o random slope:
AIC(fl1d, fl2d)#lower AIC with random slope
```

```
##      df      AIC
## fl1d  5 19295.84
## fl2d  3 19435.83
```

```
#narrow colour space

nl1<-lmer(LdiffTB~poly(Generation,2)*Bgd+(Generation|Folder), data=CWSCREEN_NARROW)
#test model assumptions
plot(nl1)
```

```
hist(resid(nl1))
```

```
nl1<-lmer(sqrt(LdiffTB)~poly(Generation,2)*Bgd+(Generation|Folder), data=CWSCREEN_NARROW)
#summary(nl1)
plot(nl1)
```

```
hist(resid(nl1))# better with transform
```

```
#simplify model
nl1b<-lmer(sqrt(LdiffTB)~poly(Generation,2)+Bgd+(Generation|Folder), data=CWSCREEN_NARROW) 
anova(nl1,nl1b)#can remove interaction
```

```
## Data: CWSCREEN_NARROW
## Models:
## nl1b: sqrt(LdiffTB) ~ poly(Generation, 2) + Bgd + (Generation | Folder)
## nl1: sqrt(LdiffTB) ~ poly(Generation, 2) * Bgd + (Generation | Folder)
##      Df    AIC    BIC  logLik deviance  Chisq Chi Df Pr(>Chisq)  
## nl1b  8 6934.4 6980.8 -3459.2   6918.4                           
## nl1  10 6933.4 6991.5 -3456.7   6913.4 4.9601      2    0.08374 .
## ---
## Signif. codes:  0 '***' 0.001 '**' 0.01 '*' 0.05 '.' 0.1 ' ' 1
```

```
nl1c<-lmer(sqrt(LdiffTB)~poly(Generation,2)+(Generation|Folder), data=CWSCREEN_NARROW) 
anova(nl1c,nl1b)#can remove bgd
```

```
## Data: CWSCREEN_NARROW
## Models:
## nl1c: sqrt(LdiffTB) ~ poly(Generation, 2) + (Generation | Folder)
## nl1b: sqrt(LdiffTB) ~ poly(Generation, 2) + Bgd + (Generation | Folder)
##      Df    AIC    BIC  logLik deviance  Chisq Chi Df Pr(>Chisq)
## nl1c  7 6935.0 6975.6 -3460.5   6921.0                         
## nl1b  8 6934.4 6980.8 -3459.2   6918.4 2.6148      1     0.1059
```

```
nl1d<-lmer(sqrt(LdiffTB)~(Generation|Folder), data=CWSCREEN_NARROW) 
anova(nl1c,nl1d)#sig effect of generation
```

```
## Data: CWSCREEN_NARROW
## Models:
## nl1d: sqrt(LdiffTB) ~ (Generation | Folder)
## nl1c: sqrt(LdiffTB) ~ poly(Generation, 2) + (Generation | Folder)
##      Df    AIC    BIC  logLik deviance  Chisq Chi Df Pr(>Chisq)    
## nl1d  5 6950.3 6979.3 -3470.2   6940.3                             
## nl1c  7 6935.0 6975.6 -3460.5   6921.0 19.315      2  6.394e-05 ***
## ---
## Signif. codes:  0 '***' 0.001 '**' 0.01 '*' 0.05 '.' 0.1 ' ' 1
```

```
#test significance of random effects
rand(nl1c)#random effects not significant
```

```
## ANOVA-like table for random-effects: Single term deletions
## 
## Model:
## sqrt(LdiffTB) ~ poly(Generation, 2) + (Generation | Folder)
##                                     npar  logLik    AIC     LRT Df Pr(>Chisq)
## <none>                                 7 -3460.5 6935.0                      
## Generation in (Generation | Folder)    5 -3461.0 6931.9 0.95762  2     0.6195
```

```
#is the polynomial useful?
nl1c_2<-lmer(sqrt(LdiffTB)~Generation+(Generation|Folder), data=CWSCREEN_NARROW) 
anova(nl1c_2, nl1c)#polynomial not needed
```

```
## Data: CWSCREEN_NARROW
## Models:
## nl1c_2: sqrt(LdiffTB) ~ Generation + (Generation | Folder)
## nl1c: sqrt(LdiffTB) ~ poly(Generation, 2) + (Generation | Folder)
##        Df    AIC    BIC  logLik deviance  Chisq Chi Df Pr(>Chisq)
## nl1c_2  6 6935.5 6970.4 -3461.8   6923.5                         
## nl1c    7 6935.0 6975.6 -3460.5   6921.0 2.5268      1     0.1119
```

```
nl1c_2b<-lmer(sqrt(LdiffTB)~(Generation|Folder), data=CWSCREEN_NARROW) 
anova(nl1c_2b, nl1c_2)#generation still significant
```

```
## Data: CWSCREEN_NARROW
## Models:
## nl1c_2b: sqrt(LdiffTB) ~ (Generation | Folder)
## nl1c_2: sqrt(LdiffTB) ~ Generation + (Generation | Folder)
##         Df    AIC    BIC  logLik deviance  Chisq Chi Df Pr(>Chisq)    
## nl1c_2b  5 6950.3 6979.3 -3470.2   6940.3                             
## nl1c_2   6 6935.5 6970.4 -3461.8   6923.5 16.788      1  4.179e-05 ***
## ---
## Signif. codes:  0 '***' 0.001 '**' 0.01 '*' 0.05 '.' 0.1 ' ' 1
```

```
rand(nl1c_2)
```

```
## ANOVA-like table for random-effects: Single term deletions
## 
## Model:
## sqrt(LdiffTB) ~ Generation + (Generation | Folder)
##                                     npar  logLik    AIC     LRT Df Pr(>Chisq)
## <none>                                 6 -3468.2 6948.3                      
## Generation in (Generation | Folder)    4 -3468.6 6945.3 0.95601  2       0.62
```

```
#model without random slope
nl2<-lmer(sqrt(LdiffTB)~poly(Generation,2)*Bgd+(1|Folder), data=CWSCREEN_NARROW)
nl2b<-lmer(sqrt(LdiffTB)~poly(Generation,2)+Bgd+(1|Folder), data=CWSCREEN_NARROW)
anova(nl2, nl2b)#remove Bgd:Generation interaction
```

```
## Data: CWSCREEN_NARROW
## Models:
## nl2b: sqrt(LdiffTB) ~ poly(Generation, 2) + Bgd + (1 | Folder)
## nl2: sqrt(LdiffTB) ~ poly(Generation, 2) * Bgd + (1 | Folder)
##      Df    AIC    BIC  logLik deviance  Chisq Chi Df Pr(>Chisq)  
## nl2b  6 6931.5 6966.3 -3459.7   6919.5                           
## nl2   8 6930.5 6976.9 -3457.3   6914.5 4.9576      2    0.08384 .
## ---
## Signif. codes:  0 '***' 0.001 '**' 0.01 '*' 0.05 '.' 0.1 ' ' 1
```

```
nl2c<-lmer(sqrt(LdiffTB)~poly(Generation,2)+(1|Folder), data=CWSCREEN_NARROW)
anova(nl2c, nl2b)
```

```
## Data: CWSCREEN_NARROW
## Models:
## nl2c: sqrt(LdiffTB) ~ poly(Generation, 2) + (1 | Folder)
## nl2b: sqrt(LdiffTB) ~ poly(Generation, 2) + Bgd + (1 | Folder)
##      Df    AIC    BIC  logLik deviance  Chisq Chi Df Pr(>Chisq)
## nl2c  5 6931.7 6960.8 -3460.9   6921.7                         
## nl2b  6 6931.5 6966.3 -3459.7   6919.5 2.2702      1     0.1319
```

```
nl2d<-lmer(sqrt(LdiffTB)~(1|Folder), data=CWSCREEN_NARROW)
anova(nl2c, nl2d)#generation still significant
```

```
## Data: CWSCREEN_NARROW
## Models:
## nl2d: sqrt(LdiffTB) ~ (1 | Folder)
## nl2c: sqrt(LdiffTB) ~ poly(Generation, 2) + (1 | Folder)
##      Df    AIC    BIC  logLik deviance  Chisq Chi Df Pr(>Chisq)    
## nl2d  3 7044.3 7061.7 -3519.2   7038.3                             
## nl2c  5 6931.7 6960.8 -3460.9   6921.7 116.59      2  < 2.2e-16 ***
## ---
## Signif. codes:  0 '***' 0.001 '**' 0.01 '*' 0.05 '.' 0.1 ' ' 1
```

```
rand(nl2c)#random effects significant
```

```
## ANOVA-like table for random-effects: Single term deletions
## 
## Model:
## sqrt(LdiffTB) ~ poly(Generation, 2) + (1 | Folder)
##              npar  logLik    AIC    LRT Df Pr(>Chisq)    
## <none>          5 -3461.0 6931.9                         
## (1 | Folder)    4 -3475.8 6959.6 29.613  1  5.275e-08 ***
## ---
## Signif. codes:  0 '***' 0.001 '**' 0.01 '*' 0.05 '.' 0.1 ' ' 1
```

```
#is the polynomial useful?
nl2c_2<-lmer(sqrt(LdiffTB)~Generation+(1|Folder), data=CWSCREEN_NARROW)
anova(nl2c_2,nl2c)#polynomial not needed
```

```
## Data: CWSCREEN_NARROW
## Models:
## nl2c_2: sqrt(LdiffTB) ~ Generation + (1 | Folder)
## nl2c: sqrt(LdiffTB) ~ poly(Generation, 2) + (1 | Folder)
##        Df    AIC    BIC  logLik deviance  Chisq Chi Df Pr(>Chisq)
## nl2c_2  4 6932.3 6955.5 -3462.1   6924.3                         
## nl2c    5 6931.7 6960.8 -3460.9   6921.7 2.5252      1      0.112
```

```
nl2d_2<-lmer(sqrt(LdiffTB)~(1|Folder), data=CWSCREEN_NARROW)
anova(nl2d_2, nl2c_2)#generation still significant
```

```
## Data: CWSCREEN_NARROW
## Models:
## nl2d_2: sqrt(LdiffTB) ~ (1 | Folder)
## nl2c_2: sqrt(LdiffTB) ~ Generation + (1 | Folder)
##        Df    AIC    BIC  logLik deviance  Chisq Chi Df Pr(>Chisq)    
## nl2d_2  3 7044.3 7061.7 -3519.2   7038.3                             
## nl2c_2  4 6932.3 6955.5 -3462.1   6924.3 114.07      1  < 2.2e-16 ***
## ---
## Signif. codes:  0 '***' 0.001 '**' 0.01 '*' 0.05 '.' 0.1 ' ' 1
```

```
#compare final models with and w/o random slope:
AIC(nl1c_2, nl2c_2)#lower AIC without random slope
```

```
##        df      AIC
## nl1c_2  6 6948.317
## nl2c_2  4 6945.273
```

##### Camouflage metrics: colour difference

```
#start with Generation (2nd order polynomial) and Bgd, interacting 
#+ random effect of Folder(population) with random slope for generation

#full colour space
fc1<-lmer(eucdisTB~poly(Generation,2)*Bgd+(Generation|Folder), data=CWSCREEN_FULL)
#summary(fc1)

#test model assumptions
plot(fc1)
```

```
hist(resid(fc1))#ok
```

```
#simplify model
fc1b<-lmer(eucdisTB~poly(Generation,2)+Bgd+(Generation|Folder), data=CWSCREEN_FULL) 
anova(fc1,fc1b)#can remove interaction
```

```
## Data: CWSCREEN_FULL
## Models:
## fc1b: eucdisTB ~ poly(Generation, 2) + Bgd + (Generation | Folder)
## fc1: eucdisTB ~ poly(Generation, 2) * Bgd + (Generation | Folder)
##      Df   AIC   BIC  logLik deviance  Chisq Chi Df Pr(>Chisq)
## fc1b  8 19395 19442 -9689.6    19379                         
## fc1  10 19399 19457 -9689.5    19379 0.1434      2     0.9308
```

```
fc1c<-lmer(eucdisTB~poly(Generation,2)+(Generation|Folder), data=CWSCREEN_FULL) 
anova(fc1c,fc1b)#remove bgd
```

```
## Data: CWSCREEN_FULL
## Models:
## fc1c: eucdisTB ~ poly(Generation, 2) + (Generation | Folder)
## fc1b: eucdisTB ~ poly(Generation, 2) + Bgd + (Generation | Folder)
##      Df   AIC   BIC  logLik deviance  Chisq Chi Df Pr(>Chisq)
## fc1c  7 19393 19434 -9689.7    19379                         
## fc1b  8 19395 19442 -9689.6    19379 0.2504      1     0.6168
```

```
fc1d<-lmer(eucdisTB~(Generation|Folder), data=CWSCREEN_FULL) 
anova(fc1c,fc1d)#can't remove generation
```

```
## Data: CWSCREEN_FULL
## Models:
## fc1d: eucdisTB ~ (Generation | Folder)
## fc1c: eucdisTB ~ poly(Generation, 2) + (Generation | Folder)
##      Df   AIC   BIC  logLik deviance  Chisq Chi Df Pr(>Chisq)  
## fc1d  5 19397 19426 -9693.3    19387                           
## fc1c  7 19393 19434 -9689.7    19379 7.2559      2    0.02657 *
## ---
## Signif. codes:  0 '***' 0.001 '**' 0.01 '*' 0.05 '.' 0.1 ' ' 1
```

```
#summary(fc1c)

#test significance of random effects
rand(fc1d)
```

```
## ANOVA-like table for random-effects: Single term deletions
## 
## Model:
## eucdisTB ~ (Generation | Folder)
##                                     npar  logLik   AIC    LRT Df Pr(>Chisq)    
## <none>                                 5 -9692.1 19394                         
## Generation in (Generation | Folder)    3 -9815.6 19637 247.19  2  < 2.2e-16 ***
## ---
## Signif. codes:  0 '***' 0.001 '**' 0.01 '*' 0.05 '.' 0.1 ' ' 1
```

```
#is the polynomial useful?
fc1c_2<-lmer(eucdisTB~Generation+(Generation|Folder), data=CWSCREEN_FULL) 
anova(fc1c,fc1c_2)#polynomial useful
```

```
## Data: CWSCREEN_FULL
## Models:
## fc1c_2: eucdisTB ~ Generation + (Generation | Folder)
## fc1c: eucdisTB ~ poly(Generation, 2) + (Generation | Folder)
##        Df   AIC   BIC  logLik deviance Chisq Chi Df Pr(>Chisq)  
## fc1c_2  6 19397 19432 -9692.6    19385                          
## fc1c    7 19393 19434 -9689.7    19379 5.854      1    0.01554 *
## ---
## Signif. codes:  0 '***' 0.001 '**' 0.01 '*' 0.05 '.' 0.1 ' ' 1
```

```
#model without random slope
fc2<-lmer(eucdisTB~poly(Generation,2)*Bgd+(1|Folder), data=CWSCREEN_FULL)
fc2b<-lmer(eucdisTB~poly(Generation,2)+Bgd+(1|Folder), data=CWSCREEN_FULL)
anova(fc2, fc2b)#remove Bgd:Generation interaction
```

```
## Data: CWSCREEN_FULL
## Models:
## fc2b: eucdisTB ~ poly(Generation, 2) + Bgd + (1 | Folder)
## fc2: eucdisTB ~ poly(Generation, 2) * Bgd + (1 | Folder)
##      Df   AIC   BIC  logLik deviance  Chisq Chi Df Pr(>Chisq)
## fc2b  6 19589 19623 -9788.2    19577                         
## fc2   8 19592 19638 -9787.8    19576 0.9161      2     0.6325
```

```
fc2c<-lmer(eucdisTB~poly(Generation,2)+(1|Folder), data=CWSCREEN_FULL)
anova(fc2c, fc2b)#can remove background
```

```
## Data: CWSCREEN_FULL
## Models:
## fc2c: eucdisTB ~ poly(Generation, 2) + (1 | Folder)
## fc2b: eucdisTB ~ poly(Generation, 2) + Bgd + (1 | Folder)
##      Df   AIC   BIC  logLik deviance  Chisq Chi Df Pr(>Chisq)
## fc2c  5 19587 19616 -9788.3    19577                         
## fc2b  6 19589 19623 -9788.2    19577 0.1277      1     0.7208
```

```
fc2d<-lmer(eucdisTB~(1|Folder), data=CWSCREEN_FULL)
anova(fc2c, fc2d)#can't remove generation
```

```
## Data: CWSCREEN_FULL
## Models:
## fc2d: eucdisTB ~ (1 | Folder)
## fc2c: eucdisTB ~ poly(Generation, 2) + (1 | Folder)
##      Df   AIC   BIC  logLik deviance  Chisq Chi Df Pr(>Chisq)    
## fc2d  3 19642 19659 -9817.7    19636                             
## fc2c  5 19587 19616 -9788.3    19577 58.867      2  1.649e-13 ***
## ---
## Signif. codes:  0 '***' 0.001 '**' 0.01 '*' 0.05 '.' 0.1 ' ' 1
```

```
#is the polynomial useful?
fc2c_2<-lmer(eucdisTB~Generation+(1|Folder), data=CWSCREEN_FULL)
anova(fc2c_2, fc2c)#polynomial signficant
```

```
## Data: CWSCREEN_FULL
## Models:
## fc2c_2: eucdisTB ~ Generation + (1 | Folder)
## fc2c: eucdisTB ~ poly(Generation, 2) + (1 | Folder)
##        Df   AIC   BIC  logLik deviance  Chisq Chi Df Pr(>Chisq)  
## fc2c_2  4 19590 19613 -9791.0    19582                           
## fc2c    5 19587 19616 -9788.3    19577 5.3741      1    0.02044 *
## ---
## Signif. codes:  0 '***' 0.001 '**' 0.01 '*' 0.05 '.' 0.1 ' ' 1
```

```
#compare final models with and w/o random slope:
AIC(fc1c, fc2c)#lower AIC with random slope
```

```
##      df      AIC
## fc1c  7 19373.34
## fc2c  5 19568.45
```

```
#narrow colour space
nc1<-lmer(eucdisTB~poly(Generation,2)*Bgd+(Generation|Folder), data=CWSCREEN_NARROW)
#summary(nc1)

#test model assumptions
plot(nc1)
```

```
hist(resid(nc1))
```

```
nc1<-lmer(sqrt(eucdisTB)~poly(Generation,2)*Bgd+(Generation|Folder), data=CWSCREEN_NARROW)
#summary(nc1)
plot(nc1)
```

```
hist(resid(nc1))#better with sqrt transform
```

```
#simplify model
nc1b<-lmer(sqrt(eucdisTB)~poly(Generation,2)+Bgd+(Generation|Folder), data=CWSCREEN_NARROW) 
anova(nc1,nc1b)#can remove interaction
```

```
## Data: CWSCREEN_NARROW
## Models:
## nc1b: sqrt(eucdisTB) ~ poly(Generation, 2) + Bgd + (Generation | Folder)
## nc1: sqrt(eucdisTB) ~ poly(Generation, 2) * Bgd + (Generation | Folder)
##      Df    AIC  BIC  logLik deviance  Chisq Chi Df Pr(>Chisq)
## nc1b  8 6106.6 6153 -3045.3   6090.6                         
## nc1  10 6107.0 6165 -3043.5   6087.0 3.6086      2     0.1646
```

```
nc1c<-lmer(sqrt(eucdisTB)~poly(Generation,2)+(Generation|Folder), data=CWSCREEN_NARROW) 
anova(nc1c,nc1b)#can remove Bgd
```

```
## Data: CWSCREEN_NARROW
## Models:
## nc1c: sqrt(eucdisTB) ~ poly(Generation, 2) + (Generation | Folder)
## nc1b: sqrt(eucdisTB) ~ poly(Generation, 2) + Bgd + (Generation | Folder)
##      Df    AIC    BIC  logLik deviance  Chisq Chi Df Pr(>Chisq)
## nc1c  7 6106.1 6146.8 -3046.1   6092.1                         
## nc1b  8 6106.6 6153.0 -3045.3   6090.6 1.5691      1     0.2103
```

```
nc1d<-lmer(sqrt(eucdisTB)~(Generation|Folder), data=CWSCREEN_NARROW) 
anova(nc1c,nc1d)#significant effect of generation
```

```
## Data: CWSCREEN_NARROW
## Models:
## nc1d: sqrt(eucdisTB) ~ (Generation | Folder)
## nc1c: sqrt(eucdisTB) ~ poly(Generation, 2) + (Generation | Folder)
##      Df    AIC    BIC  logLik deviance  Chisq Chi Df Pr(>Chisq)    
## nc1d  5 6128.3 6157.3 -3059.2   6118.3                             
## nc1c  7 6106.1 6146.8 -3046.1   6092.1 26.181      2  2.064e-06 ***
## ---
## Signif. codes:  0 '***' 0.001 '**' 0.01 '*' 0.05 '.' 0.1 ' ' 1
```

```
#test significance of random effects
rand(nc1c)#random effects not significant
```

```
## ANOVA-like table for random-effects: Single term deletions
## 
## Model:
## sqrt(eucdisTB) ~ poly(Generation, 2) + (Generation | Folder)
##                                     npar  logLik    AIC    LRT Df Pr(>Chisq)
## <none>                                 7 -3046.2 6106.4                     
## Generation in (Generation | Folder)    5 -3047.6 6105.1 2.6541  2     0.2653
```

```
#is the polynomial useful?
nc1c_2<-lmer(sqrt(eucdisTB)~Generation+(Generation|Folder), data=CWSCREEN_NARROW) 
anova(nc1c_2, nc1c)#polynomial needed
```

```
## Data: CWSCREEN_NARROW
## Models:
## nc1c_2: sqrt(eucdisTB) ~ Generation + (Generation | Folder)
## nc1c: sqrt(eucdisTB) ~ poly(Generation, 2) + (Generation | Folder)
##        Df    AIC    BIC  logLik deviance  Chisq Chi Df Pr(>Chisq)   
## nc1c_2  6 6113.4 6148.2 -3050.7   6101.4                            
## nc1c    7 6106.1 6146.8 -3046.1   6092.1 9.2703      1   0.002329 **
## ---
## Signif. codes:  0 '***' 0.001 '**' 0.01 '*' 0.05 '.' 0.1 ' ' 1
```

```
#model without random slope
nc2<-lmer(sqrt(eucdisTB)~poly(Generation,2)*Bgd+(1|Folder), data=CWSCREEN_NARROW)
nc2b<-lmer(sqrt(eucdisTB)~poly(Generation,2)+Bgd+(1|Folder), data=CWSCREEN_NARROW)
anova(nc2, nc2b)#remove Bgd:Generation interaction
```

```
## Data: CWSCREEN_NARROW
## Models:
## nc2b: sqrt(eucdisTB) ~ poly(Generation, 2) + Bgd + (1 | Folder)
## nc2: sqrt(eucdisTB) ~ poly(Generation, 2) * Bgd + (1 | Folder)
##      Df    AIC    BIC  logLik deviance  Chisq Chi Df Pr(>Chisq)
## nc2b  6 6104.4 6139.3 -3046.2   6092.4                         
## nc2   8 6104.8 6151.2 -3044.4   6088.8 3.6196      2     0.1637
```

```
nc2c<-lmer(sqrt(eucdisTB)~poly(Generation,2)+(1|Folder), data=CWSCREEN_NARROW)
anova(nc2c, nc2b)#remove Bgd
```

```
## Data: CWSCREEN_NARROW
## Models:
## nc2c: sqrt(eucdisTB) ~ poly(Generation, 2) + (1 | Folder)
## nc2b: sqrt(eucdisTB) ~ poly(Generation, 2) + Bgd + (1 | Folder)
##      Df    AIC    BIC  logLik deviance  Chisq Chi Df Pr(>Chisq)
## nc2c  5 6104.0 6133.0 -3047.0   6094.0                         
## nc2b  6 6104.4 6139.3 -3046.2   6092.4 1.5682      1     0.2105
```

```
nc2d<-lmer(sqrt(eucdisTB)~(1|Folder), data=CWSCREEN_NARROW)
anova(nc2c, nc2d)#Generation still significant
```

```
## Data: CWSCREEN_NARROW
## Models:
## nc2d: sqrt(eucdisTB) ~ (1 | Folder)
## nc2c: sqrt(eucdisTB) ~ poly(Generation, 2) + (1 | Folder)
##      Df    AIC    BIC  logLik deviance  Chisq Chi Df Pr(>Chisq)    
## nc2d  3 6291.3 6308.7 -3142.7   6285.3                             
## nc2c  5 6104.0 6133.0 -3047.0   6094.0 191.31      2  < 2.2e-16 ***
## ---
## Signif. codes:  0 '***' 0.001 '**' 0.01 '*' 0.05 '.' 0.1 ' ' 1
```

```
#is the polynomial useful?
nc2c_2<-lmer(sqrt(eucdisTB)~Generation+(1|Folder), data=CWSCREEN_NARROW)
anova(nc2c_2,nc2c)#yes
```

```
## Data: CWSCREEN_NARROW
## Models:
## nc2c_2: sqrt(eucdisTB) ~ Generation + (1 | Folder)
## nc2c: sqrt(eucdisTB) ~ poly(Generation, 2) + (1 | Folder)
##        Df    AIC    BIC  logLik deviance  Chisq Chi Df Pr(>Chisq)   
## nc2c_2  4 6111.3 6134.5 -3051.6   6103.3                            
## nc2c    5 6104.0 6133.0 -3047.0   6094.0 9.2474      1   0.002358 **
## ---
## Signif. codes:  0 '***' 0.001 '**' 0.01 '*' 0.05 '.' 0.1 ' ' 1
```

```
#compare final models with and w/o random slope:
AIC(nc1c, nc2c)#lower AIC without random slope
```

```
##      df      AIC
## nc1c  7 6106.449
## nc2c  5 6105.103
```

##### Camouflage metrics: colour difference using weighted average method

```
#start with Generation (2nd order polynomial) and Bgd, interacting 
#+ random effect of Folder (population) with random slope for generation

#as above, but using weighted average method for assessing colour difference

#Normal linear mixed effects models are not appropriate here (very poor diagnostics) due to zero-inflation. In the narrow colour space populations, lots of targets are composed of colours that are all present in the background, leading to lots of 0s.

#Tried manually creating a hurdle-like model by separating the data into two parts - a binomial response (is zero or not?) and a continuous response (if not zero, then?) - but this leads to convergence issuesfor the binomial part.
#The simplest way to analyse these data is to round the colour difference values and apply functions used for count data. Using these data, the same models can be tested here as for all other variables, for the full colour space. Diagnostics are still poor for the narrow colour space (but no zero-inflation), so a negative binomial family is used. 

#create new variable, rounding the weighted average colour difference
CWSCREEN_COLDIST$ColDistRound<-round(CWSCREEN_COLDIST$NewColMap_DeltaS)

CWSCREEN_COLDIST_FULL<-CWSCREEN_COLDIST%>%
  filter(ColSpace=="Full")
CWSCREEN_COLDIST_NARROW<-CWSCREEN_COLDIST%>%
  filter(ColSpace=="Narrow")

#full colour space
fcc1<-lmer(NewColMap_DeltaS~poly(Generation,2)*Bgd+(Generation|Folder), data=CWSCREEN_COLDIST_FULL)
#summary(fcc1)

#test model assumptions
plot(fcc1)
```

```
hist(resid(fcc1))#ok
```

```
#simplify model
fcc1b<-lmer(NewColMap_DeltaS~poly(Generation,2)+Bgd+(Generation|Folder), data=CWSCREEN_COLDIST_FULL) 
anova(fcc1,fcc1b)#can't remove interaction
```

```
## Data: CWSCREEN_COLDIST_FULL
## Models:
## fcc1b: NewColMap_DeltaS ~ poly(Generation, 2) + Bgd + (Generation | 
## fcc1b:     Folder)
## fcc1: NewColMap_DeltaS ~ poly(Generation, 2) * Bgd + (Generation | 
## fcc1:     Folder)
##       Df   AIC   BIC  logLik deviance  Chisq Chi Df Pr(>Chisq)  
## fcc1b  8 16332 16379 -8158.2    16316                           
## fcc1  10 16330 16388 -8154.8    16310 6.9645      2    0.03074 *
## ---
## Signif. codes:  0 '***' 0.001 '**' 0.01 '*' 0.05 '.' 0.1 ' ' 1
```

```
fcc1c<-lmer(NewColMap_DeltaS~Generation*Bgd+(Generation|Folder), data=CWSCREEN_COLDIST_FULL) 
anova(fcc1c,fcc1)#polynomial important
```

```
## Data: CWSCREEN_COLDIST_FULL
## Models:
## fcc1c: NewColMap_DeltaS ~ Generation * Bgd + (Generation | Folder)
## fcc1: NewColMap_DeltaS ~ poly(Generation, 2) * Bgd + (Generation | 
## fcc1:     Folder)
##       Df   AIC   BIC  logLik deviance  Chisq Chi Df Pr(>Chisq)  
## fcc1c  8 16333 16380 -8158.6    16317                           
## fcc1  10 16330 16388 -8154.8    16310 7.7598      2    0.02065 *
## ---
## Signif. codes:  0 '***' 0.001 '**' 0.01 '*' 0.05 '.' 0.1 ' ' 1
```

```
#test significance of random effects
rand(fcc1)
```

```
## ANOVA-like table for random-effects: Single term deletions
## 
## Model:
## NewColMap_DeltaS ~ poly(Generation, 2) + Bgd + (Generation | 
##     Folder) + poly(Generation, 2):Bgd
##                                     npar  logLik   AIC    LRT Df Pr(>Chisq)   
## <none>                                10 -8138.1 16296                        
## Generation in (Generation | Folder)    8 -8145.0 16306 13.732  2   0.001043 **
## ---
## Signif. codes:  0 '***' 0.001 '**' 0.01 '*' 0.05 '.' 0.1 ' ' 1
```

```
#model without random slope
fcc2<-lmer(NewColMap_DeltaS~poly(Generation,2)*Bgd+(1|Folder), data=CWSCREEN_COLDIST_FULL)
fcc2b<-lmer(NewColMap_DeltaS~poly(Generation,2)+Bgd+(1|Folder), data=CWSCREEN_COLDIST_FULL)
anova(fcc2, fcc2b)#can't remove Bgd:Generation interaction
```

```
## Data: CWSCREEN_COLDIST_FULL
## Models:
## fcc2b: NewColMap_DeltaS ~ poly(Generation, 2) + Bgd + (1 | Folder)
## fcc2: NewColMap_DeltaS ~ poly(Generation, 2) * Bgd + (1 | Folder)
##       Df   AIC   BIC  logLik deviance  Chisq Chi Df Pr(>Chisq)    
## fcc2b  6 16366 16401 -8177.2    16354                             
## fcc2   8 16337 16383 -8160.4    16321 33.763      2  4.661e-08 ***
## ---
## Signif. codes:  0 '***' 0.001 '**' 0.01 '*' 0.05 '.' 0.1 ' ' 1
```

```
fcc2c<-lmer(NewColMap_DeltaS~Generation*Bgd+(1|Folder), data=CWSCREEN_COLDIST_FULL)
anova(fcc2c, fcc2)#need polynomial
```

```
## Data: CWSCREEN_COLDIST_FULL
## Models:
## fcc2c: NewColMap_DeltaS ~ Generation * Bgd + (1 | Folder)
## fcc2: NewColMap_DeltaS ~ poly(Generation, 2) * Bgd + (1 | Folder)
##       Df   AIC   BIC  logLik deviance  Chisq Chi Df Pr(>Chisq)  
## fcc2c  6 16340 16375 -8164.2    16328                           
## fcc2   8 16337 16383 -8160.4    16321 7.7037      2    0.02124 *
## ---
## Signif. codes:  0 '***' 0.001 '**' 0.01 '*' 0.05 '.' 0.1 ' ' 1
```

```
#compare final models with and w/o random slope:
AIC(fcc1, fcc2)#lower AIC with random slope
```

```
##      df      AIC
## fcc1 10 16296.17
## fcc2  8 16305.90
```

```
#narrow colour space

ncc1<-glmmTMB(ColDistRound ~ poly(Generation,2)*Bgd+(Generation|Folder), data=CWSCREEN_COLDIST_NARROW, family=nbinom1 )
#check assumptions
res<-simulateResiduals(ncc1, plot=T)
```

```
#check dispersion
par(mfrow = c(1,2))
testDispersion(res)
```

```
## 
##  DHARMa nonparametric dispersion test via sd of residuals fitted vs.
##  simulated
## 
## data:  simulationOutput
## ratioObsSim = 0.98189, p-value = 1
## alternative hypothesis: two.sided
```

```
testZeroInflation(res)# no underdispersion, no zero-inflation
```

```
## 
##  DHARMa zero-inflation test via comparison to expected zeros with
##  simulation under H0 = fitted model
## 
## data:  simulationOutput
## ratioObsSim = 0.99636, p-value = 0.864
## alternative hypothesis: two.sided
```

```
#model simplification
ncc1b<-glmmTMB(ColDistRound ~ poly(Generation,2)+Bgd+(Generation|Folder), data=CWSCREEN_COLDIST_NARROW, family=nbinom1 )
anova(ncc1, ncc1b)#interaction significant
```

```
## Data: CWSCREEN_COLDIST_NARROW
## Models:
## ncc1b: ColDistRound ~ poly(Generation, 2) + Bgd + (Generation | Folder), zi=~0, disp=~1
## ncc1: ColDistRound ~ poly(Generation, 2) * Bgd + (Generation | Folder), zi=~0, disp=~1
##       Df    AIC    BIC  logLik deviance  Chisq Chi Df Pr(>Chisq)  
## ncc1b  8 4517.2 4563.7 -2250.6   4501.2                           
## ncc1  10 4513.8 4571.8 -2246.9   4493.8 7.4339      2    0.02431 *
## ---
## Signif. codes:  0 '***' 0.001 '**' 0.01 '*' 0.05 '.' 0.1 ' ' 1
```

```
#summary(ncc1)

#is the polynomial useful?
ncc1_2<-glmmTMB(ColDistRound ~ Generation*Bgd+(Generation|Folder), data=CWSCREEN_COLDIST_NARROW, family=nbinom1 )
anova(ncc1_2, ncc1)#yes
```

```
## Data: CWSCREEN_COLDIST_NARROW
## Models:
## ncc1_2: ColDistRound ~ Generation * Bgd + (Generation | Folder), zi=~0, disp=~1
## ncc1: ColDistRound ~ poly(Generation, 2) * Bgd + (Generation | Folder), zi=~0, disp=~1
##        Df    AIC    BIC  logLik deviance  Chisq Chi Df Pr(>Chisq)  
## ncc1_2  8 4517.6 4564.1 -2250.8   4501.6                           
## ncc1   10 4513.8 4571.8 -2246.9   4493.8 7.8275      2    0.01997 *
## ---
## Signif. codes:  0 '***' 0.001 '**' 0.01 '*' 0.05 '.' 0.1 ' ' 1
```

```
#try model without random slope for generation
ncc2<-glmmTMB(ColDistRound ~ poly(Generation,2)*Bgd+(1|Folder), data=CWSCREEN_COLDIST_NARROW, family=nbinom1 )
#check assumptions
res<-simulateResiduals(ncc2, plot=T)
```

```
#check dispersion
par(mfrow = c(1,2))
testDispersion(res)
```

```
## 
##  DHARMa nonparametric dispersion test via sd of residuals fitted vs.
##  simulated
## 
## data:  simulationOutput
## ratioObsSim = 1.0188, p-value = 0.784
## alternative hypothesis: two.sided
```

```
testZeroInflation(res)#no underdispersion, no zero-inflation
```

```
## 
##  DHARMa zero-inflation test via comparison to expected zeros with
##  simulation under H0 = fitted model
## 
## data:  simulationOutput
## ratioObsSim = 0.9917, p-value = 0.808
## alternative hypothesis: two.sided
```

```
#model simplification
ncc2b<-glmmTMB(ColDistRound ~ poly(Generation,2)+Bgd+(1|Folder), data=CWSCREEN_COLDIST_NARROW, family=nbinom1 )
anova(ncc2, ncc2b)#interaction is significant
```

```
## Data: CWSCREEN_COLDIST_NARROW
## Models:
## ncc2b: ColDistRound ~ poly(Generation, 2) + Bgd + (1 | Folder), zi=~0, disp=~1
## ncc2: ColDistRound ~ poly(Generation, 2) * Bgd + (1 | Folder), zi=~0, disp=~1
##       Df    AIC    BIC  logLik deviance  Chisq Chi Df Pr(>Chisq)  
## ncc2b  6 4556.2 4591.0 -2272.1   4544.2                           
## ncc2   8 4552.9 4599.3 -2268.4   4536.9 7.3128      2    0.02583 *
## ---
## Signif. codes:  0 '***' 0.001 '**' 0.01 '*' 0.05 '.' 0.1 ' ' 1
```

```
#summary(ncc2)
#is polynomial useful?
ncc2_2<-glmmTMB(ColDistRound ~ Generation*Bgd+(1|Folder), data=CWSCREEN_COLDIST_NARROW, family=nbinom1 )
anova(ncc2, ncc2_2)#yes
```

```
## Data: CWSCREEN_COLDIST_NARROW
## Models:
## ncc2_2: ColDistRound ~ Generation * Bgd + (1 | Folder), zi=~0, disp=~1
## ncc2: ColDistRound ~ poly(Generation, 2) * Bgd + (1 | Folder), zi=~0, disp=~1
##        Df    AIC    BIC  logLik deviance  Chisq Chi Df Pr(>Chisq)  
## ncc2_2  6 4557.6 4592.4 -2272.8   4545.6                           
## ncc2    8 4552.9 4599.3 -2268.4   4536.9 8.7366      2    0.01267 *
## ---
## Signif. codes:  0 '***' 0.001 '**' 0.01 '*' 0.05 '.' 0.1 ' ' 1
```

```
#compare best models with and without random slope
AIC(ncc1,ncc2)
```

```
##      df      AIC
## ncc1 10 4513.805
## ncc2  8 4552.877
```

##### Camouflage metrics: edge disruption

```
#start with Generation (2nd order polynomial) and Bgd, interacting 
#+ random effect of Folder(Population) with random slope for generation

#full
fgr1<-lmer(Target_L_GabRat~poly(Generation,2)*Bgd+(Generation|Folder), data=CWSCREEN_FULL)

#test model assumptions
res<-simulateResiduals(fgr1, plot=T)
```

```
par(mfrow = c(1,1))
hist(resid(fgr1))
```

```
plot(fgr1)
```

```
#model simplification
fgr1b<-lmer(Target_L_GabRat~poly(Generation,2)+Bgd+(Generation|Folder), data=CWSCREEN_FULL)
anova(fgr1, fgr1b)#remove interaction
```

```
## Data: CWSCREEN_FULL
## Models:
## fgr1b: Target_L_GabRat ~ poly(Generation, 2) + Bgd + (Generation | Folder)
## fgr1: Target_L_GabRat ~ poly(Generation, 2) * Bgd + (Generation | Folder)
##       Df     AIC     BIC logLik deviance  Chisq Chi Df Pr(>Chisq)
## fgr1b  8 -6341.8 -6295.4 3178.9  -6357.8                         
## fgr1  10 -6338.3 -6280.3 3179.1  -6358.3 0.4808      2     0.7863
```

```
fgr1c<-lmer(Target_L_GabRat~poly(Generation,2)+(Generation|Folder), data=CWSCREEN_FULL)
anova(fgr1c, fgr1b)#remove Bgd
```

```
## Data: CWSCREEN_FULL
## Models:
## fgr1c: Target_L_GabRat ~ poly(Generation, 2) + (Generation | Folder)
## fgr1b: Target_L_GabRat ~ poly(Generation, 2) + Bgd + (Generation | Folder)
##       Df     AIC     BIC logLik deviance  Chisq Chi Df Pr(>Chisq)
## fgr1c  7 -6343.4 -6302.7 3178.7  -6357.4                         
## fgr1b  8 -6341.8 -6295.4 3178.9  -6357.8 0.4436      1     0.5054
```

```
fgr1d<-lmer(Target_L_GabRat~(Generation|Folder), data=CWSCREEN_FULL)
anova(fgr1c, fgr1d)#remove Generation
```

```
## Data: CWSCREEN_FULL
## Models:
## fgr1d: Target_L_GabRat ~ (Generation | Folder)
## fgr1c: Target_L_GabRat ~ poly(Generation, 2) + (Generation | Folder)
##       Df     AIC     BIC logLik deviance  Chisq Chi Df Pr(>Chisq)
## fgr1d  5 -6345.3 -6316.3 3177.7  -6355.3                         
## fgr1c  7 -6343.4 -6302.7 3178.7  -6357.4 2.0297      2     0.3625
```

```
#model without random slope
fgr2<-lmer(Target_L_GabRat~poly(Generation,2)*Bgd+(1|Folder), data=CWSCREEN_FULL)
fgr2b<-lmer(Target_L_GabRat~poly(Generation,2)+Bgd+(1|Folder), data=CWSCREEN_FULL)
anova(fgr2, fgr2b)#remove interaction
```

```
## Data: CWSCREEN_FULL
## Models:
## fgr2b: Target_L_GabRat ~ poly(Generation, 2) + Bgd + (1 | Folder)
## fgr2: Target_L_GabRat ~ poly(Generation, 2) * Bgd + (1 | Folder)
##       Df     AIC     BIC logLik deviance  Chisq Chi Df Pr(>Chisq)
## fgr2b  6 -6236.9 -6202.1 3124.5  -6248.9                         
## fgr2   8 -6233.8 -6187.4 3124.9  -6249.8 0.8965      2     0.6387
```

```
fgr2c<-lmer(Target_L_GabRat~poly(Generation,2)+(1|Folder), data=CWSCREEN_FULL)
anova(fgr2c, fgr2b)#remove Bgd
```

```
## Data: CWSCREEN_FULL
## Models:
## fgr2c: Target_L_GabRat ~ poly(Generation, 2) + (1 | Folder)
## fgr2b: Target_L_GabRat ~ poly(Generation, 2) + Bgd + (1 | Folder)
##       Df     AIC     BIC logLik deviance  Chisq Chi Df Pr(>Chisq)
## fgr2c  5 -6238.8 -6209.8 3124.4  -6248.8                         
## fgr2b  6 -6236.9 -6202.1 3124.5  -6248.9 0.0906      1     0.7634
```

```
fgr2d<-lmer(Target_L_GabRat~(1|Folder), data=CWSCREEN_FULL)
anova(fgr2c, fgr2d)#can't remove Generation
```

```
## Data: CWSCREEN_FULL
## Models:
## fgr2d: Target_L_GabRat ~ (1 | Folder)
## fgr2c: Target_L_GabRat ~ poly(Generation, 2) + (1 | Folder)
##       Df     AIC     BIC logLik deviance  Chisq Chi Df Pr(>Chisq)  
## fgr2d  3 -6236.1 -6218.7 3121.1  -6242.1                           
## fgr2c  5 -6238.8 -6209.8 3124.4  -6248.8 6.7083      2    0.03494 *
## ---
## Signif. codes:  0 '***' 0.001 '**' 0.01 '*' 0.05 '.' 0.1 ' ' 1
```

```
#summary(fgr2c)

#is polynomial useful?
fgr2c_2<-lmer(Target_L_GabRat~Generation+(1|Folder), data=CWSCREEN_FULL)
anova(fgr2c_2, fgr2c)#no
```

```
## Data: CWSCREEN_FULL
## Models:
## fgr2c_2: Target_L_GabRat ~ Generation + (1 | Folder)
## fgr2c: Target_L_GabRat ~ poly(Generation, 2) + (1 | Folder)
##         Df     AIC     BIC logLik deviance  Chisq Chi Df Pr(>Chisq)
## fgr2c_2  4 -6239.1 -6215.9 3123.6  -6247.1                         
## fgr2c    5 -6238.8 -6209.8 3124.4  -6248.8 1.7055      1     0.1916
```

```
#further simplification
fgr2d_2<-lmer(Target_L_GabRat~(1|Folder), data=CWSCREEN_FULL)
anova(fgr2d_2, fgr2c_2)#still a significant effect of generation
```

```
## Data: CWSCREEN_FULL
## Models:
## fgr2d_2: Target_L_GabRat ~ (1 | Folder)
## fgr2c_2: Target_L_GabRat ~ Generation + (1 | Folder)
##         Df     AIC     BIC logLik deviance  Chisq Chi Df Pr(>Chisq)  
## fgr2d_2  3 -6236.1 -6218.7 3121.1  -6242.1                           
## fgr2c_2  4 -6239.1 -6215.9 3123.6  -6247.1 5.0029      1    0.02531 *
## ---
## Signif. codes:  0 '***' 0.001 '**' 0.01 '*' 0.05 '.' 0.1 ' ' 1
```

```
#compare models with and w/o random effects
AIC(fgr2c_2, fgr1d)
```

```
##         df       AIC
## fgr2c_2  4 -6217.703
## fgr1d    5 -6337.540
```

```
#narrow
ngr1<-lmer(Target_L_GabRat~poly(Generation,2)*Bgd+(Generation|Folder), data=CWSCREEN_NARROW)
#model fails to converge - use model w/o slope

#model without random slope
ngr2<-lmer(Target_L_GabRat~poly(Generation,2)*Bgd+(1|Folder), data=CWSCREEN_NARROW)
hist(resid(ngr2))
```

```
plot(ngr2)
```

```
ngr2b<-lmer(Target_L_GabRat~poly(Generation,2)+Bgd+(1|Folder), data=CWSCREEN_NARROW)
anova(ngr2, ngr2b)#remove interaction
```

```
## Data: CWSCREEN_NARROW
## Models:
## ngr2b: Target_L_GabRat ~ poly(Generation, 2) + Bgd + (1 | Folder)
## ngr2: Target_L_GabRat ~ poly(Generation, 2) * Bgd + (1 | Folder)
##       Df     AIC     BIC logLik deviance  Chisq Chi Df Pr(>Chisq)
## ngr2b  6 -8036.0 -8001.1 4024.0  -8048.0                         
## ngr2   8 -8034.3 -7987.9 4025.2  -8050.3 2.3629      2     0.3068
```

```
ngr2c<-lmer(Target_L_GabRat~poly(Generation,2)+(1|Folder), data=CWSCREEN_NARROW)
anova(ngr2c, ngr2b)#remove Bgd
```

```
## Data: CWSCREEN_NARROW
## Models:
## ngr2c: Target_L_GabRat ~ poly(Generation, 2) + (1 | Folder)
## ngr2b: Target_L_GabRat ~ poly(Generation, 2) + Bgd + (1 | Folder)
##       Df     AIC     BIC logLik deviance  Chisq Chi Df Pr(>Chisq)
## ngr2c  5 -8037.9 -8008.9   4024  -8047.9                         
## ngr2b  6 -8036.0 -8001.1   4024  -8048.0 0.0199      1     0.8879
```

```
ngr2d<-lmer(Target_L_GabRat~(1|Folder), data=CWSCREEN_NARROW)
anova(ngr2c, ngr2d)#can't remove Generation
```

```
## Data: CWSCREEN_NARROW
## Models:
## ngr2d: Target_L_GabRat ~ (1 | Folder)
## ngr2c: Target_L_GabRat ~ poly(Generation, 2) + (1 | Folder)
##       Df     AIC     BIC logLik deviance  Chisq Chi Df Pr(>Chisq)  
## ngr2d  3 -8033.4 -8016.0 4019.7  -8039.4                           
## ngr2c  5 -8037.9 -8008.9 4024.0  -8047.9 8.5564      2    0.01387 *
## ---
## Signif. codes:  0 '***' 0.001 '**' 0.01 '*' 0.05 '.' 0.1 ' ' 1
```

```
#summary(ngr2c)

#is polynomial useful?
ngr2c_2<-lmer(Target_L_GabRat~Generation+(1|Folder), data=CWSCREEN_NARROW)
anova(ngr2c_2, ngr2c)#no
```

```
## Data: CWSCREEN_NARROW
## Models:
## ngr2c_2: Target_L_GabRat ~ Generation + (1 | Folder)
## ngr2c: Target_L_GabRat ~ poly(Generation, 2) + (1 | Folder)
##         Df     AIC     BIC logLik deviance  Chisq Chi Df Pr(>Chisq)
## ngr2c_2  4 -8039.4 -8016.2 4023.7  -8047.4                         
## ngr2c    5 -8037.9 -8008.9 4024.0  -8047.9 0.5136      1     0.4736
```

```
#further simplification
ngr2d_2<-lmer(Target_L_GabRat~(1|Folder), data=CWSCREEN_NARROW)
anova(ngr2d_2, ngr2c_2)#still a significant effect of generation
```

```
## Data: CWSCREEN_NARROW
## Models:
## ngr2d_2: Target_L_GabRat ~ (1 | Folder)
## ngr2c_2: Target_L_GabRat ~ Generation + (1 | Folder)
##         Df     AIC     BIC logLik deviance  Chisq Chi Df Pr(>Chisq)   
## ngr2d_2  3 -8033.4 -8016.0 4019.7  -8039.4                            
## ngr2c_2  4 -8039.4 -8016.2 4023.7  -8047.4 8.0428      1   0.004568 **
## ---
## Signif. codes:  0 '***' 0.001 '**' 0.01 '*' 0.05 '.' 0.1 ' ' 1
```

##### Camouflage metrics: pattern difference

```
#start with Generation (2nd order polynomial) and Bgd, interacting 
#+ random effect of Folder(population) with random slope for generation

CWSCREEN_PATTERN_FULL<-CWSCREEN_PATTERN%>%
  filter(ColSpace=="Full")
CWSCREEN_PATTERN_NARROW<-CWSCREEN_PATTERN%>%
  filter(ColSpace=="Narrow")

#full colour space
fp1<-lmer(pattern_energy_diff~poly(Generation,2)*Bgd+(Generation|Folder), data=CWSCREEN_PATTERN_FULL)

#test model assumptions
res<-simulateResiduals(fp1, plot=T)
```

```
hist(resid(fp1))
```

```
plot(fp1)#ok - no transforms improve diagnostics
```

```
#model simplification
fp1b<-lmer(pattern_energy_diff~poly(Generation,2)+Bgd+(Generation|Folder), data=CWSCREEN_PATTERN_FULL)
anova(fp1,fp1b)#remove interaction
```

```
## Data: CWSCREEN_PATTERN_FULL
## Models:
## fp1b: pattern_energy_diff ~ poly(Generation, 2) + Bgd + (Generation | 
## fp1b:     Folder)
## fp1: pattern_energy_diff ~ poly(Generation, 2) * Bgd + (Generation | 
## fp1:     Folder)
##      Df   AIC   BIC  logLik deviance  Chisq Chi Df Pr(>Chisq)
## fp1b  8 17020 17066 -8501.8    17004                         
## fp1  10 17019 17077 -8499.6    16999 4.4542      2     0.1078
```

```
fp1c<-lmer(pattern_energy_diff~poly(Generation,2)+(Generation|Folder), data=CWSCREEN_PATTERN_FULL)
anova(fp1c,fp1b)#remove Bgd
```

```
## Data: CWSCREEN_PATTERN_FULL
## Models:
## fp1c: pattern_energy_diff ~ poly(Generation, 2) + (Generation | Folder)
## fp1b: pattern_energy_diff ~ poly(Generation, 2) + Bgd + (Generation | 
## fp1b:     Folder)
##      Df   AIC   BIC  logLik deviance  Chisq Chi Df Pr(>Chisq)
## fp1c  7 17018 17058 -8501.9    17004                         
## fp1b  8 17020 17066 -8501.8    17004 0.0196      1     0.8885
```

```
fp1d<-lmer(pattern_energy_diff~(Generation|Folder), data=CWSCREEN_PATTERN_FULL)
anova(fp1c,fp1d)#remove Generation
```

```
## Data: CWSCREEN_PATTERN_FULL
## Models:
## fp1d: pattern_energy_diff ~ (Generation | Folder)
## fp1c: pattern_energy_diff ~ poly(Generation, 2) + (Generation | Folder)
##      Df   AIC   BIC  logLik deviance Chisq Chi Df Pr(>Chisq)
## fp1d  5 17017 17046 -8503.3    17007                        
## fp1c  7 17018 17058 -8501.9    17004 2.788      2     0.2481
```

```
rand(fp1d)
```

```
## ANOVA-like table for random-effects: Single term deletions
## 
## Model:
## pattern_energy_diff ~ (Generation | Folder)
##                                     npar  logLik   AIC    LRT Df Pr(>Chisq)    
## <none>                                 5 -8502.5 17015                         
## Generation in (Generation | Folder)    3 -8595.7 17197 186.43  2  < 2.2e-16 ***
## ---
## Signif. codes:  0 '***' 0.001 '**' 0.01 '*' 0.05 '.' 0.1 ' ' 1
```

```
#model without random slope
fp2<-lmer(pattern_energy_diff~poly(Generation,2)*Bgd+(1|Folder), data=CWSCREEN_PATTERN_FULL)
fp2b<-lmer(pattern_energy_diff~poly(Generation,2)+Bgd+(1|Folder), data=CWSCREEN_PATTERN_FULL)
anova(fp2,fp2b)#can't remove  interaction
```

```
## Data: CWSCREEN_PATTERN_FULL
## Models:
## fp2b: pattern_energy_diff ~ poly(Generation, 2) + Bgd + (1 | Folder)
## fp2: pattern_energy_diff ~ poly(Generation, 2) * Bgd + (1 | Folder)
##      Df   AIC   BIC  logLik deviance  Chisq Chi Df Pr(>Chisq)   
## fp2b  6 17203 17237 -8595.3    17191                            
## fp2   8 17197 17244 -8590.6    17181 9.4713      2   0.008777 **
## ---
## Signif. codes:  0 '***' 0.001 '**' 0.01 '*' 0.05 '.' 0.1 ' ' 1
```

```
#is polynomial useful?
fp2_2<-lmer(pattern_energy_diff~Generation*Bgd+(1|Folder), data=CWSCREEN_PATTERN_FULL)
anova(fp2, fp2_2)#yes
```

```
## Data: CWSCREEN_PATTERN_FULL
## Models:
## fp2_2: pattern_energy_diff ~ Generation * Bgd + (1 | Folder)
## fp2: pattern_energy_diff ~ poly(Generation, 2) * Bgd + (1 | Folder)
##       Df   AIC   BIC  logLik deviance  Chisq Chi Df Pr(>Chisq)  
## fp2_2  6 17200 17234 -8593.8    17188                           
## fp2    8 17197 17244 -8590.6    17181 6.4956      2    0.03886 *
## ---
## Signif. codes:  0 '***' 0.001 '**' 0.01 '*' 0.05 '.' 0.1 ' ' 1
```

```
#summary(fp2)

#compare models with and w/o random slope
AIC(fp1d,fp2)#model with random slope is best
```

```
##      df      AIC
## fp1d  5 17015.01
## fp2   8 17162.25
```

```
#narrow colour space
np1<-lmer(pattern_energy_diff~poly(Generation,2)*Bgd+(Generation|Folder), data=CWSCREEN_PATTERN_NARROW)
#test model assumptions
res<-simulateResiduals(np1, plot=T)
```

```
hist(resid(np1))
```

```
plot(np1)#ok - no transforms improve diagnostics
```

```
#model simplification
np1b<-lmer(pattern_energy_diff~poly(Generation,2)+Bgd+(Generation|Folder), data=CWSCREEN_PATTERN_NARROW)
anova(np1,np1b)#remove interaction
```

```
## Data: CWSCREEN_PATTERN_NARROW
## Models:
## np1b: pattern_energy_diff ~ poly(Generation, 2) + Bgd + (Generation | 
## np1b:     Folder)
## np1: pattern_energy_diff ~ poly(Generation, 2) * Bgd + (Generation | 
## np1:     Folder)
##      Df   AIC   BIC  logLik deviance  Chisq Chi Df Pr(>Chisq)
## np1b  8 12542 12588 -6262.9    12526                         
## np1  10 12546 12604 -6262.7    12526 0.2565      2     0.8796
```

```
np1c<-lmer(pattern_energy_diff~poly(Generation,2)+(Generation|Folder), data=CWSCREEN_PATTERN_NARROW)
anova(np1c,np1b)#remove Bgd
```

```
## Data: CWSCREEN_PATTERN_NARROW
## Models:
## np1c: pattern_energy_diff ~ poly(Generation, 2) + (Generation | Folder)
## np1b: pattern_energy_diff ~ poly(Generation, 2) + Bgd + (Generation | 
## np1b:     Folder)
##      Df   AIC   BIC  logLik deviance  Chisq Chi Df Pr(>Chisq)
## np1c  7 12540 12581 -6263.1    12526                         
## np1b  8 12542 12588 -6262.9    12526 0.5383      1     0.4631
```

```
np1d<-lmer(pattern_energy_diff~(Generation|Folder), data=CWSCREEN_PATTERN_NARROW)
anova(np1c,np1d)#can't remove Generation
```

```
## Data: CWSCREEN_PATTERN_NARROW
## Models:
## np1d: pattern_energy_diff ~ (Generation | Folder)
## np1c: pattern_energy_diff ~ poly(Generation, 2) + (Generation | Folder)
##      Df   AIC   BIC  logLik deviance  Chisq Chi Df Pr(>Chisq)   
## np1d  5 12547 12576 -6268.4    12537                            
## np1c  7 12540 12581 -6263.1    12526 10.471      2   0.005324 **
## ---
## Signif. codes:  0 '***' 0.001 '**' 0.01 '*' 0.05 '.' 0.1 ' ' 1
```

```
#is polynomial needed?
np1c_2<-lmer(pattern_energy_diff~Generation+(Generation|Folder), data=CWSCREEN_PATTERN_NARROW)
anova(np1c,np1c_2)#no
```

```
## Data: CWSCREEN_PATTERN_NARROW
## Models:
## np1c_2: pattern_energy_diff ~ Generation + (Generation | Folder)
## np1c: pattern_energy_diff ~ poly(Generation, 2) + (Generation | Folder)
##        Df   AIC   BIC  logLik deviance  Chisq Chi Df Pr(>Chisq)
## np1c_2  6 12540 12575 -6264.1    12528                         
## np1c    7 12540 12581 -6263.1    12526 2.0048      1     0.1568
```

```
#further simplification?
np1c_2b<-lmer(pattern_energy_diff~(Generation|Folder), data=CWSCREEN_PATTERN_NARROW)
anova(np1c_2b,np1c_2)#Generation still significant
```

```
## Data: CWSCREEN_PATTERN_NARROW
## Models:
## np1c_2b: pattern_energy_diff ~ (Generation | Folder)
## np1c_2: pattern_energy_diff ~ Generation + (Generation | Folder)
##         Df   AIC   BIC  logLik deviance  Chisq Chi Df Pr(>Chisq)   
## np1c_2b  5 12547 12576 -6268.4    12537                            
## np1c_2   6 12540 12575 -6264.1    12528 8.4664      1   0.003618 **
## ---
## Signif. codes:  0 '***' 0.001 '**' 0.01 '*' 0.05 '.' 0.1 ' ' 1
```

```
#model without random slope
np2<-lmer(pattern_energy_diff~poly(Generation,2)*Bgd+(1|Folder), data=CWSCREEN_PATTERN_NARROW)
np2b<-lmer(pattern_energy_diff~poly(Generation,2)+Bgd+(1|Folder), data=CWSCREEN_PATTERN_NARROW)
anova(np2,np2b)#remove  interaction
```

```
## Data: CWSCREEN_PATTERN_NARROW
## Models:
## np2b: pattern_energy_diff ~ poly(Generation, 2) + Bgd + (1 | Folder)
## np2: pattern_energy_diff ~ poly(Generation, 2) * Bgd + (1 | Folder)
##      Df   AIC   BIC  logLik deviance  Chisq Chi Df Pr(>Chisq)
## np2b  6 12546 12580 -6266.8    12534                         
## np2   8 12549 12596 -6266.6    12533 0.4981      2     0.7795
```

```
np2c<-lmer(pattern_energy_diff~poly(Generation,2)+(1|Folder), data=CWSCREEN_PATTERN_NARROW)
anova(np2c,np2b)#remove Bgd
```

```
## Data: CWSCREEN_PATTERN_NARROW
## Models:
## np2c: pattern_energy_diff ~ poly(Generation, 2) + (1 | Folder)
## np2b: pattern_energy_diff ~ poly(Generation, 2) + Bgd + (1 | Folder)
##      Df   AIC   BIC  logLik deviance  Chisq Chi Df Pr(>Chisq)
## np2c  5 12544 12573 -6266.8    12534                         
## np2b  6 12546 12580 -6266.8    12534 0.0313      1     0.8596
```

```
np2d<-lmer(pattern_energy_diff~(1|Folder), data=CWSCREEN_PATTERN_NARROW)
anova(np2c,np2d)#need Generation
```

```
## Data: CWSCREEN_PATTERN_NARROW
## Models:
## np2d: pattern_energy_diff ~ (1 | Folder)
## np2c: pattern_energy_diff ~ poly(Generation, 2) + (1 | Folder)
##      Df   AIC   BIC  logLik deviance  Chisq Chi Df Pr(>Chisq)    
## np2d  3 12584 12601 -6289.0    12578                             
## np2c  5 12544 12573 -6266.8    12534 44.267      2  2.441e-10 ***
## ---
## Signif. codes:  0 '***' 0.001 '**' 0.01 '*' 0.05 '.' 0.1 ' ' 1
```

```
#is polynomial useful?
np2c_2<-lmer(pattern_energy_diff~Generation+(1|Folder), data=CWSCREEN_PATTERN_NARROW)
anova(np2c_2, np2c)#no
```

```
## Data: CWSCREEN_PATTERN_NARROW
## Models:
## np2c_2: pattern_energy_diff ~ Generation + (1 | Folder)
## np2c: pattern_energy_diff ~ poly(Generation, 2) + (1 | Folder)
##        Df   AIC   BIC  logLik deviance  Chisq Chi Df Pr(>Chisq)
## np2c_2  4 12544 12567 -6267.8    12536                         
## np2c    5 12544 12573 -6266.8    12534 1.9984      1     0.1575
```

```
anova(np2c_2,np2d)#Generation still significant
```

```
## Data: CWSCREEN_PATTERN_NARROW
## Models:
## np2d: pattern_energy_diff ~ (1 | Folder)
## np2c_2: pattern_energy_diff ~ Generation + (1 | Folder)
##        Df   AIC   BIC  logLik deviance  Chisq Chi Df Pr(>Chisq)    
## np2d    3 12584 12601 -6289.0    12578                             
## np2c_2  4 12544 12567 -6267.8    12536 42.268      1  7.957e-11 ***
## ---
## Signif. codes:  0 '***' 0.001 '**' 0.01 '*' 0.05 '.' 0.1 ' ' 1
```

```
#summary(np2c_2)

#compare models with and w/o random slope
AIC(np1c_2,np2c_2)#model with random slope is best
```

```
##        df      AIC
## np1c_2  6 12547.29
## np2c_2  4 12550.86
```

##### Capture time

How does capture time change across generations, in populations on different backgrounds & using different colour spaces?

```
#remove generation 16 from capture time data, as this was not played by participants
#and restrict to first-clicked target per screen (ie. rank 1)
CWSCREEN15<-CWSCREEN%>%
  filter(Generation!=16)%>%
  filter(Rank==1)
#separate full and narrow colour spaces for subsequent analyses
CWSCREEN15_FULL<-CWSCREEN15%>%
  filter(ColSpace=="Full")
CWSCREEN15_NARROW<-CWSCREEN15%>%
  filter(ColSpace=="Narrow")

#linear mixed effects models for each colour space in turn
#include Generation (2nd order polynomial) and Bgd, interacting + random effect of Folder(Population) with random slope for generation
fct1<-lmer(Capture_Time~poly(Generation,2)*Bgd+(Generation|Folder), data=CWSCREEN15_FULL)
#summary(fct1)

#test model assumptions
plot(fct1)
```

```
hist(resid(fct1))
```

```
#very poor, try log transform
fct1<-lmer(log(Capture_Time)~poly(Generation,2)*Bgd+(Generation|Folder), data=CWSCREEN15_FULL) 
hist(resid(fct1))
```

```
plot(fct1)#better
```

```
#simplify model
fct1b<-lmer(log(Capture_Time)~poly(Generation,2)+Bgd+(Generation|Folder), data=CWSCREEN15_FULL) 
anova(fct1,fct1b)#significant effect of background interaction
```

```
## Data: CWSCREEN15_FULL
## Models:
## fct1b: log(Capture_Time) ~ poly(Generation, 2) + Bgd + (Generation | 
## fct1b:     Folder)
## fct1: log(Capture_Time) ~ poly(Generation, 2) * Bgd + (Generation | 
## fct1:     Folder)
##       Df     AIC     BIC logLik deviance  Chisq Chi Df Pr(>Chisq)   
## fct1b  8 -216.44 -181.59 116.22  -232.44                            
## fct1  10 -223.27 -179.71 121.64  -243.27 10.832      2   0.004445 **
## ---
## Signif. codes:  0 '***' 0.001 '**' 0.01 '*' 0.05 '.' 0.1 ' ' 1
```

```
#test significance of random effects
rand(fct1)
```

```
## ANOVA-like table for random-effects: Single term deletions
## 
## Model:
## log(Capture_Time) ~ poly(Generation, 2) + Bgd + (Generation | 
##     Folder) + poly(Generation, 2):Bgd
##                                     npar logLik     AIC    LRT Df Pr(>Chisq)   
## <none>                                10 116.96 -213.93                        
## Generation in (Generation | Folder)    8 111.59 -207.19 10.739  2   0.004657 **
## ---
## Signif. codes:  0 '***' 0.001 '**' 0.01 '*' 0.05 '.' 0.1 ' ' 1
```

```
#is the polynomial useful?
fct1_2<-lmer(log(Capture_Time)~Generation*Bgd+(Generation|Folder), data=CWSCREEN15_FULL) 
anova(fct1, fct1_2)#significant effect of polynomial, so keep it in final model
```

```
## Data: CWSCREEN15_FULL
## Models:
## fct1_2: log(Capture_Time) ~ Generation * Bgd + (Generation | Folder)
## fct1: log(Capture_Time) ~ poly(Generation, 2) * Bgd + (Generation | 
## fct1:     Folder)
##        Df     AIC     BIC logLik deviance  Chisq Chi Df Pr(>Chisq)    
## fct1_2  8 -211.45 -176.60 113.72  -227.45                             
## fct1   10 -223.27 -179.71 121.64  -243.27 15.824      2  0.0003663 ***
## ---
## Signif. codes:  0 '***' 0.001 '**' 0.01 '*' 0.05 '.' 0.1 ' ' 1
```

```
#model without random slope
fct2<-lmer(log(Capture_Time)~poly(Generation,2)*Bgd+(1|Folder), data=CWSCREEN15_FULL)
fct2b<-lmer(log(Capture_Time)~poly(Generation,2)+Bgd+(1|Folder), data=CWSCREEN15_FULL)
anova(fct2, fct2b)#can't remove Bgd:Generation interaction
```

```
## Data: CWSCREEN15_FULL
## Models:
## fct2b: log(Capture_Time) ~ poly(Generation, 2) + Bgd + (1 | Folder)
## fct2: log(Capture_Time) ~ poly(Generation, 2) * Bgd + (1 | Folder)
##       Df     AIC     BIC logLik deviance  Chisq Chi Df Pr(>Chisq)    
## fct2b  6 -206.63 -180.50 109.32  -218.63                             
## fct2   8 -218.58 -183.73 117.29  -234.58 15.949      2  0.0003442 ***
## ---
## Signif. codes:  0 '***' 0.001 '**' 0.01 '*' 0.05 '.' 0.1 ' ' 1
```

```
#is the polynomial useful?
fct2_2<-lmer(log(Capture_Time)~Generation*Bgd+(1|Folder), data=CWSCREEN15_FULL)
anova(fct2, fct2_2)#yes, significant effect of polynomial
```

```
## Data: CWSCREEN15_FULL
## Models:
## fct2_2: log(Capture_Time) ~ Generation * Bgd + (1 | Folder)
## fct2: log(Capture_Time) ~ poly(Generation, 2) * Bgd + (1 | Folder)
##        Df     AIC     BIC logLik deviance  Chisq Chi Df Pr(>Chisq)    
## fct2_2  6 -207.13 -181.00 109.57  -219.13                             
## fct2    8 -218.58 -183.73 117.29  -234.58 15.447      2  0.0004423 ***
## ---
## Signif. codes:  0 '***' 0.001 '**' 0.01 '*' 0.05 '.' 0.1 ' ' 1
```

```
#compare final models with and w/o random slope:
AIC(fct1, fct2)#lower AIC with random slope
```

```
##      df       AIC
## fct1 10 -213.9282
## fct2  8 -207.1893
```

```
#now narrow colour space

nct1<-lmer(Capture_Time~poly(Generation,2)*Bgd+(Generation|Folder), data=CWSCREEN15_NARROW)
#summary(nct1)
#model does not converge so use model without random slope

#model without random slope
nct2<-lmer(Capture_Time~poly(Generation,2)*Bgd+(Generation|Folder), data=CWSCREEN15_NARROW) 
#test model assumptions
plot(nct2)
```

```
hist(resid(nct2))
```

```
#very poor, try log transform
nct2<-lmer(log(Capture_Time)~poly(Generation,2)*Bgd+(Generation|Folder), data=CWSCREEN15_NARROW) 
hist(resid(nct2))
```

```
plot(nct2)#better
```

```
nct2<-lmer(log(Capture_Time)~poly(Generation,2)*Bgd+(1|Folder), data=CWSCREEN15_NARROW)
nct2b<-lmer(log(Capture_Time)~poly(Generation,2)+Bgd+(1|Folder), data=CWSCREEN15_NARROW)
anova(nct2, nct2b)#can remove Bgd:Generation interaction
```

```
## Data: CWSCREEN15_NARROW
## Models:
## nct2b: log(Capture_Time) ~ poly(Generation, 2) + Bgd + (1 | Folder)
## nct2: log(Capture_Time) ~ poly(Generation, 2) * Bgd + (1 | Folder)
##       Df    AIC    BIC  logLik deviance Chisq Chi Df Pr(>Chisq)  
## nct2b  6 369.15 395.28 -178.57   357.15                          
## nct2   8 368.38 403.23 -176.19   352.38 4.762      2    0.09246 .
## ---
## Signif. codes:  0 '***' 0.001 '**' 0.01 '*' 0.05 '.' 0.1 ' ' 1
```

```
nct2c<-lmer(log(Capture_Time)~poly(Generation,2)+(1|Folder), data=CWSCREEN15_NARROW)
anova(nct2c, nct2b)#can remove Bgd
```

```
## Data: CWSCREEN15_NARROW
## Models:
## nct2c: log(Capture_Time) ~ poly(Generation, 2) + (1 | Folder)
## nct2b: log(Capture_Time) ~ poly(Generation, 2) + Bgd + (1 | Folder)
##       Df    AIC    BIC  logLik deviance  Chisq Chi Df Pr(>Chisq)  
## nct2c  5 370.97 392.75 -180.49   360.97                           
## nct2b  6 369.15 395.28 -178.57   357.15 3.8261      1    0.05046 .
## ---
## Signif. codes:  0 '***' 0.001 '**' 0.01 '*' 0.05 '.' 0.1 ' ' 1
```

```
nct2d<-lmer(log(Capture_Time)~1+(1|Folder), data=CWSCREEN15_NARROW)
anova(nct2c, nct2d)#can't remove Generation
```

```
## Data: CWSCREEN15_NARROW
## Models:
## nct2d: log(Capture_Time) ~ 1 + (1 | Folder)
## nct2c: log(Capture_Time) ~ poly(Generation, 2) + (1 | Folder)
##       Df    AIC    BIC  logLik deviance  Chisq Chi Df Pr(>Chisq)  
## nct2d  3 373.31 386.38 -183.66   367.31                           
## nct2c  5 370.97 392.75 -180.49   360.97 6.3402      2      0.042 *
## ---
## Signif. codes:  0 '***' 0.001 '**' 0.01 '*' 0.05 '.' 0.1 ' ' 1
```

```
#is the polynomial useful?
nct2c_2<-lmer(log(Capture_Time)~Generation+(1|Folder), data=CWSCREEN15_NARROW)
anova(nct2c, nct2c_2)#no significant effect of polynomial
```

```
## Data: CWSCREEN15_NARROW
## Models:
## nct2c_2: log(Capture_Time) ~ Generation + (1 | Folder)
## nct2c: log(Capture_Time) ~ poly(Generation, 2) + (1 | Folder)
##         Df    AIC    BIC  logLik deviance Chisq Chi Df Pr(>Chisq)
## nct2c_2  4 368.97 386.40 -180.49   360.97                        
## nct2c    5 370.97 392.75 -180.49   360.97     0      1     0.9983
```

```
nct2d_2<-lmer(log(Capture_Time)~1+(1|Folder), data=CWSCREEN15_NARROW)
anova(nct2c_2, nct2d_2)#can't remove Generation
```

```
## Data: CWSCREEN15_NARROW
## Models:
## nct2d_2: log(Capture_Time) ~ 1 + (1 | Folder)
## nct2c_2: log(Capture_Time) ~ Generation + (1 | Folder)
##         Df    AIC    BIC  logLik deviance  Chisq Chi Df Pr(>Chisq)  
## nct2d_2  3 373.31 386.38 -183.66   367.31                           
## nct2c_2  4 368.97 386.40 -180.49   360.97 6.3402      1     0.0118 *
## ---
## Signif. codes:  0 '***' 0.001 '**' 0.01 '*' 0.05 '.' 0.1 ' ' 1
```

##### Plot results

Figure 5: Changes in camouflage metrics in the screen-based experiments

##### Experimental versus control populations

Do the trajectories of experimental (selected) populations and control (neutral drift) replicates diverge across generations? Key variables are as above; SelectionType represents the scenario described in SI (control or experimental)

```
#load control population data
CWSCREEN_CONTROL<-read.csv("CamoWild_ScreenControl.csv", header=TRUE)
CWSCREEN_COLDIST_CONTROL<-read.csv("CamoWild_ScreenControl_ColDist.csv", header=TRUE)

#add useful variables
#colour difference between means - background to target (euclidean distance) 
CWSCREEN_CONTROL$eucdisTB<-sqrt((CWSCREEN_CONTROL$Target_L_Mean-CWSCREEN_CONTROL$BG_L_Mean)^2+
                                  (CWSCREEN_CONTROL$Target_A_Mean-CWSCREEN_CONTROL$BG_A_Mean)^2+
                                 (CWSCREEN_CONTROL$ Target_B_Mean-CWSCREEN_CONTROL$BG_B_Mean)^2)
#luminance difference (difference in L values)
CWSCREEN_CONTROL$LdiffTB<-abs(CWSCREEN_CONTROL$Target_L_Mean-CWSCREEN_CONTROL$BG_L_Mean)

#separate into full and narrow colour spaces
CWSCREEN_CONTROL_FULL<-CWSCREEN_CONTROL%>%
  filter(ColSpace=="Full")
CWSCREEN_CONTROL_NARROW<-CWSCREEN_CONTROL%>%
  filter(ColSpace=="Narrow")
CWSCREEN_COLDIST_CONTROL_FULL<-CWSCREEN_COLDIST_CONTROL%>%
  filter(ColSpace=="Full")
CWSCREEN_COLDIST_CONTROL_NARROW<-CWSCREEN_COLDIST_CONTROL%>%
  filter(ColSpace=="Narrow")

#combine experimental and control datasets

CWSCREEN15_FULL2<-CWSCREEN15_FULL%>%
  select(Generation,ID, ParticipantID,Folder,ColSpace,Bgd,SelectionType,eucdisTB,LdiffTB,Target_L_GabRat)
CWSCREEN15_NARROW2<-CWSCREEN15_NARROW%>%
  select(Generation,ID, ParticipantID,Folder,ColSpace,Bgd,SelectionType,eucdisTB,LdiffTB,Target_L_GabRat)
CWSCREEN_CONTROL_FULL2<-CWSCREEN_CONTROL_FULL%>%
  select(Generation,ID, ParticipantID,Folder,ColSpace,Bgd,SelectionType,eucdisTB,LdiffTB,Target_L_GabRat)
CWSCREEN_CONTROL_NARROW2<-CWSCREEN_CONTROL_NARROW%>%
  select(Generation,ID, ParticipantID,Folder,ColSpace,Bgd,SelectionType,eucdisTB,LdiffTB,Target_L_GabRat)

COMBINED_FULL<-rbind(CWSCREEN15_FULL2,CWSCREEN_CONTROL_FULL2)
COMBINED_NARROW<-rbind(CWSCREEN15_NARROW2,CWSCREEN_CONTROL_NARROW2)

#colour distance dataset

#restrict experimental datasets to 15 generations only
CWSCREEN15_COLDIST_FULL<-CWSCREEN_COLDIST_FULL%>%
  filter(Generation!=16)
CWSCREEN15_COLDIST_NARROW<-CWSCREEN_COLDIST_NARROW%>%
  filter(Generation!=16)

#add rounded colour distance and restrict to useful variaibles only
CWSCREEN_COLDIST_CONTROL_FULL$ColDistRound<-round(CWSCREEN_COLDIST_CONTROL_FULL$NewColMap_DeltaS)
CWSCREEN_COLDIST_CONTROL_NARROW$ColDistRound<-round(CWSCREEN_COLDIST_CONTROL_NARROW$NewColMap_DeltaS)
CWSCREEN_COLDIST_CONTROL_FULL<-CWSCREEN_COLDIST_CONTROL_FULL%>%
  select(ColSpace, Bgd, Folder, Generation, ID, NewColMap_DeltaS, ColDistRound, SelectionType)
CWSCREEN_COLDIST_CONTROL_NARROW<-CWSCREEN_COLDIST_CONTROL_NARROW%>%
  select(ColSpace, Bgd, Folder, Generation, ID, NewColMap_DeltaS, ColDistRound, SelectionType)
#combine datasets
COMBINED_COLDIST_FULL<-rbind(CWSCREEN15_COLDIST_FULL,CWSCREEN_COLDIST_CONTROL_FULL)
COMBINED_COLDIST_NARROW<-rbind(CWSCREEN15_COLDIST_NARROW,CWSCREEN_COLDIST_CONTROL_NARROW)
```

###### Trends in luminance difference

Full colour space:

```
combimodl1<-lmer(sqrt(LdiffTB)~poly(Generation,2)*SelectionType+(Generation|Folder), data=COMBINED_FULL)
plot(combimodl1)
```

```
hist(resid(combimodl1))#sqrt transform is best
```

```
combimodl1b<-lmer(sqrt(LdiffTB)~poly(Generation,2)+SelectionType+(Generation|Folder), data=COMBINED_FULL)
anova(combimodl1, combimodl1b)#remove interaction - no difference between exptl and control runs
```

```
## Data: COMBINED_FULL
## Models:
## combimodl1b: sqrt(LdiffTB) ~ poly(Generation, 2) + SelectionType + (Generation | 
## combimodl1b:     Folder)
## combimodl1: sqrt(LdiffTB) ~ poly(Generation, 2) * SelectionType + (Generation | 
## combimodl1:     Folder)
##             Df   AIC   BIC  logLik deviance  Chisq Chi Df Pr(>Chisq)
## combimodl1b  8 11273 11321 -5628.6    11257                         
## combimodl1  10 11277 11336 -5628.3    11257 0.4851      2     0.7846
```

```
combimodl1c<-lmer(sqrt(LdiffTB)~poly(Generation,2)+(Generation|Folder), data=COMBINED_FULL)
anova(combimodl1c, combimodl1b)#remove selection type
```

```
## Data: COMBINED_FULL
## Models:
## combimodl1c: sqrt(LdiffTB) ~ poly(Generation, 2) + (Generation | Folder)
## combimodl1b: sqrt(LdiffTB) ~ poly(Generation, 2) + SelectionType + (Generation | 
## combimodl1b:     Folder)
##             Df   AIC   BIC  logLik deviance  Chisq Chi Df Pr(>Chisq)
## combimodl1c  7 11273 11315 -5629.7    11259                         
## combimodl1b  8 11273 11321 -5628.6    11257 2.2021      1     0.1378
```

```
combimodl1d<-lmer(sqrt(LdiffTB)~(Generation|Folder), data=COMBINED_FULL)
anova(combimodl1c, combimodl1d)#significant effect of generation
```

```
## Data: COMBINED_FULL
## Models:
## combimodl1d: sqrt(LdiffTB) ~ (Generation | Folder)
## combimodl1c: sqrt(LdiffTB) ~ poly(Generation, 2) + (Generation | Folder)
##             Df   AIC   BIC  logLik deviance  Chisq Chi Df Pr(>Chisq)  
## combimodl1d  5 11278 11308 -5633.9    11268                           
## combimodl1c  7 11273 11315 -5629.7    11259 8.5352      2    0.01402 *
## ---
## Signif. codes:  0 '***' 0.001 '**' 0.01 '*' 0.05 '.' 0.1 ' ' 1
```

```
#is the polynomial useful?
combimodl1e<-lmer(sqrt(LdiffTB)~Generation+(Generation|Folder), data=COMBINED_FULL)
anova(combimodl1c, combimodl1e)#yes
```

```
## Data: COMBINED_FULL
## Models:
## combimodl1e: sqrt(LdiffTB) ~ Generation + (Generation | Folder)
## combimodl1c: sqrt(LdiffTB) ~ poly(Generation, 2) + (Generation | Folder)
##             Df   AIC   BIC  logLik deviance  Chisq Chi Df Pr(>Chisq)   
## combimodl1e  6 11278 11314 -5633.2    11266                            
## combimodl1c  7 11273 11315 -5629.7    11259 6.9944      1   0.008177 **
## ---
## Signif. codes:  0 '***' 0.001 '**' 0.01 '*' 0.05 '.' 0.1 ' ' 1
```

```
#and random effects?
rand(combimodl1c)#yes
```

```
## ANOVA-like table for random-effects: Single term deletions
## 
## Model:
## sqrt(LdiffTB) ~ poly(Generation, 2) + (Generation | Folder)
##                                     npar  logLik   AIC   LRT Df Pr(>Chisq)    
## <none>                                 7 -5626.9 11268                        
## Generation in (Generation | Folder)    5 -5642.6 11295 31.47  2  1.467e-07 ***
## ---
## Signif. codes:  0 '***' 0.001 '**' 0.01 '*' 0.05 '.' 0.1 ' ' 1
```

```
#without random slope
combimodl1i<-lmer(sqrt(LdiffTB)~poly(Generation,2)*SelectionType+(1|Folder), data=COMBINED_FULL)
plot(combimodl1)
```

```
hist(resid(combimodl1))#ok
```

```
combimodl1ib<-lmer(sqrt(LdiffTB)~poly(Generation,2)+SelectionType+(1|Folder), data=COMBINED_FULL)
anova(combimodl1i, combimodl1ib)#remove interaction - no difference between exptl and random run
combimodl1ic<-lmer(sqrt(LdiffTB)~poly(Generation,2)+(1|Folder), data=COMBINED_FULL)
anova(combimodl1ic, combimodl1ib)#remove selection type
combimodl1id<-lmer(sqrt(LdiffTB)~(1|Folder), data=COMBINED_FULL)
anova(combimodl1ic, combimodl1id)#significant effect of generation
#polynomial useful?
combimodl1ie<-lmer(sqrt(LdiffTB)~Generation+(1|Folder), data=COMBINED_FULL)
anova(combimodl1ic, combimodl1ie)#yes
```

```
AIC(combimodl1c,combimodl1ic)#use model with random slope
```

```
##              df      AIC
## combimodl1c   7 11267.81
## combimodl1ic  5 11295.28
```

Narrow colour space:

```
combimodl2<-lmer(LdiffTB~poly(Generation,2)*SelectionType+(Generation|Folder), data=COMBINED_NARROW)
plot(combimodl2)
```

```
hist(resid(combimodl2))
```

```
combimodl2<-lmer(sqrt(LdiffTB)~poly(Generation,2)*SelectionType+(Generation|Folder), data=COMBINED_NARROW)
plot(combimodl2)
```

```
hist(resid(combimodl2))#better
```

```
combimodl2b<-lmer(sqrt(LdiffTB)~poly(Generation,2)+SelectionType+(Generation|Folder), data=COMBINED_NARROW)
anova(combimodl2, combimodl2b)#sig interaction - difference between exptl and random runs
```

```
## Data: COMBINED_NARROW
## Models:
## combimodl2b: sqrt(LdiffTB) ~ poly(Generation, 2) + SelectionType + (Generation | 
## combimodl2b:     Folder)
## combimodl2: sqrt(LdiffTB) ~ poly(Generation, 2) * SelectionType + (Generation | 
## combimodl2:     Folder)
##             Df   AIC   BIC  logLik deviance  Chisq Chi Df Pr(>Chisq)   
## combimodl2b  8 10384 10431 -5183.8    10368                            
## combimodl2  10 10378 10438 -5179.0    10358 9.6499      2   0.008027 **
## ---
## Signif. codes:  0 '***' 0.001 '**' 0.01 '*' 0.05 '.' 0.1 ' ' 1
```

```
combimodl2c<-lmer(sqrt(LdiffTB)~Generation*SelectionType+(Generation|Folder), data=COMBINED_NARROW)
anova(combimodl2c, combimodl2)#polynomial not needed
```

```
## Data: COMBINED_NARROW
## Models:
## combimodl2c: sqrt(LdiffTB) ~ Generation * SelectionType + (Generation | Folder)
## combimodl2: sqrt(LdiffTB) ~ poly(Generation, 2) * SelectionType + (Generation | 
## combimodl2:     Folder)
##             Df   AIC   BIC  logLik deviance  Chisq Chi Df Pr(>Chisq)
## combimodl2c  8 10375 10422 -5179.3    10359                         
## combimodl2  10 10378 10438 -5179.0    10358 0.5865      2     0.7458
```

```
combimodl2d<-lmer(sqrt(LdiffTB)~Generation+SelectionType+(Generation|Folder), data=COMBINED_NARROW)
anova(combimodl2c, combimodl2d)#interaction still needed
```

```
## Data: COMBINED_NARROW
## Models:
## combimodl2d: sqrt(LdiffTB) ~ Generation + SelectionType + (Generation | Folder)
## combimodl2c: sqrt(LdiffTB) ~ Generation * SelectionType + (Generation | Folder)
##             Df   AIC   BIC  logLik deviance  Chisq Chi Df Pr(>Chisq)   
## combimodl2d  7 10382 10424 -5184.0    10368                            
## combimodl2c  8 10375 10422 -5179.3    10359 9.4426      1    0.00212 **
## ---
## Signif. codes:  0 '***' 0.001 '**' 0.01 '*' 0.05 '.' 0.1 ' ' 1
```

```
rand(combimodl2c)#random effects useful
```

```
## ANOVA-like table for random-effects: Single term deletions
## 
## Model:
## sqrt(LdiffTB) ~ Generation + SelectionType + (Generation | Folder) + 
##     Generation:SelectionType
##                                     npar  logLik   AIC    LRT Df Pr(>Chisq)   
## <none>                                 8 -5189.8 10396                        
## Generation in (Generation | Folder)    6 -5195.7 10403 11.636  2   0.002973 **
## ---
## Signif. codes:  0 '***' 0.001 '**' 0.01 '*' 0.05 '.' 0.1 ' ' 1
```

```
#without random slope
combimodl2c_simple<-lmer(sqrt(LdiffTB)~poly(Generation,2)*SelectionType+(1|Folder), data=COMBINED_NARROW)
combimodl2d_simple<-lmer(sqrt(LdiffTB)~poly(Generation,2)+SelectionType+(1|Folder), data=COMBINED_NARROW)
anova(combimodl2c_simple,combimodl2d_simple)#needs interaction
combimodl2e_simple<-lmer(sqrt(LdiffTB)~Generation*SelectionType+(1|Folder), data=COMBINED_NARROW)
anova(combimodl2c_simple,combimodl2e_simple)#don't need poly
combimodl2f_simple<-lmer(sqrt(LdiffTB)~Generation+SelectionType+(1|Folder), data=COMBINED_NARROW)
anova(combimodl2e_simple,combimodl2f_simple)#still a significant interaction
```

```
AIC(combimodl2e_simple,combimodl2c)#random slope best
```

```
##                    df      AIC
## combimodl2e_simple  6 10403.34
## combimodl2c         8 10395.70
```

###### Trends in colour difference

Full colour space:

```
combimodc1<-lmer(sqrt(eucdisTB)~poly(Generation,2)*SelectionType+(Generation|Folder), data=COMBINED_FULL)
plot(combimodc1)
```

```
hist(resid(combimodc1))#ok
```

```
combimodc1b<-lmer(sqrt(eucdisTB)~poly(Generation,2)+SelectionType+(Generation|Folder), data=COMBINED_FULL)
anova(combimodc1, combimodc1b)#can remove interaction
```

```
## Data: COMBINED_FULL
## Models:
## combimodc1b: sqrt(eucdisTB) ~ poly(Generation, 2) + SelectionType + (Generation | 
## combimodc1b:     Folder)
## combimodc1: sqrt(eucdisTB) ~ poly(Generation, 2) * SelectionType + (Generation | 
## combimodc1:     Folder)
##             Df    AIC    BIC  logLik deviance  Chisq Chi Df Pr(>Chisq)
## combimodc1b  8 9861.9 9909.6 -4923.0   9845.9                         
## combimodc1  10 9864.2 9923.8 -4922.1   9844.2 1.7411      2     0.4187
```

```
combimodc1c<-lmer(sqrt(eucdisTB)~poly(Generation,2)+(Generation|Folder), data=COMBINED_FULL)
anova(combimodc1c, combimodc1b)#remove scenario
```

```
## Data: COMBINED_FULL
## Models:
## combimodc1c: sqrt(eucdisTB) ~ poly(Generation, 2) + (Generation | Folder)
## combimodc1b: sqrt(eucdisTB) ~ poly(Generation, 2) + SelectionType + (Generation | 
## combimodc1b:     Folder)
##             Df    AIC    BIC logLik deviance  Chisq Chi Df Pr(>Chisq)
## combimodc1c  7 9860.0 9901.7  -4923   9846.0                         
## combimodc1b  8 9861.9 9909.6  -4923   9845.9 0.0586      1     0.8087
```

```
combimodc1d<-lmer(sqrt(eucdisTB)~1+(Generation|Folder), data=COMBINED_FULL)
anova(combimodc1c, combimodc1d)#need generation
```

```
## Data: COMBINED_FULL
## Models:
## combimodc1d: sqrt(eucdisTB) ~ 1 + (Generation | Folder)
## combimodc1c: sqrt(eucdisTB) ~ poly(Generation, 2) + (Generation | Folder)
##             Df    AIC    BIC  logLik deviance  Chisq Chi Df Pr(>Chisq)  
## combimodc1d  5 9862.3 9892.1 -4926.2   9852.3                           
## combimodc1c  7 9860.0 9901.7 -4923.0   9846.0 6.3456      2    0.04189 *
## ---
## Signif. codes:  0 '***' 0.001 '**' 0.01 '*' 0.05 '.' 0.1 ' ' 1
```

```
combimodc1e<-lmer(sqrt(eucdisTB)~Generation+(Generation|Folder), data=COMBINED_FULL)
anova(combimodc1e, combimodc1c)#polynomial useful
```

```
## Data: COMBINED_FULL
## Models:
## combimodc1e: sqrt(eucdisTB) ~ Generation + (Generation | Folder)
## combimodc1c: sqrt(eucdisTB) ~ poly(Generation, 2) + (Generation | Folder)
##             Df  AIC    BIC logLik deviance  Chisq Chi Df Pr(>Chisq)  
## combimodc1e  6 9864 9899.8  -4926     9852                           
## combimodc1c  7 9860 9901.7  -4923     9846 6.0354      1    0.01402 *
## ---
## Signif. codes:  0 '***' 0.001 '**' 0.01 '*' 0.05 '.' 0.1 ' ' 1
```

```
#and random effects?
rand(combimodc1c)#yes
```

```
## ANOVA-like table for random-effects: Single term deletions
## 
## Model:
## sqrt(eucdisTB) ~ poly(Generation, 2) + (Generation | Folder)
##                                     npar  logLik    AIC    LRT Df Pr(>Chisq)
## <none>                                 7 -4920.9 9855.8                     
## Generation in (Generation | Folder)    5 -4951.8 9913.5 61.746  2  3.908e-14
##                                        
## <none>                                 
## Generation in (Generation | Folder) ***
## ---
## Signif. codes:  0 '***' 0.001 '**' 0.01 '*' 0.05 '.' 0.1 ' ' 1
```

```
#summary(combimodc1c)
```

```
#without random slope
combimodc1i<-lmer(sqrt(eucdisTB)~poly(Generation,2)*SelectionType+(1|Folder), data=COMBINED_FULL)
plot(combimodc1)
```

```
hist(resid(combimodc1))#ok
```

```
combimodc1bi<-lmer(sqrt(eucdisTB)~poly(Generation,2)+SelectionType+(1|Folder), data=COMBINED_FULL)
anova(combimodc1i, combimodc1bi)#can't remove interaction  
combimodc1ci<-lmer(sqrt(eucdisTB)~Generation*SelectionType+(1|Folder), data=COMBINED_FULL)
anova(combimodc1ci, combimodc1i)#polynomial useful
```

```
AIC(combimodc1c,combimodc1i)#use model with random slope
```

```
##             df      AIC
## combimodc1c  7 9855.766
## combimodc1i  8 9901.019
```

Narrow colour space:

```
combimodc2<-lmer(eucdisTB~poly(Generation,2)*SelectionType+(Generation|Folder), data=COMBINED_NARROW)
plot(combimodc2)
```

```
hist(resid(combimodc2))
```

```
combimodc2<-lmer(sqrt(eucdisTB)~poly(Generation,2)*SelectionType+(Generation|Folder), data=COMBINED_NARROW)
plot(combimodc2)
```

```
hist(resid(combimodc2))#better
```

```
combimodc2b<-lmer(sqrt(eucdisTB)~poly(Generation,2)+SelectionType+(Generation|Folder), data=COMBINED_NARROW)
anova(combimodc2, combimodc2b)#sig interaction - difference between exptl and random runs
```

```
## Data: COMBINED_NARROW
## Models:
## combimodc2b: sqrt(eucdisTB) ~ poly(Generation, 2) + SelectionType + (Generation | 
## combimodc2b:     Folder)
## combimodc2: sqrt(eucdisTB) ~ poly(Generation, 2) * SelectionType + (Generation | 
## combimodc2:     Folder)
##             Df    AIC    BIC  logLik deviance  Chisq Chi Df Pr(>Chisq)    
## combimodc2b  8 9141.1 9188.8 -4562.5   9125.1                             
## combimodc2  10 9128.1 9187.8 -4554.1   9108.1 16.968      2  0.0002067 ***
## ---
## Signif. codes:  0 '***' 0.001 '**' 0.01 '*' 0.05 '.' 0.1 ' ' 1
```

```
combimodc2c<-lmer(sqrt(eucdisTB)~Generation*SelectionType+(Generation|Folder), data=COMBINED_NARROW)
anova(combimodc2c, combimodc2)#polynomial not needed
```

```
## Data: COMBINED_NARROW
## Models:
## combimodc2c: sqrt(eucdisTB) ~ Generation * SelectionType + (Generation | Folder)
## combimodc2: sqrt(eucdisTB) ~ poly(Generation, 2) * SelectionType + (Generation | 
## combimodc2:     Folder)
##             Df    AIC    BIC  logLik deviance  Chisq Chi Df Pr(>Chisq)
## combimodc2c  8 9126.4 9174.1 -4555.2   9110.4                         
## combimodc2  10 9128.1 9187.8 -4554.1   9108.1 2.2505      2     0.3246
```

```
combimodc2d<-lmer(sqrt(eucdisTB)~Generation+SelectionType+(Generation|Folder), data=COMBINED_NARROW)
anova(combimodc2c, combimodc2d)#interaction still needed
```

```
## Data: COMBINED_NARROW
## Models:
## combimodc2d: sqrt(eucdisTB) ~ Generation + SelectionType + (Generation | Folder)
## combimodc2c: sqrt(eucdisTB) ~ Generation * SelectionType + (Generation | Folder)
##             Df    AIC    BIC  logLik deviance  Chisq Chi Df Pr(>Chisq)    
## combimodc2d  7 9139.9 9181.7 -4563.0   9125.9                             
## combimodc2c  8 9126.4 9174.1 -4555.2   9110.4 15.557      1  8.003e-05 ***
## ---
## Signif. codes:  0 '***' 0.001 '**' 0.01 '*' 0.05 '.' 0.1 ' ' 1
```

```
rand(combimodc2c)#random effects useful
```

```
## ANOVA-like table for random-effects: Single term deletions
## 
## Model:
## sqrt(eucdisTB) ~ Generation + SelectionType + (Generation | Folder) + 
##     Generation:SelectionType
##                                     npar  logLik    AIC  LRT Df Pr(>Chisq)   
## <none>                                 8 -4566.8 9149.6                      
## Generation in (Generation | Folder)    6 -4572.8 9157.7 12.1  2   0.002358 **
## ---
## Signif. codes:  0 '***' 0.001 '**' 0.01 '*' 0.05 '.' 0.1 ' ' 1
```

```
#without random slope
combimodc2i<-lmer(sqrt(eucdisTB)~poly(Generation,2)*SelectionType+(1|Folder), data=COMBINED_NARROW)
combimodc2ib<-lmer(sqrt(eucdisTB)~poly(Generation,2)+SelectionType+(1|Folder), data=COMBINED_NARROW)
anova(combimodc2i, combimodc2ib)#sig interaction - difference between exptl and random runs
combimodc2ic<-lmer(sqrt(eucdisTB)~Generation*SelectionType+(1|Folder), data=COMBINED_NARROW)
anova(combimodc2ic, combimodc2i)#polynomial not needed
combimodc2id<-lmer(sqrt(eucdisTB)~Generation+SelectionType+(1|Folder), data=COMBINED_NARROW)
anova(combimodc2ic, combimodc2id)
```

```
AIC(combimodc2c,combimodc2ic)#random slope best
```

```
##              df      AIC
## combimodc2c   8 9149.559
## combimodc2ic  6 9157.659
```

###### Trends in colour difference (weighted average method)

Full colour space:

```
#use regular model for full, negative binomial for narrow colour space
combimodc3<-lmer(NewColMap_DeltaS~poly(Generation,2)*SelectionType+(Generation|Folder), data=COMBINED_COLDIST_FULL)
plot(combimodc3)
```

```
hist(resid(combimodc3))#ok
```

```
combimodc3b<-lmer(NewColMap_DeltaS~poly(Generation,2)+SelectionType+(Generation|Folder), data=COMBINED_COLDIST_FULL)
anova(combimodc3, combimodc3b)#remove interaction - no difference between exptl and control run
```

```
## Data: COMBINED_COLDIST_FULL
## Models:
## combimodc3b: NewColMap_DeltaS ~ poly(Generation, 2) + SelectionType + (Generation | 
## combimodc3b:     Folder)
## combimodc3: NewColMap_DeltaS ~ poly(Generation, 2) * SelectionType + (Generation | 
## combimodc3:     Folder)
##             Df   AIC   BIC logLik deviance Chisq Chi Df Pr(>Chisq)  
## combimodc3b  8 30860 30911 -15422    30844                          
## combimodc3  10 30858 30923 -15419    30838 5.577      2    0.06151 .
## ---
## Signif. codes:  0 '***' 0.001 '**' 0.01 '*' 0.05 '.' 0.1 ' ' 1
```

```
combimodc3c<-lmer(NewColMap_DeltaS~poly(Generation,2)+(Generation|Folder), data=COMBINED_COLDIST_FULL)
anova(combimodc3c, combimodc3b)#remove selection type
```

```
## Data: COMBINED_COLDIST_FULL
## Models:
## combimodc3c: NewColMap_DeltaS ~ poly(Generation, 2) + (Generation | Folder)
## combimodc3b: NewColMap_DeltaS ~ poly(Generation, 2) + SelectionType + (Generation | 
## combimodc3b:     Folder)
##             Df   AIC   BIC logLik deviance  Chisq Chi Df Pr(>Chisq)
## combimodc3c  7 30858 30903 -15422    30844                         
## combimodc3b  8 30860 30911 -15422    30844 0.3392      1     0.5603
```

```
combimodc3d<-lmer(NewColMap_DeltaS~(Generation|Folder), data=COMBINED_COLDIST_FULL)
anova(combimodc3c, combimodc3d)#significant effect of generation
```

```
## Data: COMBINED_COLDIST_FULL
## Models:
## combimodc3d: NewColMap_DeltaS ~ (Generation | Folder)
## combimodc3c: NewColMap_DeltaS ~ poly(Generation, 2) + (Generation | Folder)
##             Df   AIC   BIC logLik deviance  Chisq Chi Df Pr(>Chisq)    
## combimodc3d  5 30869 30901 -15429    30859                             
## combimodc3c  7 30858 30903 -15422    30844 14.411      2  0.0007424 ***
## ---
## Signif. codes:  0 '***' 0.001 '**' 0.01 '*' 0.05 '.' 0.1 ' ' 1
```

```
#polynomial useful?
combimodc3e<-lmer(NewColMap_DeltaS~Generation+(Generation|Folder), data=COMBINED_COLDIST_FULL)
anova(combimodc3c, combimodc3e)#no
```

```
## Data: COMBINED_COLDIST_FULL
## Models:
## combimodc3e: NewColMap_DeltaS ~ Generation + (Generation | Folder)
## combimodc3c: NewColMap_DeltaS ~ poly(Generation, 2) + (Generation | Folder)
##             Df   AIC   BIC logLik deviance  Chisq Chi Df Pr(>Chisq)
## combimodc3e  6 30858 30897 -15423    30846                         
## combimodc3c  7 30858 30903 -15422    30844 2.1733      1     0.1404
```

```
combimodc3f<-lmer(NewColMap_DeltaS~(Generation|Folder), data=COMBINED_COLDIST_FULL)
anova(combimodc3f, combimodc3e)#generation still significant
```

```
## Data: COMBINED_COLDIST_FULL
## Models:
## combimodc3f: NewColMap_DeltaS ~ (Generation | Folder)
## combimodc3e: NewColMap_DeltaS ~ Generation + (Generation | Folder)
##             Df   AIC   BIC logLik deviance  Chisq Chi Df Pr(>Chisq)    
## combimodc3f  5 30869 30901 -15429    30859                             
## combimodc3e  6 30858 30897 -15423    30846 12.238      1  0.0004682 ***
## ---
## Signif. codes:  0 '***' 0.001 '**' 0.01 '*' 0.05 '.' 0.1 ' ' 1
```

```
#and random effects?
rand(combimodc3e)#yes
```

```
## ANOVA-like table for random-effects: Single term deletions
## 
## Model:
## NewColMap_DeltaS ~ Generation + (Generation | Folder)
##                                     npar logLik   AIC   LRT Df Pr(>Chisq)    
## <none>                                 6 -15424 30861                        
## Generation in (Generation | Folder)    4 -15469 30946 89.48  2  < 2.2e-16 ***
## ---
## Signif. codes:  0 '***' 0.001 '**' 0.01 '*' 0.05 '.' 0.1 ' ' 1
```

```
#summary(combimodc3e)
```

```
#without random slope
combimodc3_2<-lmer(NewColMap_DeltaS~poly(Generation,2)*SelectionType+(1|Folder), data=COMBINED_COLDIST_FULL)
combimodc3b_2<-lmer(NewColMap_DeltaS~poly(Generation,2)+SelectionType+(1|Folder), data=COMBINED_COLDIST_FULL)
anova(combimodc3_2, combimodc3b_2)#cannot remove interaction - difference between exptl and control run
combimodc3c_2<-lmer(NewColMap_DeltaS~Generation*SelectionType+(1|Folder), data=COMBINED_COLDIST_FULL)
anova(combimodc3_2, combimodc3c_2)#polynomial significant
```

```
AIC(combimodc3e,combimodc3_2)#use model with random slope
```

```
##              df      AIC
## combimodc3e   6 30860.87
## combimodc3_2  8 30909.01
```

Narrow colour space:

```
combimodc4<-glmmTMB(ColDistRound ~ poly(Generation,2)*SelectionType+(Generation|Folder), data=COMBINED_COLDIST_NARROW, family=nbinom1 )
#check assumptions
res<-simulateResiduals(combimodc4, plot=T)
```

```
#check dispersion
par(mfrow = c(1,2))
testDispersion(res)
```

```
## 
##  DHARMa nonparametric dispersion test via sd of residuals fitted vs.
##  simulated
## 
## data:  simulationOutput
## ratioObsSim = 0.91366, p-value = 0.344
## alternative hypothesis: two.sided
```

```
testZeroInflation(res)#no dispersion problems, no zero-inflation
```

```
## 
##  DHARMa zero-inflation test via comparison to expected zeros with
##  simulation under H0 = fitted model
## 
## data:  simulationOutput
## ratioObsSim = 1.0045, p-value = 1
## alternative hypothesis: two.sided
```

```
#model simplification
combimodc4b<-glmmTMB(ColDistRound ~ poly(Generation,2)+SelectionType+(Generation|Folder), data=COMBINED_COLDIST_NARROW, family=nbinom1 )
anova(combimodc4, combimodc4b)#interaction not significant
```

```
## Data: COMBINED_COLDIST_NARROW
## Models:
## combimodc4b: ColDistRound ~ poly(Generation, 2) + SelectionType + (Generation | , zi=~0, disp=~1
## combimodc4b:     Folder), zi=~0, disp=~1
## combimodc4: ColDistRound ~ poly(Generation, 2) * SelectionType + (Generation | , zi=~0, disp=~1
## combimodc4:     Folder), zi=~0, disp=~1
##             Df   AIC   BIC  logLik deviance  Chisq Chi Df Pr(>Chisq)
## combimodc4b  8 10408 10460 -5196.1    10392                         
## combimodc4  10 10410 10474 -5195.0    10390 2.2144      2     0.3305
```

```
combimodc4c<-glmmTMB(ColDistRound ~ poly(Generation,2)+(Generation|Folder), data=COMBINED_COLDIST_NARROW, family=nbinom1 )
anova(combimodc4c, combimodc4b)#need selection type
```

```
## Data: COMBINED_COLDIST_NARROW
## Models:
## combimodc4c: ColDistRound ~ poly(Generation, 2) + (Generation | Folder), zi=~0, disp=~1
## combimodc4b: ColDistRound ~ poly(Generation, 2) + SelectionType + (Generation | , zi=~0, disp=~1
## combimodc4b:     Folder), zi=~0, disp=~1
##             Df   AIC   BIC  logLik deviance Chisq Chi Df Pr(>Chisq)   
## combimodc4c  7 10416 10461 -5200.8    10402                           
## combimodc4b  8 10408 10460 -5196.1    10392 9.313      1   0.002275 **
## ---
## Signif. codes:  0 '***' 0.001 '**' 0.01 '*' 0.05 '.' 0.1 ' ' 1
```

```
combimodc4d<-glmmTMB(ColDistRound ~ SelectionType+(Generation|Folder), data=COMBINED_COLDIST_NARROW, family=nbinom1 )
anova(combimodc4d, combimodc4b)#can remove generation
```

```
## Data: COMBINED_COLDIST_NARROW
## Models:
## combimodc4d: ColDistRound ~ SelectionType + (Generation | Folder), zi=~0, disp=~1
## combimodc4b: ColDistRound ~ poly(Generation, 2) + SelectionType + (Generation | , zi=~0, disp=~1
## combimodc4b:     Folder), zi=~0, disp=~1
##             Df   AIC   BIC  logLik deviance  Chisq Chi Df Pr(>Chisq)  
## combimodc4d  6 10409 10448 -5198.5    10397                           
## combimodc4b  8 10408 10460 -5196.1    10392 4.7841      2    0.09144 .
## ---
## Signif. codes:  0 '***' 0.001 '**' 0.01 '*' 0.05 '.' 0.1 ' ' 1
```

```
combimodc4e<-glmmTMB(ColDistRound ~ (Generation|Folder), data=COMBINED_COLDIST_NARROW, family=nbinom1 )
anova(combimodc4d, combimodc4e)#need selection type still
```

```
## Data: COMBINED_COLDIST_NARROW
## Models:
## combimodc4e: ColDistRound ~ (Generation | Folder), zi=~0, disp=~1
## combimodc4d: ColDistRound ~ SelectionType + (Generation | Folder), zi=~0, disp=~1
##             Df   AIC   BIC  logLik deviance Chisq Chi Df Pr(>Chisq)   
## combimodc4e  5 10416 10448 -5203.2    10406                           
## combimodc4d  6 10409 10448 -5198.5    10397 9.352      1   0.002227 **
## ---
## Signif. codes:  0 '***' 0.001 '**' 0.01 '*' 0.05 '.' 0.1 ' ' 1
```

```
#without random slope
combimodc4_2<-glmmTMB(ColDistRound ~ poly(Generation,2)*SelectionType+(1|Folder), data=COMBINED_COLDIST_NARROW, family=nbinom1 )
combimodc4b_2<-glmmTMB(ColDistRound ~ poly(Generation,2)+SelectionType+(1|Folder), data=COMBINED_COLDIST_NARROW, family=nbinom1 )
anova(combimodc4_2, combimodc4b_2)#interaction not significant 
combimodc4c_2<-glmmTMB(ColDistRound ~ poly(Generation,2)+(1|Folder), data=COMBINED_COLDIST_NARROW, family=nbinom1 )
anova(combimodc4c_2, combimodc4b_2)#scenario not needed
combimodc4d_2<-glmmTMB(ColDistRound ~ 1+(1|Folder), data=COMBINED_COLDIST_NARROW, family=nbinom1 )
anova(combimodc4d_2, combimodc4c_2)#need generation
combimodc4e_2<-glmmTMB(ColDistRound ~ Generation+(1|Folder), data=COMBINED_COLDIST_NARROW, family=nbinom1 )
anova(combimodc4c_2, combimodc4e_2)#don't need polynomial
combimodc4f_2<-glmmTMB(ColDistRound ~ 1+(1|Folder), data=COMBINED_COLDIST_NARROW, family=nbinom1 )
anova(combimodc4f_2, combimodc4e_2)#generation still sig
```

```
AIC(combimodc4e_2,combimodc4d)#random slope best
```

```
##               df      AIC
## combimodc4e_2  4 10592.34
## combimodc4d    6 10408.99
```

###### Edge disruption

Full colour space:

```
combimodg1<-lmer(Target_L_GabRat~poly(Generation,2)*SelectionType+(Generation|Folder), data=COMBINED_FULL)
#model convergence issues
```

```
#without random slope
combimodg1i<-lmer(Target_L_GabRat~poly(Generation,2)*SelectionType+(1|Folder), data=COMBINED_FULL)
combimodg1ib<-lmer(Target_L_GabRat~poly(Generation,2)+SelectionType+(1|Folder), data=COMBINED_FULL)
anova(combimodg1i, combimodg1ib)#remove interaction - no difference between exptl and control run
```

```
## Data: COMBINED_FULL
## Models:
## combimodg1ib: Target_L_GabRat ~ poly(Generation, 2) + SelectionType + (1 | 
## combimodg1ib:     Folder)
## combimodg1i: Target_L_GabRat ~ poly(Generation, 2) * SelectionType + (1 | 
## combimodg1i:     Folder)
##              Df     AIC     BIC logLik deviance  Chisq Chi Df Pr(>Chisq)
## combimodg1ib  6 -6182.9 -6147.1 3097.4  -6194.9                         
## combimodg1i   8 -6179.3 -6131.6 3097.6  -6195.3 0.4262      2     0.8081
```

```
combimodg1ic<-lmer(Target_L_GabRat~poly(Generation,2)+(1|Folder), data=COMBINED_FULL)
anova(combimodg1ic, combimodg1ib)#remove selection type
```

```
## Data: COMBINED_FULL
## Models:
## combimodg1ic: Target_L_GabRat ~ poly(Generation, 2) + (1 | Folder)
## combimodg1ib: Target_L_GabRat ~ poly(Generation, 2) + SelectionType + (1 | 
## combimodg1ib:     Folder)
##              Df     AIC     BIC logLik deviance  Chisq Chi Df Pr(>Chisq)
## combimodg1ic  5 -6182.4 -6152.5 3096.2  -6192.4                         
## combimodg1ib  6 -6182.9 -6147.1 3097.4  -6194.9 2.4874      1     0.1148
```

```
combimodg1id<-lmer(Target_L_GabRat~(1|Folder), data=COMBINED_FULL)
anova(combimodg1ic, combimodg1id)# no effect of generation
```

```
## Data: COMBINED_FULL
## Models:
## combimodg1id: Target_L_GabRat ~ (1 | Folder)
## combimodg1ic: Target_L_GabRat ~ poly(Generation, 2) + (1 | Folder)
##              Df     AIC     BIC logLik deviance  Chisq Chi Df Pr(>Chisq)
## combimodg1id  3 -6182.5 -6164.6 3094.3  -6188.5                         
## combimodg1ic  5 -6182.4 -6152.5 3096.2  -6192.4 3.8486      2      0.146
```

Narrow colour space:

```
combimodg2<-lmer(Target_L_GabRat~poly(Generation,2)*SelectionType+(Generation|Folder), data=COMBINED_NARROW)
plot(combimodg2)
```

```
hist(resid(combimodg2))
```

```
#model convergence issues during simplification, so use model without random slope
```

```
#without random slope
combimodg2_2<-lmer(Target_L_GabRat~poly(Generation,2)*SelectionType+(1|Folder), data=COMBINED_NARROW)
plot(combimodg2_2)
```

```
hist(resid(combimodg2_2))
```

```
combimodg2b_2<-lmer(Target_L_GabRat~poly(Generation,2)+SelectionType+(1|Folder), data=COMBINED_NARROW)
anova(combimodg2_2, combimodg2b_2)#no significant  interaction - no difference between exptl and control runs
```

```
## Data: COMBINED_NARROW
## Models:
## combimodg2b_2: Target_L_GabRat ~ poly(Generation, 2) + SelectionType + (1 | 
## combimodg2b_2:     Folder)
## combimodg2_2: Target_L_GabRat ~ poly(Generation, 2) * SelectionType + (1 | 
## combimodg2_2:     Folder)
##               Df     AIC     BIC logLik deviance  Chisq Chi Df Pr(>Chisq)
## combimodg2b_2  6 -7053.0 -7017.3 3532.5  -7065.0                         
## combimodg2_2   8 -7049.4 -7001.7 3532.7  -7065.4 0.3431      2     0.8423
```

```
combimodg2c_2<-lmer(Target_L_GabRat~poly(Generation,2)+(1|Folder), data=COMBINED_NARROW)
anova(combimodg2c_2, combimodg2b_2)#significant effect of selection type, but not interacting
```

```
## Data: COMBINED_NARROW
## Models:
## combimodg2c_2: Target_L_GabRat ~ poly(Generation, 2) + (1 | Folder)
## combimodg2b_2: Target_L_GabRat ~ poly(Generation, 2) + SelectionType + (1 | 
## combimodg2b_2:     Folder)
##               Df     AIC     BIC logLik deviance Chisq Chi Df Pr(>Chisq)   
## combimodg2c_2  5 -7046.7 -7016.9 3528.4  -7056.7                           
## combimodg2b_2  6 -7053.0 -7017.3 3532.5  -7065.0 8.338      1   0.003882 **
## ---
## Signif. codes:  0 '***' 0.001 '**' 0.01 '*' 0.05 '.' 0.1 ' ' 1
```

```
combimodg2d_2<-lmer(Target_L_GabRat~SelectionType+(1|Folder), data=COMBINED_NARROW)
anova(combimodg2b_2, combimodg2d_2)#generation significant
```

```
## Data: COMBINED_NARROW
## Models:
## combimodg2d_2: Target_L_GabRat ~ SelectionType + (1 | Folder)
## combimodg2b_2: Target_L_GabRat ~ poly(Generation, 2) + SelectionType + (1 | 
## combimodg2b_2:     Folder)
##               Df   AIC     BIC logLik deviance Chisq Chi Df Pr(>Chisq)  
## combimodg2d_2  4 -7050 -7026.1 3529.0    -7058                          
## combimodg2b_2  6 -7053 -7017.3 3532.5    -7065 7.064      2    0.02925 *
## ---
## Signif. codes:  0 '***' 0.001 '**' 0.01 '*' 0.05 '.' 0.1 ' ' 1
```

```
combimodg2e_2<-lmer(Target_L_GabRat~Generation+SelectionType+(1|Folder), data=COMBINED_NARROW)
anova(combimodg2e_2, combimodg2b_2)#polynomial not needed
```

```
## Data: COMBINED_NARROW
## Models:
## combimodg2e_2: Target_L_GabRat ~ Generation + SelectionType + (1 | Folder)
## combimodg2b_2: Target_L_GabRat ~ poly(Generation, 2) + SelectionType + (1 | 
## combimodg2b_2:     Folder)
##               Df     AIC     BIC logLik deviance  Chisq Chi Df Pr(>Chisq)
## combimodg2e_2  5 -7054.3 -7024.4 3532.1  -7064.3                         
## combimodg2b_2  6 -7053.0 -7017.3 3532.5  -7065.0 0.7772      1      0.378
```

```
#summary(combimodg2e_2)
combimodg2f_2<-lmer(Target_L_GabRat~SelectionType+(1|Folder), data=COMBINED_NARROW)
anova(combimodg2e_2, combimodg2f_2)#generation still significant
```

```
## Data: COMBINED_NARROW
## Models:
## combimodg2f_2: Target_L_GabRat ~ SelectionType + (1 | Folder)
## combimodg2e_2: Target_L_GabRat ~ Generation + SelectionType + (1 | Folder)
##               Df     AIC     BIC logLik deviance  Chisq Chi Df Pr(>Chisq)  
## combimodg2f_2  4 -7050.0 -7026.1 3529.0  -7058.0                           
## combimodg2e_2  5 -7054.3 -7024.4 3532.1  -7064.3 6.2868      1    0.01216 *
## ---
## Signif. codes:  0 '***' 0.001 '**' 0.01 '*' 0.05 '.' 0.1 ' ' 1
```

```
combimodg2g_2<-lmer(Target_L_GabRat~Generation+(1|Folder), data=COMBINED_NARROW)
anova(combimodg2e_2, combimodg2g_2)#selection type still significant
```

```
## Data: COMBINED_NARROW
## Models:
## combimodg2g_2: Target_L_GabRat ~ Generation + (1 | Folder)
## combimodg2e_2: Target_L_GabRat ~ Generation + SelectionType + (1 | Folder)
##               Df     AIC     BIC logLik deviance  Chisq Chi Df Pr(>Chisq)   
## combimodg2g_2  4 -7047.9 -7024.1 3528.0  -7055.9                            
## combimodg2e_2  5 -7054.3 -7024.4 3532.1  -7064.3 8.3376      1   0.003883 **
## ---
## Signif. codes:  0 '***' 0.001 '**' 0.01 '*' 0.05 '.' 0.1 ' ' 1
```

###### Plot results

Supplementary Figure 6: Changes in camouflage metrics in the screen-based experiments, experimental and control populations
