## Supporting data and code for "Adapting genetic algorithms for artificial evolution of visual patterns under selection from wild predators": CamoWild_Markdown.Rmd

````
---
title: "CamoEvoField_Markdown"
author: "E.S.Briolat"
date: "27th October 2023"
output: html_document
---

```{r setup, include=FALSE, warning =FALSE, message=FALSE}
knitr::opts_chunk$set(echo = TRUE)

library(tidyverse)
library(lme4)
library(lmerTest)
library(LMERConvenienceFunctions)
library(DHARMa)
library(broom)
library(glmmTMB)
library(ggeffects)
library(viridis)
library(knitr)
library(patchwork)
```

## Field trials
```{r prep_data_1, message=FALSE, warning=FALSE}
#load data
R1ALL<-read.csv("CamoField_Run1.csv", header=TRUE)
  #data for prey patterns in run1, from both smooth and furrowed populations
R2ALL<-read.csv("CamoField_Run2.csv", header=TRUE)
  #data for prey patterns in run2, from both smooth and furrowed populations

R1ALL_GR<-read.csv("CamoField_GabRat_Run1.csv", header=TRUE)
R2ALL_GR<-read.csv("CamoField_GabRat_Run2.csv", header=TRUE)
  #GabRat data for prey patterns in runs  and 2

#prepare data
R1ALL$Treatment <- relevel(R1ALL$Treatment, "Smooth")
R2ALL$Treatment <- relevel(R2ALL$Treatment, "Smooth")
R1ALL_GR$Treatment<-relevel(R1ALL_GR$Treatment, "Smooth")
R2ALL_GR$Treatment<-relevel(R2ALL_GR$Treatment, "Smooth")

#plotting theme
my_theme<-theme(panel.grid.major = element_blank(), panel.grid.minor = element_blank(),
                panel.background = element_blank(), axis.line = element_line(colour = "black"))

```

### Field trials - how do camouflage metrics change across generations?

Linear models test the interacting effects of generation and background type (treatment). 
Generation is initially fitted as a 2nd order polynomial, and the significance of the polynomial term is tested in the final model.
  
#### Run 1

Colour difference between mean colours (prey against gravel backgrounds):

```{r colour_difference, message=FALSE, warning=FALSE, fig.show='hide'}

r1mod1<-lm(dS~poly(Generation,2)*Treatment, data=R1ALL)
par(mfrow=c(2,2))
plot(r1mod1)
hist(resid(r1mod1))

#model simplification
r1mod1b<-lm(dS~poly(Generation,2)+Treatment, data=R1ALL)
anova(r1mod1b, r1mod1)#significant interaction

#test effect of polynomial
r1mod1c<-lm(dS~Generation*Treatment, data=R1ALL)
anova(r1mod1,r1mod1c)#polynomial is useful

summary(r1mod1)

#plot

#make predictions
pred<-predict(r1mod1, newdata = R1ALL, interval = 'confidence')
R1ALLp<-cbind(R1ALL, pred)

#make plot
rep1col<-ggplot(R1ALLp, aes(Generation, dS, col=Treatment,linetype=Treatment, fill=Treatment))+
  geom_point(alpha=0.3)+
  scale_colour_manual(values=c("black","grey50"))+
  scale_fill_manual(values=c("black","grey50"))+
  facet_grid(.~Treatment)+
  scale_x_continuous(breaks=c(0,5,10,15))+
  geom_ribbon(aes(ymin=lwr,ymax=upr), alpha=0.2, colour=NA)+
  geom_line(aes(Generation, fit), size=1)+
  guides(col=FALSE)+guides(linetype=FALSE)+guides(fill=FALSE)+
  ylab("Colour difference between means (Delta-S)")+
  my_theme+
  theme(axis.title = element_text(size = 16))  +
  theme(axis.text = element_text(size = 14)) +
  theme(strip.background = element_rect( fill="grey90"))+
  theme(strip.text.x = element_blank())
```

Colour difference based on weighted average distance:

```{r colour_difference_newmethod, message=FALSE, warning=FALSE, fig.show='hide'}

r1mod1d<-lm(NewMapComp_DeltaS~poly(Generation,2)*Treatment, data=R1ALL)
par(mfrow=c(2,2))
plot(r1mod1d)
hist(resid(r1mod1d))

#model simplification
r1mod1e<-lm(NewMapComp_DeltaS~poly(Generation,2)+Treatment, data=R1ALL)
anova(r1mod1d, r1mod1e)#significant interaction

#test effect of polynomial
r1mod1f<-lm(NewMapComp_DeltaS~Generation*Treatment, data=R1ALL)
anova(r1mod1d,r1mod1f)#polynomial is useful

summary(r1mod1d)

#plot

#make predictions
pred<-predict(r1mod1d, newdata = R1ALL, interval = 'confidence')
R1ALLp2<-cbind(R1ALL, pred)

#make plot
rep1colnew<-ggplot(R1ALLp2, aes(Generation, NewMapComp_DeltaS, col=Treatment,linetype=Treatment, fill=Treatment))+
  geom_point(alpha=0.3)+
  scale_colour_manual(values=c("black","grey50"))+
  scale_fill_manual(values=c("black","grey50"))+
  facet_grid(.~Treatment)+
  scale_x_continuous(breaks=c(0,5,10,15))+
  geom_ribbon(aes(ymin=lwr,ymax=upr), alpha=0.2, colour=NA)+
  geom_line(aes(Generation, fit), size=1)+
  guides(col=FALSE)+guides(linetype=FALSE)+guides(fill=FALSE)+
  ylab("Weighted average colour difference (Delta-S)")+
  my_theme+
  theme(axis.title = element_text(size = 16))  +
  theme(axis.text = element_text(size = 14)) +
  theme(strip.background = element_rect( fill="grey90"))+
  theme(strip.text.x = element_blank())
```


Luminance difference:

```{r luminance_difference, message=FALSE, warning=FALSE, fig.show='hide'}

r1mod2<-lm(dL~poly(Generation,2)*Treatment, data=R1ALL)
par(mfrow=c(2,2))
plot(r1mod2)
hist(resid(r1mod2))
#best transform is square root
r1mod2<-lm(sqrt(dL)~poly(Generation,2)*Treatment, data=R1ALL)
par(mfrow=c(2,2))
plot(r1mod2)
hist(resid(r1mod2))

#model simplification
r1mod2b<-lm(sqrt(dL)~poly(Generation,2)+Treatment, data=R1ALL)
anova(r1mod2, r1mod2b)#significant interaction
#is polynomial useful?
r1mod2c<-lm(sqrt(dL)~Generation*Treatment, data=R1ALL)
anova(r1mod2,r1mod2c)#yes

summary(r1mod2)

#make predictions
pred2<-predict(r1mod2, newdata = R1ALL, interval = 'confidence')
R1ALL$sqrtdL<-sqrt(R1ALL$dL)
R1ALLp2<-cbind(R1ALL, pred2)

rep1lum<-ggplot(R1ALLp2, aes(Generation, sqrtdL, col=Treatment,linetype=Treatment, fill=Treatment))+
  geom_point(alpha=0.3)+
  scale_colour_manual(values=c("black","grey50"))+
  scale_fill_manual(values=c("black","grey50"))+
  facet_grid(.~Treatment)+
  scale_x_continuous(breaks=c(0,5,10,15))+
  geom_ribbon(aes(ymin=lwr,ymax=upr), alpha=0.2, colour=NA)+
  geom_line(aes(Generation, fit), size=1)+
  guides(col=FALSE)+guides(linetype=FALSE)+guides(fill=FALSE)+
  ylab("Luminance difference (sqrt(Delta-L))")+
  my_theme+
  theme(axis.title = element_text(size = 16))  +
  theme(axis.text = element_text(size = 14)) +
  theme(strip.background = element_rect( fill="grey90"))+
  theme(strip.text.x = element_blank())
```

Pattern difference:

```{r pattern_difference, message=FALSE, warning=FALSE, fig.show='hide'}
r1mod3<-lm(pattern_energy_difference~poly(Generation,2)*Treatment, data=R1ALL)
par(mfrow=c(2,2))
plot(r1mod3)
hist(resid(r1mod3))
#try a log transform
r1mod3<-lm(log(pattern_energy_difference)~poly(Generation,2)*Treatment, data=R1ALL)
par(mfrow=c(2,2))
plot(r1mod3)
hist(resid(r1mod3))#better

#model simplification
r1mod3b<-lm(log(pattern_energy_difference)~poly(Generation,2)+Treatment, data=R1ALL)
anova(r1mod3, r1mod3b)#significant interaction
#is polynomial useful?
r1mod3c<-lm(log(pattern_energy_difference)~Generation*Treatment, data=R1ALL)
anova(r1mod3c, r1mod3)#polynomial not significant
#further simplification
r1mod3d<-lm(log(pattern_energy_difference)~Generation+Treatment, data=R1ALL)
anova(r1mod3c, r1mod3d)#significant interaction

summary(r1mod3c)

#make predictions
pred3<-predict(r1mod3c, newdata = R1ALL, interval = 'confidence')
R1ALL$logpat<-log(R1ALL$pattern_energy_difference)
R1ALLp3<-cbind(R1ALL, pred3)

rep1pat<-ggplot(R1ALLp3, aes(Generation, logpat, col=Treatment,linetype=Treatment, fill=Treatment))+
  geom_point(alpha=0.3)+
  scale_colour_manual(values=c("black","grey50"))+
  scale_fill_manual(values=c("black","grey50"))+
  facet_grid(.~Treatment)+
  scale_x_continuous(breaks=c(0,5,10,15))+
  geom_ribbon(aes(ymin=lwr,ymax=upr), alpha=0.2, colour=NA)+
  geom_line(aes(Generation, fit), size=1)+
  guides(col=FALSE)+guides(linetype=FALSE)+guides(fill=FALSE)+
  ylab("Pattern difference (log(energy difference))")+
  my_theme+
  theme(axis.title = element_text(size = 16))  +
  theme(axis.text = element_text(size = 14)) +
  theme(strip.background = element_rect( fill="grey90"))+
  theme(strip.text.x = element_blank())
```

Edge disruption (GabRat):

```{r gabrat_difference, message=FALSE, warning=FALSE, fig.show='hide'}

r1mod4<-lm(GabRat~poly(Generation,2)*Treatment, data=R1ALL_GR)
par(mfrow=c(2,2))
plot(r1mod4)
hist(resid(r1mod4))
#log transform is better
r1mod4<-lm(log(GabRat)~poly(Generation,2)*Treatment, data=R1ALL_GR)
par(mfrow=c(2,2))
plot(r1mod4)
hist(resid(r1mod4))

#model simplification
r1mod4b<-lm(log(GabRat)~poly(Generation,2)+Treatment, data=R1ALL_GR)
anova(r1mod4, r1mod4b)#significant interaction
#test polynomial
r1mod4c<-lm(log(GabRat)~Generation*Treatment, data=R1ALL_GR)
anova(r1mod4,r1mod4c)#polynomial is useful

summary(r1mod4)

#make predictions
R1ALL_GR$loggr<-log(R1ALL_GR$GabRat)
r1mod4<-lm(loggr~poly(Generation,2)*Treatment, data=R1ALL_GR)

pred<-predict(r1mod4, newdata = R1ALL_GR, interval = 'confidence')
R1ALL_GRp<-cbind(R1ALL_GR, pred)

rep1gr<-ggplot(R1ALL_GRp, aes(Generation, loggr, col=Treatment,linetype=Treatment, fill=Treatment))+
  geom_point(alpha=0.3)+
  scale_colour_manual(values=c("black","grey50"))+
  scale_fill_manual(values=c("black","grey50"))+
  facet_grid(.~Treatment)+
  scale_x_continuous(breaks=c(0,5,10,15))+
  geom_ribbon(aes(ymin=lwr,ymax=upr), alpha=0.2, colour=NA)+
  geom_line(aes(Generation, fit), size=1)+
  guides(col=FALSE)+guides(linetype=FALSE)+guides(fill=FALSE)+
  ylab("Edge disruption (log(GabRat))")+
  my_theme+
  theme(axis.title = element_text(size = 16))  +
  theme(axis.text = element_text(size = 14)) +
  theme(strip.background = element_rect( fill="grey90"))+
  theme(strip.text.x = element_blank())
```
 

```{r summarise_run1, message=FALSE,warning= FALSE, echo=FALSE}

#prepare data
R1ALL011<-R1ALL%>%
  filter(Generation==0 | Generation==11 )
R1ALL011$Generation<-as.factor(R1ALL011$Generation)

#summarise
kable(R1ALL011%>%
  group_by(Generation, Treatment)%>%
  summarise_at(c("dS","NewMapComp_DeltaS", "dL"),mean),
  caption ="Table 1: Run 1 - mean colour and luminance contrast (prey-background) in first and final generations")

```

####Run 2

Colour difference between means:

```{r colour_difference_2, message=FALSE, warning=FALSE, fig.show='hide'}
r2mod1<-lm(dS~poly(Generation,2)*Treatment, data=R2ALL)
par(mfrow=c(2,2))
plot(r2mod1)
hist(resid(r2mod1))

#model simplification
r2mod1b<-lm(dS~poly(Generation,2)+Treatment, data=R2ALL)
anova(r2mod1,r2mod1b)#significant interaction

#is polynomial useful?
r2mod1c<-lm(dS~Generation*Treatment, data=R2ALL)
anova(r2mod1,r2mod1c)#yes

summary(r2mod1)

#make predictions
pred<-predict(r2mod1, newdata = R2ALL, interval = 'confidence')
R2ALLp<-cbind(R2ALL, pred)

rep2col<-ggplot(R2ALLp, aes(Generation, dS, col=Treatment,linetype=Treatment, fill=Treatment))+
  geom_point(alpha=0.3)+
  scale_colour_manual(values=c("darkorange3","darkorange"))+
  scale_fill_manual(values=c("darkorange3","darkorange"))+
  facet_grid(.~Treatment)+
  scale_x_continuous(breaks=c(0,5,10,15))+
  geom_ribbon(aes(ymin=lwr,ymax=upr), alpha=0.2, colour=NA)+
  geom_line(aes(Generation, fit), size=1)+
  guides(col=FALSE)+guides(linetype=FALSE)+guides(fill=FALSE)+
  ylab("Colour difference between means (Delta-S)")+
  my_theme+
  theme(axis.title = element_text(size = 16))  +
  theme(axis.text = element_text(size = 14)) +
  theme(strip.background = element_rect( fill="grey90"))+
  theme(strip.text.x = element_blank())

```
Weighted average colour difference with new method:

```{r colour_difference_newmethod2, message=FALSE, warning=FALSE, fig.show='hide'}
r2mod2<-lm(NewMapComp_DeltaS~poly(Generation,2)*Treatment, data=R2ALL)
par(mfrow=c(2,2))
plot(r2mod2)
hist(resid(r2mod2))

#model simplification
r2mod2b<-lm(NewMapComp_DeltaS~poly(Generation,2)+Treatment, data=R2ALL)
anova(r2mod2,r2mod2b)#significant interaction
#summary(r2mod2)

#is polynomial useful?
r2mod2c<-lm(NewMapComp_DeltaS~Generation*Treatment, data=R2ALL)
anova(r2mod2,r2mod2c)#no

#further simplification?
r2mod2d<-lm(NewMapComp_DeltaS~Generation+Treatment, data=R2ALL)
anova(r2mod2d,r2mod2c)#interaction still significant
summary(r2mod2c)

#make predictions
pred<-predict(r2mod2c, newdata = R2ALL, interval = 'confidence')
R2ALLp<-cbind(R2ALL, pred)

rep2colnew<-ggplot(R2ALLp, aes(Generation, NewMapComp_DeltaS, col=Treatment,linetype=Treatment, fill=Treatment))+
  geom_point(alpha=0.3)+
  scale_colour_manual(values=c("darkorange3","darkorange"))+
  scale_fill_manual(values=c("darkorange3","darkorange"))+
  facet_grid(.~Treatment)+
  scale_x_continuous(breaks=c(0,5,10,15))+
  geom_ribbon(aes(ymin=lwr,ymax=upr), alpha=0.2, colour=NA)+
  geom_line(aes(Generation, fit), size=1)+
  guides(col=FALSE)+guides(linetype=FALSE)+guides(fill=FALSE)+
  ylab("Weighted average colour difference (Delta-S)")+
  my_theme+
  theme(axis.title = element_text(size = 16))  +
  theme(axis.text = element_text(size = 14)) +
  theme(strip.background = element_rect( fill="grey90"))+
  theme(strip.text.x = element_blank())

```

Luminance difference:

```{r luminance_difference_2, message=FALSE, warning=FALSE, fig.show='hide'}
r2mod2<-lm(dL~poly(Generation,2)*Treatment, data=R2ALL)
par(mfrow=c(2,2))
plot(r2mod2)
hist(resid(r2mod2))

#model simplification
r2mod2b<-lm(dL~poly(Generation,2)+Treatment, data=R2ALL)
anova(r2mod2,r2mod2b)#significant interaction
#is polynomial useful?
r2mod2c<-lm(dL~Generation*Treatment, data=R2ALL)
anova(r2mod2,r2mod2c)#no
#further simplification?
r2mod2d<-lm(dL~Generation+Treatment, data=R2ALL)
anova(r2mod2d,r2mod2c)#interaction still significant
summary(r2mod2c)

#make predictions
pred<-predict(r2mod2c, newdata = R2ALL, interval = 'confidence')
R2ALLp<-cbind(R2ALL, pred)

rep2lum<-ggplot(R2ALLp, aes(Generation, dL, col=Treatment,linetype=Treatment, fill=Treatment))+
  geom_point(alpha=0.3)+
  scale_colour_manual(values=c("darkorange3","darkorange"))+
  scale_fill_manual(values=c("darkorange3","darkorange"))+
  facet_grid(.~Treatment)+
  scale_x_continuous(breaks=c(0,5,10,15))+
  geom_ribbon(aes(ymin=lwr,ymax=upr), alpha=0.2, colour=NA)+
  geom_line(aes(Generation, fit), size=1)+
  guides(col=FALSE)+guides(linetype=FALSE)+guides(fill=FALSE)+
  ylab("Luminance difference (Delta-L)")+
  my_theme+
  theme(axis.title = element_text(size = 16))  +
  theme(axis.text = element_text(size = 14)) +
  theme(strip.background = element_rect( fill="grey90"))+
  theme(strip.text.x = element_blank())

```

Pattern difference: 

```{r pattern_difference_2, message=FALSE, warning=FALSE, fig.show='hide'}
r2mod3<-lm(pattern_energy_difference~poly(Generation,2)*Treatment, data=R2ALL)
par(mfrow=c(2,2))
plot(r2mod3)
hist(resid(r2mod3))
#log transform is better
r2mod3<-lm(log(pattern_energy_difference)~poly(Generation,2)*Treatment, data=R2ALL)
par(mfrow=c(2,2))
plot(r2mod3)
hist(resid(r2mod3))

#model simplication
r2mod3b<-lm(log(pattern_energy_difference)~poly(Generation,2)+Treatment, data=R2ALL)
anova(r2mod3, r2mod3b)#interaction not significant
r2mod3c<-lm(log(pattern_energy_difference)~poly(Generation,2), data=R2ALL)
anova(r2mod3c, r2mod3b)#no significant difference between backgrounds
r2mod3d<-lm(log(pattern_energy_difference)~1, data=R2ALL)
anova(r2mod3c, r2mod3d) #no effect of generation either

#plot data only
rep2pat<-ggplot(R2ALL, aes(Generation, log(pattern_energy_difference), col=Treatment))+
  geom_point(alpha=0.4)+
  scale_colour_manual(values=c("darkorange3","darkorange"))+
  facet_grid(.~Treatment)+
  scale_x_continuous(breaks=c(0,5,10,15))+
  guides(col=FALSE)+guides(linetype=FALSE)+
  ylab("Pattern difference log(energy difference)")+
  my_theme+
  theme(axis.title = element_text(size = 16))  +
  theme(axis.text = element_text(size = 14)) +
  theme(strip.background = element_rect( fill="grey90"))+
  theme(strip.text.x = element_blank())

```

Edge disruption (GabRat):

```{r gabrat_2, message=FALSE, warning=FALSE, fig.show='hide'}
r2mod4<-lm(GabRat~poly(Generation,2)*Treatment, data=R2ALL_GR)
par(mfrow=c(2,2))
plot(r2mod4)
hist(resid(r2mod4))
#log transform is better
r2mod4<-lm(log(GabRat)~poly(Generation,2)*Treatment, data=R2ALL_GR)
par(mfrow=c(2,2))
plot(r2mod4)
hist(resid(r2mod4))

#model simplification
r2mod4b<-lm(log(GabRat)~poly(Generation,2)+Treatment, data=R2ALL_GR)
anova(r2mod4,r2mod4b)#not significant
r2mod4c<-lm(log(GabRat)~poly(Generation,2), data=R2ALL_GR)
anova(r2mod4c,r2mod4b)#background type also not significant
r2mod4d<-lm(log(GabRat)~1, data=R2ALL_GR)
anova(r2mod4d,r2mod4c)#significant effect of generation

#is polynomial useful?
r2mod4e<-lm(log(GabRat)~Generation, data=R2ALL_GR)
anova(r2mod4c,r2mod4e)#no

#check generation is still significant
anova(r2mod4e,r2mod4d)#yes
summary(r2mod4e)

#make predictions
R2ALL_GR$loggr<-log(R2ALL_GR$GabRat)
pred<-predict(r2mod4e, newdata = R2ALL_GR, interval = 'confidence')
R2GRp<-cbind(R2ALL_GR, pred)

rep2gr<-ggplot(R2GRp, aes(Generation, loggr, col=Treatment,linetype=Treatment, fill=Treatment))+
  geom_point(alpha=0.3)+
  scale_colour_manual(values=c("darkorange3","darkorange"))+
  scale_fill_manual(values=c("darkorange3","darkorange"))+
  facet_grid(.~Treatment)+
  scale_x_continuous(breaks=c(0,5,10,15))+
  geom_ribbon(aes(ymin=lwr,ymax=upr), alpha=0.2, colour=NA)+
  geom_line(aes(Generation, fit), size=1)+
  guides(col=FALSE)+guides(linetype=FALSE)+guides(fill=FALSE)+
  ylab("Edge disruption (log(GabRat))")+
  my_theme+
  theme(axis.title = element_text(size = 16))  +
  theme(axis.text = element_text(size = 14)) +
  theme(strip.background = element_rect( fill="grey90"))+
  theme(strip.text.x = element_blank())


```

```{r summarise_run2, message=FALSE,warning= FALSE, echo =FALSE}

#prepare data
R2ALL016<-R2ALL%>%
  filter(Generation==0 | Generation==16 )
R2ALL016$Generation<-as.factor(R2ALL016$Generation)

#summarise
kable(R2ALL016%>%
  group_by(Generation, Treatment)%>%
  summarise_at(c("dS","NewMapComp_DeltaS", "dL"),mean),
  caption ="Table 2: Run 2 - mean colour and luminance contrast (prey-background) in first and final generations")

```

#### Plots for field trials

```{r plot_field, message=FALSE,warning= FALSE, echo=FALSE, fig.dim=c(20,30),fig.cap= "Figure 4: Camouflage metrics across generations, in field trials"}

(rep1col+rep2col)/
  (rep1colnew+ rep2colnew)/
  (rep1lum+rep2lum)/
  (rep1pat+rep2pat)/
  (rep1gr+rep2gr)
```

## Screen-based replicates

### Load and prepare data

Output from CamoEvo, and additional pattern difference and colour distance (weighted average) analyses
  
Key variables in main dataset are: Bgd (Background - furrowed (Fr) or smooth (Sm)), ColSpace (colour space - narrow or full), Folder/Participant ID (identify individual populations/participants), Generation (integer) + all CamoEvo metrics and gene data.

```{r prep_data_2, message=FALSE}

CWSCREEN<-read.csv("CamoWild_ScreenData.csv", header=TRUE)
CWSCREEN_PATTERN<-read.csv("CamoWild_ScreenData_Pattern.csv", header=TRUE)
CWSCREEN_COLDIST<-read.csv("CamoWild_ScreenData_ColDist.csv", header=TRUE)

#colour difference between means - background to target (euclidean distance) 
CWSCREEN$eucdisTB<-sqrt((CWSCREEN$Target_L_Mean-CWSCREEN$BG_L_Mean)^2+(CWSCREEN$Target_A_Mean-CWSCREEN$BG_A_Mean)^2+
                          (CWSCREEN$Target_B_Mean-CWSCREEN$BG_B_Mean)^2)
#luminance difference (difference in L values)
CWSCREEN$LdiffTB<-abs(CWSCREEN$Target_L_Mean-CWSCREEN$BG_L_Mean)

#separate full and narrow colour spaces for subsequent analyses
CWSCREEN_FULL<-CWSCREEN%>%
  filter(ColSpace=="Full")
CWSCREEN_NARROW<-CWSCREEN%>%
  filter(ColSpace=="Narrow")

```

How do metrics of camouflage efficacy change across generations, in populations on different backgrounds & using different colour spaces?

### Camouflage metrics: luminance difference

```{r analysis_luminance_difference_1, message=FALSE, warning=FALSE,  fig.show='hide'}
#start with Generation (2nd order polynomial) and Bgd, interacting, 
#+ random effect of Folder(Population) with random slope for generation

#full colour space
fl1<-lmer(LdiffTB~poly(Generation,2)*Bgd+(Generation|Folder), data=CWSCREEN_FULL)
#summary(fl1)

#test model assumptions
plot(fl1)
hist(resid(fl1))#not bad, not improved by transforms

#simplify model
fl1b<-lmer(LdiffTB~poly(Generation,2)+Bgd+(Generation|Folder), data=CWSCREEN_FULL) 
anova(fl1,fl1b)#can remove interaction
#summary(fl1b)
fl1c<-lmer(LdiffTB~poly(Generation,2)+(Generation|Folder), data=CWSCREEN_FULL) 
anova(fl1c,fl1b)#remove bgd
fl1d<-lmer(LdiffTB~(Generation|Folder), data=CWSCREEN_FULL) 
anova(fl1c,fl1d)#remove generation

#test significance of random effects
rand(fl1d)

#without polynomial?
fl1_2<-lmer(LdiffTB~Generation*Bgd+(Generation|Folder), data=CWSCREEN_FULL) 
fl1_2b<-lmer(LdiffTB~Generation+Bgd+(Generation|Folder), data=CWSCREEN_FULL) 
anova(fl1_2b, fl1_2)#remove interaction
fl1_2c<-lmer(LdiffTB~Generation+(Generation|Folder), data=CWSCREEN_FULL) 
anova(fl1_2b, fl1_2c)#remove bgd
fl1_2d<-lmer(LdiffTB~(Generation|Folder), data=CWSCREEN_FULL) 
anova(fl1_2d, fl1_2c)#remove generation - back to earlier model

#model without random slope
fl2<-lmer(LdiffTB~poly(Generation,2)*Bgd+(1|Folder), data=CWSCREEN_FULL)
fl2b<-lmer(LdiffTB~poly(Generation,2)+Bgd+(1|Folder), data=CWSCREEN_FULL)
anova(fl2, fl2b)#remove Bgd:Generation interaction
fl2c<-lmer(LdiffTB~poly(Generation,2)+(1|Folder), data=CWSCREEN_FULL)
anova(fl2c, fl2b)
fl2d<-lmer(LdiffTB~(1|Folder), data=CWSCREEN_FULL)
anova(fl2c, fl2d)#remove - no effects

#without polynomial?
fl2_2<-lmer(LdiffTB~Generation*Bgd+(1|Folder), data=CWSCREEN_FULL)
fl2b_2<-lmer(LdiffTB~Generation+Bgd+(1|Folder), data=CWSCREEN_FULL)
anova(fl2_2, fl2b_2)#remove Bgd:Generation interaction
fl2c_2<-lmer(LdiffTB~Generation+(1|Folder), data=CWSCREEN_FULL)
anova(fl2c_2, fl2b_2)
fl2d_2<-lmer(LdiffTB~(1|Folder), data=CWSCREEN_FULL)
anova(fl2c_2, fl2d_2)#remove all

#compare final models with and w/o random slope:
AIC(fl1d, fl2d)#lower AIC with random slope

#narrow colour space

nl1<-lmer(LdiffTB~poly(Generation,2)*Bgd+(Generation|Folder), data=CWSCREEN_NARROW)
#test model assumptions
plot(nl1)
hist(resid(nl1))
nl1<-lmer(sqrt(LdiffTB)~poly(Generation,2)*Bgd+(Generation|Folder), data=CWSCREEN_NARROW)
#summary(nl1)
plot(nl1)
hist(resid(nl1))# better with transform

#simplify model
nl1b<-lmer(sqrt(LdiffTB)~poly(Generation,2)+Bgd+(Generation|Folder), data=CWSCREEN_NARROW) 
anova(nl1,nl1b)#can remove interaction
nl1c<-lmer(sqrt(LdiffTB)~poly(Generation,2)+(Generation|Folder), data=CWSCREEN_NARROW) 
anova(nl1c,nl1b)#can remove bgd
nl1d<-lmer(sqrt(LdiffTB)~(Generation|Folder), data=CWSCREEN_NARROW) 
anova(nl1c,nl1d)#sig effect of generation

#test significance of random effects
rand(nl1c)#random effects not significant

#is the polynomial useful?
nl1c_2<-lmer(sqrt(LdiffTB)~Generation+(Generation|Folder), data=CWSCREEN_NARROW) 
anova(nl1c_2, nl1c)#polynomial not needed
nl1c_2b<-lmer(sqrt(LdiffTB)~(Generation|Folder), data=CWSCREEN_NARROW) 
anova(nl1c_2b, nl1c_2)#generation still significant
rand(nl1c_2)

#model without random slope
nl2<-lmer(sqrt(LdiffTB)~poly(Generation,2)*Bgd+(1|Folder), data=CWSCREEN_NARROW)
nl2b<-lmer(sqrt(LdiffTB)~poly(Generation,2)+Bgd+(1|Folder), data=CWSCREEN_NARROW)
anova(nl2, nl2b)#remove Bgd:Generation interaction
nl2c<-lmer(sqrt(LdiffTB)~poly(Generation,2)+(1|Folder), data=CWSCREEN_NARROW)
anova(nl2c, nl2b)
nl2d<-lmer(sqrt(LdiffTB)~(1|Folder), data=CWSCREEN_NARROW)
anova(nl2c, nl2d)#generation still significant
rand(nl2c)#random effects significant

#is the polynomial useful?
nl2c_2<-lmer(sqrt(LdiffTB)~Generation+(1|Folder), data=CWSCREEN_NARROW)
anova(nl2c_2,nl2c)#polynomial not needed

nl2d_2<-lmer(sqrt(LdiffTB)~(1|Folder), data=CWSCREEN_NARROW)
anova(nl2d_2, nl2c_2)#generation still significant

#compare final models with and w/o random slope:
AIC(nl1c_2, nl2c_2)#lower AIC without random slope
```


### Camouflage metrics: colour difference

```{r analysis_colour_difference_means, message=FALSE,  warning=FALSE,  fig.show='hide'}
#start with Generation (2nd order polynomial) and Bgd, interacting 
#+ random effect of Folder(population) with random slope for generation

#full colour space
fc1<-lmer(eucdisTB~poly(Generation,2)*Bgd+(Generation|Folder), data=CWSCREEN_FULL)
#summary(fc1)

#test model assumptions
plot(fc1)
hist(resid(fc1))#ok

#simplify model
fc1b<-lmer(eucdisTB~poly(Generation,2)+Bgd+(Generation|Folder), data=CWSCREEN_FULL) 
anova(fc1,fc1b)#can remove interaction
fc1c<-lmer(eucdisTB~poly(Generation,2)+(Generation|Folder), data=CWSCREEN_FULL) 
anova(fc1c,fc1b)#remove bgd
fc1d<-lmer(eucdisTB~(Generation|Folder), data=CWSCREEN_FULL) 
anova(fc1c,fc1d)#can't remove generation
#summary(fc1c)

#test significance of random effects
rand(fc1d)

#is the polynomial useful?
fc1c_2<-lmer(eucdisTB~Generation+(Generation|Folder), data=CWSCREEN_FULL) 
anova(fc1c,fc1c_2)#polynomial useful

#model without random slope
fc2<-lmer(eucdisTB~poly(Generation,2)*Bgd+(1|Folder), data=CWSCREEN_FULL)
fc2b<-lmer(eucdisTB~poly(Generation,2)+Bgd+(1|Folder), data=CWSCREEN_FULL)
anova(fc2, fc2b)#remove Bgd:Generation interaction
fc2c<-lmer(eucdisTB~poly(Generation,2)+(1|Folder), data=CWSCREEN_FULL)
anova(fc2c, fc2b)#can remove background
fc2d<-lmer(eucdisTB~(1|Folder), data=CWSCREEN_FULL)
anova(fc2c, fc2d)#can't remove generation 

#is the polynomial useful?
fc2c_2<-lmer(eucdisTB~Generation+(1|Folder), data=CWSCREEN_FULL)
anova(fc2c_2, fc2c)#polynomial signficant

#compare final models with and w/o random slope:
AIC(fc1c, fc2c)#lower AIC with random slope

#narrow colour space
nc1<-lmer(eucdisTB~poly(Generation,2)*Bgd+(Generation|Folder), data=CWSCREEN_NARROW)
#summary(nc1)

#test model assumptions
plot(nc1)
hist(resid(nc1))
nc1<-lmer(sqrt(eucdisTB)~poly(Generation,2)*Bgd+(Generation|Folder), data=CWSCREEN_NARROW)
#summary(nc1)
plot(nc1)
hist(resid(nc1))#better with sqrt transform

#simplify model
nc1b<-lmer(sqrt(eucdisTB)~poly(Generation,2)+Bgd+(Generation|Folder), data=CWSCREEN_NARROW) 
anova(nc1,nc1b)#can remove interaction
nc1c<-lmer(sqrt(eucdisTB)~poly(Generation,2)+(Generation|Folder), data=CWSCREEN_NARROW) 
anova(nc1c,nc1b)#can remove Bgd
nc1d<-lmer(sqrt(eucdisTB)~(Generation|Folder), data=CWSCREEN_NARROW) 
anova(nc1c,nc1d)#significant effect of generation

#test significance of random effects
rand(nc1c)#random effects not significant

#is the polynomial useful?
nc1c_2<-lmer(sqrt(eucdisTB)~Generation+(Generation|Folder), data=CWSCREEN_NARROW) 
anova(nc1c_2, nc1c)#polynomial needed

#model without random slope
nc2<-lmer(sqrt(eucdisTB)~poly(Generation,2)*Bgd+(1|Folder), data=CWSCREEN_NARROW)
nc2b<-lmer(sqrt(eucdisTB)~poly(Generation,2)+Bgd+(1|Folder), data=CWSCREEN_NARROW)
anova(nc2, nc2b)#remove Bgd:Generation interaction
nc2c<-lmer(sqrt(eucdisTB)~poly(Generation,2)+(1|Folder), data=CWSCREEN_NARROW)
anova(nc2c, nc2b)#remove Bgd
nc2d<-lmer(sqrt(eucdisTB)~(1|Folder), data=CWSCREEN_NARROW)
anova(nc2c, nc2d)#Generation still significant

#is the polynomial useful?
nc2c_2<-lmer(sqrt(eucdisTB)~Generation+(1|Folder), data=CWSCREEN_NARROW)
anova(nc2c_2,nc2c)#yes

#compare final models with and w/o random slope:
AIC(nc1c, nc2c)#lower AIC without random slope
```

### Camouflage metrics: colour difference using weighted average method  

```{r analysis_colour_difference_coldist, message=FALSE, warning=FALSE,  fig.show='hide'}
#start with Generation (2nd order polynomial) and Bgd, interacting 
#+ random effect of Folder (population) with random slope for generation

#as above, but using weighted average method for assessing colour difference

#Normal linear mixed effects models are not appropriate here (very poor diagnostics) due to zero-inflation. In the narrow colour space populations, lots of targets are composed of colours that are all present in the background, leading to lots of 0s.

#Tried manually creating a hurdle-like model by separating the data into two parts - a binomial response (is zero or not?) and a continuous response (if not zero, then?) - but this leads to convergence issuesfor the binomial part.
#The simplest way to analyse these data is to round the colour difference values and apply functions used for count data. Using these data, the same models can be tested here as for all other variables, for the full colour space. Diagnostics are still poor for the narrow colour space (but no zero-inflation), so a negative binomial family is used. 

#create new variable, rounding the weighted average colour difference
CWSCREEN_COLDIST$ColDistRound<-round(CWSCREEN_COLDIST$NewColMap_DeltaS)

CWSCREEN_COLDIST_FULL<-CWSCREEN_COLDIST%>%
  filter(ColSpace=="Full")
CWSCREEN_COLDIST_NARROW<-CWSCREEN_COLDIST%>%
  filter(ColSpace=="Narrow")

#full colour space
fcc1<-lmer(NewColMap_DeltaS~poly(Generation,2)*Bgd+(Generation|Folder), data=CWSCREEN_COLDIST_FULL)
#summary(fcc1)

#test model assumptions
plot(fcc1)
hist(resid(fcc1))#ok

#simplify model
fcc1b<-lmer(NewColMap_DeltaS~poly(Generation,2)+Bgd+(Generation|Folder), data=CWSCREEN_COLDIST_FULL) 
anova(fcc1,fcc1b)#can't remove interaction
fcc1c<-lmer(NewColMap_DeltaS~Generation*Bgd+(Generation|Folder), data=CWSCREEN_COLDIST_FULL) 
anova(fcc1c,fcc1)#polynomial important

#test significance of random effects
rand(fcc1)

#model without random slope
fcc2<-lmer(NewColMap_DeltaS~poly(Generation,2)*Bgd+(1|Folder), data=CWSCREEN_COLDIST_FULL)
fcc2b<-lmer(NewColMap_DeltaS~poly(Generation,2)+Bgd+(1|Folder), data=CWSCREEN_COLDIST_FULL)
anova(fcc2, fcc2b)#can't remove Bgd:Generation interaction
fcc2c<-lmer(NewColMap_DeltaS~Generation*Bgd+(1|Folder), data=CWSCREEN_COLDIST_FULL)
anova(fcc2c, fcc2)#need polynomial

#compare final models with and w/o random slope:
AIC(fcc1, fcc2)#lower AIC with random slope

#narrow colour space

ncc1<-glmmTMB(ColDistRound ~ poly(Generation,2)*Bgd+(Generation|Folder), data=CWSCREEN_COLDIST_NARROW, family=nbinom1 )
#check assumptions
res<-simulateResiduals(ncc1, plot=T)

#check dispersion
par(mfrow = c(1,2))
testDispersion(res)
testZeroInflation(res)# no underdispersion, no zero-inflation

#model simplification
ncc1b<-glmmTMB(ColDistRound ~ poly(Generation,2)+Bgd+(Generation|Folder), data=CWSCREEN_COLDIST_NARROW, family=nbinom1 )
anova(ncc1, ncc1b)#interaction significant 
#summary(ncc1)

#is the polynomial useful?
ncc1_2<-glmmTMB(ColDistRound ~ Generation*Bgd+(Generation|Folder), data=CWSCREEN_COLDIST_NARROW, family=nbinom1 )
anova(ncc1_2, ncc1)#yes

#try model without random slope for generation
ncc2<-glmmTMB(ColDistRound ~ poly(Generation,2)*Bgd+(1|Folder), data=CWSCREEN_COLDIST_NARROW, family=nbinom1 )
#check assumptions
res<-simulateResiduals(ncc2, plot=T)
#check dispersion
par(mfrow = c(1,2))
testDispersion(res)
testZeroInflation(res)#no underdispersion, no zero-inflation

#model simplification
ncc2b<-glmmTMB(ColDistRound ~ poly(Generation,2)+Bgd+(1|Folder), data=CWSCREEN_COLDIST_NARROW, family=nbinom1 )
anova(ncc2, ncc2b)#interaction is significant 
#summary(ncc2)
#is polynomial useful?
ncc2_2<-glmmTMB(ColDistRound ~ Generation*Bgd+(1|Folder), data=CWSCREEN_COLDIST_NARROW, family=nbinom1 )
anova(ncc2, ncc2_2)#yes

#compare best models with and without random slope
AIC(ncc1,ncc2)

```


### Camouflage metrics: edge disruption

```{r analysis_gabrat_1, message=FALSE, warning=FALSE,  fig.show='hide'}
#start with Generation (2nd order polynomial) and Bgd, interacting 
#+ random effect of Folder(Population) with random slope for generation

#full
fgr1<-lmer(Target_L_GabRat~poly(Generation,2)*Bgd+(Generation|Folder), data=CWSCREEN_FULL)

#test model assumptions
res<-simulateResiduals(fgr1, plot=T)
par(mfrow = c(1,1))
hist(resid(fgr1))
plot(fgr1)

#model simplification
fgr1b<-lmer(Target_L_GabRat~poly(Generation,2)+Bgd+(Generation|Folder), data=CWSCREEN_FULL)
anova(fgr1, fgr1b)#remove interaction
fgr1c<-lmer(Target_L_GabRat~poly(Generation,2)+(Generation|Folder), data=CWSCREEN_FULL)
anova(fgr1c, fgr1b)#remove Bgd
fgr1d<-lmer(Target_L_GabRat~(Generation|Folder), data=CWSCREEN_FULL)
anova(fgr1c, fgr1d)#remove Generation

#model without random slope
fgr2<-lmer(Target_L_GabRat~poly(Generation,2)*Bgd+(1|Folder), data=CWSCREEN_FULL)
fgr2b<-lmer(Target_L_GabRat~poly(Generation,2)+Bgd+(1|Folder), data=CWSCREEN_FULL)
anova(fgr2, fgr2b)#remove interaction
fgr2c<-lmer(Target_L_GabRat~poly(Generation,2)+(1|Folder), data=CWSCREEN_FULL)
anova(fgr2c, fgr2b)#remove Bgd
fgr2d<-lmer(Target_L_GabRat~(1|Folder), data=CWSCREEN_FULL)
anova(fgr2c, fgr2d)#can't remove Generation
#summary(fgr2c)

#is polynomial useful?
fgr2c_2<-lmer(Target_L_GabRat~Generation+(1|Folder), data=CWSCREEN_FULL)
anova(fgr2c_2, fgr2c)#no
#further simplification
fgr2d_2<-lmer(Target_L_GabRat~(1|Folder), data=CWSCREEN_FULL)
anova(fgr2d_2, fgr2c_2)#still a significant effect of generation

#compare models with and w/o random effects
AIC(fgr2c_2, fgr1d)

#narrow
ngr1<-lmer(Target_L_GabRat~poly(Generation,2)*Bgd+(Generation|Folder), data=CWSCREEN_NARROW)
#model fails to converge - use model w/o slope

#model without random slope
ngr2<-lmer(Target_L_GabRat~poly(Generation,2)*Bgd+(1|Folder), data=CWSCREEN_NARROW)
hist(resid(ngr2))
plot(ngr2)

ngr2b<-lmer(Target_L_GabRat~poly(Generation,2)+Bgd+(1|Folder), data=CWSCREEN_NARROW)
anova(ngr2, ngr2b)#remove interaction
ngr2c<-lmer(Target_L_GabRat~poly(Generation,2)+(1|Folder), data=CWSCREEN_NARROW)
anova(ngr2c, ngr2b)#remove Bgd
ngr2d<-lmer(Target_L_GabRat~(1|Folder), data=CWSCREEN_NARROW)
anova(ngr2c, ngr2d)#can't remove Generation
#summary(ngr2c)

#is polynomial useful?
ngr2c_2<-lmer(Target_L_GabRat~Generation+(1|Folder), data=CWSCREEN_NARROW)
anova(ngr2c_2, ngr2c)#no
#further simplification
ngr2d_2<-lmer(Target_L_GabRat~(1|Folder), data=CWSCREEN_NARROW)
anova(ngr2d_2, ngr2c_2)#still a significant effect of generation

```

### Camouflage metrics: pattern difference

```{r analysis_pattern_difference_1, message=FALSE, warning=FALSE,  fig.show='hide'}
#start with Generation (2nd order polynomial) and Bgd, interacting 
#+ random effect of Folder(population) with random slope for generation

CWSCREEN_PATTERN_FULL<-CWSCREEN_PATTERN%>%
  filter(ColSpace=="Full")
CWSCREEN_PATTERN_NARROW<-CWSCREEN_PATTERN%>%
  filter(ColSpace=="Narrow")

#full colour space
fp1<-lmer(pattern_energy_diff~poly(Generation,2)*Bgd+(Generation|Folder), data=CWSCREEN_PATTERN_FULL)

#test model assumptions
res<-simulateResiduals(fp1, plot=T)
hist(resid(fp1))
plot(fp1)#ok - no transforms improve diagnostics 

#model simplification
fp1b<-lmer(pattern_energy_diff~poly(Generation,2)+Bgd+(Generation|Folder), data=CWSCREEN_PATTERN_FULL)
anova(fp1,fp1b)#remove interaction
fp1c<-lmer(pattern_energy_diff~poly(Generation,2)+(Generation|Folder), data=CWSCREEN_PATTERN_FULL)
anova(fp1c,fp1b)#remove Bgd
fp1d<-lmer(pattern_energy_diff~(Generation|Folder), data=CWSCREEN_PATTERN_FULL)
anova(fp1c,fp1d)#remove Generation
rand(fp1d)

#model without random slope
fp2<-lmer(pattern_energy_diff~poly(Generation,2)*Bgd+(1|Folder), data=CWSCREEN_PATTERN_FULL)
fp2b<-lmer(pattern_energy_diff~poly(Generation,2)+Bgd+(1|Folder), data=CWSCREEN_PATTERN_FULL)
anova(fp2,fp2b)#can't remove  interaction

#is polynomial useful?
fp2_2<-lmer(pattern_energy_diff~Generation*Bgd+(1|Folder), data=CWSCREEN_PATTERN_FULL)
anova(fp2, fp2_2)#yes
#summary(fp2)

#compare models with and w/o random slope
AIC(fp1d,fp2)#model with random slope is best

#narrow colour space
np1<-lmer(pattern_energy_diff~poly(Generation,2)*Bgd+(Generation|Folder), data=CWSCREEN_PATTERN_NARROW)
#test model assumptions
res<-simulateResiduals(np1, plot=T)
hist(resid(np1))
plot(np1)#ok - no transforms improve diagnostics 

#model simplification
np1b<-lmer(pattern_energy_diff~poly(Generation,2)+Bgd+(Generation|Folder), data=CWSCREEN_PATTERN_NARROW)
anova(np1,np1b)#remove interaction
np1c<-lmer(pattern_energy_diff~poly(Generation,2)+(Generation|Folder), data=CWSCREEN_PATTERN_NARROW)
anova(np1c,np1b)#remove Bgd
np1d<-lmer(pattern_energy_diff~(Generation|Folder), data=CWSCREEN_PATTERN_NARROW)
anova(np1c,np1d)#can't remove Generation

#is polynomial needed?
np1c_2<-lmer(pattern_energy_diff~Generation+(Generation|Folder), data=CWSCREEN_PATTERN_NARROW)
anova(np1c,np1c_2)#no
#further simplification?
np1c_2b<-lmer(pattern_energy_diff~(Generation|Folder), data=CWSCREEN_PATTERN_NARROW)
anova(np1c_2b,np1c_2)#Generation still significant

#model without random slope
np2<-lmer(pattern_energy_diff~poly(Generation,2)*Bgd+(1|Folder), data=CWSCREEN_PATTERN_NARROW)
np2b<-lmer(pattern_energy_diff~poly(Generation,2)+Bgd+(1|Folder), data=CWSCREEN_PATTERN_NARROW)
anova(np2,np2b)#remove  interaction
np2c<-lmer(pattern_energy_diff~poly(Generation,2)+(1|Folder), data=CWSCREEN_PATTERN_NARROW)
anova(np2c,np2b)#remove Bgd
np2d<-lmer(pattern_energy_diff~(1|Folder), data=CWSCREEN_PATTERN_NARROW)
anova(np2c,np2d)#need Generation

#is polynomial useful?
np2c_2<-lmer(pattern_energy_diff~Generation+(1|Folder), data=CWSCREEN_PATTERN_NARROW)
anova(np2c_2, np2c)#no
anova(np2c_2,np2d)#Generation still significant
#summary(np2c_2)

#compare models with and w/o random slope
AIC(np1c_2,np2c_2)#model with random slope is best
```

### Capture time
How does capture time change across generations, in populations on different backgrounds & using different colour spaces?

```{r analysis_capture_time_1, message=FALSE, warning=FALSE,  fig.show='hide'}

#remove generation 16 from capture time data, as this was not played by participants
#and restrict to first-clicked target per screen (ie. rank 1)
CWSCREEN15<-CWSCREEN%>%
  filter(Generation!=16)%>%
  filter(Rank==1)
#separate full and narrow colour spaces for subsequent analyses
CWSCREEN15_FULL<-CWSCREEN15%>%
  filter(ColSpace=="Full")
CWSCREEN15_NARROW<-CWSCREEN15%>%
  filter(ColSpace=="Narrow")

#linear mixed effects models for each colour space in turn
#include Generation (2nd order polynomial) and Bgd, interacting + random effect of Folder(Population) with random slope for generation
fct1<-lmer(Capture_Time~poly(Generation,2)*Bgd+(Generation|Folder), data=CWSCREEN15_FULL)
#summary(fct1)

#test model assumptions
plot(fct1)
hist(resid(fct1))
#very poor, try log transform
fct1<-lmer(log(Capture_Time)~poly(Generation,2)*Bgd+(Generation|Folder), data=CWSCREEN15_FULL) 
hist(resid(fct1))
plot(fct1)#better

#simplify model
fct1b<-lmer(log(Capture_Time)~poly(Generation,2)+Bgd+(Generation|Folder), data=CWSCREEN15_FULL) 
anova(fct1,fct1b)#significant effect of background interaction

#test significance of random effects
rand(fct1)

#is the polynomial useful?
fct1_2<-lmer(log(Capture_Time)~Generation*Bgd+(Generation|Folder), data=CWSCREEN15_FULL) 
anova(fct1, fct1_2)#significant effect of polynomial, so keep it in final model

#model without random slope
fct2<-lmer(log(Capture_Time)~poly(Generation,2)*Bgd+(1|Folder), data=CWSCREEN15_FULL)
fct2b<-lmer(log(Capture_Time)~poly(Generation,2)+Bgd+(1|Folder), data=CWSCREEN15_FULL)
anova(fct2, fct2b)#can't remove Bgd:Generation interaction

#is the polynomial useful?
fct2_2<-lmer(log(Capture_Time)~Generation*Bgd+(1|Folder), data=CWSCREEN15_FULL)
anova(fct2, fct2_2)#yes, significant effect of polynomial

#compare final models with and w/o random slope:
AIC(fct1, fct2)#lower AIC with random slope

#now narrow colour space

nct1<-lmer(Capture_Time~poly(Generation,2)*Bgd+(Generation|Folder), data=CWSCREEN15_NARROW)
#summary(nct1)
#model does not converge so use model without random slope

#model without random slope
nct2<-lmer(Capture_Time~poly(Generation,2)*Bgd+(Generation|Folder), data=CWSCREEN15_NARROW) 
#test model assumptions
plot(nct2)
hist(resid(nct2))
#very poor, try log transform
nct2<-lmer(log(Capture_Time)~poly(Generation,2)*Bgd+(Generation|Folder), data=CWSCREEN15_NARROW) 
hist(resid(nct2))
plot(nct2)#better
nct2<-lmer(log(Capture_Time)~poly(Generation,2)*Bgd+(1|Folder), data=CWSCREEN15_NARROW)
nct2b<-lmer(log(Capture_Time)~poly(Generation,2)+Bgd+(1|Folder), data=CWSCREEN15_NARROW)
anova(nct2, nct2b)#can remove Bgd:Generation interaction
nct2c<-lmer(log(Capture_Time)~poly(Generation,2)+(1|Folder), data=CWSCREEN15_NARROW)
anova(nct2c, nct2b)#can remove Bgd
nct2d<-lmer(log(Capture_Time)~1+(1|Folder), data=CWSCREEN15_NARROW)
anova(nct2c, nct2d)#can't remove Generation

#is the polynomial useful?
nct2c_2<-lmer(log(Capture_Time)~Generation+(1|Folder), data=CWSCREEN15_NARROW)
anova(nct2c, nct2c_2)#no significant effect of polynomial
nct2d_2<-lmer(log(Capture_Time)~1+(1|Folder), data=CWSCREEN15_NARROW)
anova(nct2c_2, nct2d_2)#can't remove Generation

```


### Plot results
```{r figure5_prep, message=FALSE, results = 'hide', echo=FALSE, warning=FALSE}
#colours for full and narrow colour spaces
colvec<-c("#BD0026","#245D99")
colvec_full<-c( "#DD333C" ,"#FD5253","#FF6F6C","#FF8C85","#FFA99F" ,"#FFC6BA")
colvec_narrow<-c( "#4575B4" ,"#628ED0","#7EA9EC","#9BC4FF","#B7E0FF" ,"#D4FCFF")

#Luminance difference

#re-order factor levels
levels(CWSCREEN_FULL$Bgd)<-levels(CWSCREEN_NARROW$Bgd)<-c("Smooth", "Furrowed")
CWSCREEN_FULL$Folder<-factor(CWSCREEN_FULL$Folder, levels=c("Full_Sm_1", "Full_Sm_2", "Full_Sm_3","Full_Fr_1", "Full_Fr_2" ,"Full_Fr_3","Nar_Sm_1",  "Nar_Sm_2" , "Nar_Sm_3" ,"Nar_Fr_1" , "Nar_Fr_2",  "Nar_Fr_3"))
CWSCREEN_NARROW$Folder<-factor(CWSCREEN_NARROW$Folder, levels=c("Full_Sm_1", "Full_Sm_2", "Full_Sm_3","Full_Fr_1", "Full_Fr_2" ,"Full_Fr_3","Nar_Sm_1",  "Nar_Sm_2" , "Nar_Sm_3" ,"Nar_Fr_1" , "Nar_Fr_2",  "Nar_Fr_3"))

CWSCREEN_NARROW$sqrtLdiffTB<-sqrt(CWSCREEN_NARROW$LdiffTB)
CWSCREEN_NARROW$sqrtLdiffTB<-sqrt(CWSCREEN_NARROW$LdiffTB)

#full colour space, best model:
fl1d<-lmer(LdiffTB~(Generation|Folder), data=CWSCREEN_FULL) 
fltpred<-augment(fl1d)
for (i in 1: length(fltpred$Folder)){
  fltpred$Bgd[i]<-str_split( fltpred$Folder[i], "_")[[1]][2]
}

flt.sum <- fltpred %>%
  group_by(Generation,Bgd,Folder)%>%
  select(LdiffTB, .fitted, .fixed) %>% 
  summarise_each(funs(mean = mean, 
                      sd = sd))
flt.sum$lCTse<-flt.sum$LdiffTB_sd/sqrt(24)
flt.sum$Bgd<-as.factor(flt.sum$Bgd)
flt.sum$Bgd<-factor(flt.sum$Bgd,levels=c("Sm","Fr"))

indpred_flt<-ggplot(flt.sum, aes(x=Generation, y=LdiffTB_mean, group=Folder, col=Folder))+  
  geom_pointrange(aes(ymin=LdiffTB_mean-lCTse, ymax=LdiffTB_mean+lCTse, shape=Bgd), alpha=1,position = position_dodge(width = 0.3), fill="white") +
  geom_line(aes(x=Generation, y=.fitted_mean),size=0.5, linetype=3)+
  scale_colour_manual(values=colvec_full)+
  scale_shape_manual(values=c(19,21))+
  scale_linetype_manual(values=c(1,2))+
  my_theme+
  guides(col=FALSE)+guides(linetype=FALSE)+guides(shape=FALSE)+
  xlab("Generation")+ylab("L difference")+
  theme(axis.title = element_text(size = 14))  +
  theme(axis.text = element_text(size = 12)) +
  ggtitle("(i) Luminance difference") +
  theme(plot.title = element_text(face = "italic"))

#narrow colour space
nl2c_2<-lmer(sqrtLdiffTB~Generation+(1|Folder), data=CWSCREEN_NARROW)
nltpred<-augment(nl2c_2)#make predictions
for (i in 1: length(nltpred$Folder)){
  nltpred$Bgd[i]<-str_split( nltpred$Folder[i], "_")[[1]][2]
}

nlt.sum <- nltpred %>%
  group_by(Generation,Bgd,Folder)%>%
  select(sqrtLdiffTB, .fitted, .fixed) %>% 
  summarise_each(funs(mean = mean, 
                      sd = sd))
nlt.sum$lCTse<-nlt.sum$sqrtLdiffTB_sd/sqrt(24)
nlt.sum$Bgd<-as.factor(nlt.sum$Bgd)
nlt.sum$Bgd<-factor(nlt.sum$Bgd,levels=c("Sm","Fr"))

indpred_nlt<-ggplot(nlt.sum, aes(x=Generation, y=sqrtLdiffTB_mean, group=Folder, col=Folder))+  
  geom_pointrange(aes(ymin=sqrtLdiffTB_mean-lCTse, ymax=sqrtLdiffTB_mean+lCTse, shape=Bgd), alpha=1,position = position_dodge(width = 0.3), fill="white") +
  geom_line(aes(x=Generation, y=.fitted_mean),size=0.5, linetype=3)+
  geom_line(aes(x=Generation, y=.fixed_mean), size=1.5, linetype=4, col=colvec[2])+
  scale_colour_manual(values=colvec_narrow)+
  scale_shape_manual(values=c(19,21))+
  scale_linetype_manual(values=c(1,2))+
  my_theme+
  guides(col=FALSE)+guides(linetype=FALSE)+guides(shape=FALSE)+
  xlab("Generation")+ylab("sqrt(L difference)")+
  theme(axis.title = element_text(size = 14))  +
  theme(axis.text = element_text(size = 12)) +
  ggtitle("(i) Luminance difference") +
  theme(plot.title = element_text(face = "italic"))

#Colour difference

#full colour space
fc1c<-lmer(eucdisTB~poly(Generation,2)+(Generation|Folder), data=CWSCREEN_FULL) 
fcpred<-augment(fc1c)#make predictions
for (i in 1: length(fcpred$Folder)){
  fcpred$Bgd[i]<-str_split( fcpred$Folder[i], "_")[[1]][2]
}

fc.sum <- fcpred %>%
  group_by(Generation,Bgd,Folder)%>%
  select(eucdisTB, .fitted, .fixed) %>% 
  summarise_each(funs(mean = mean, 
                      sd = sd))
fc.sum$lCTse<-fc.sum$eucdisTB_sd/sqrt(24)
fc.sum$Bgd<-as.factor(fc.sum$Bgd)
fc.sum$Bgd<-factor(fc.sum$Bgd,levels=c("Sm","Fr"))

indpred_fc<-ggplot(fc.sum, aes(x=Generation, y=eucdisTB_mean, group=Folder, col=Folder))+  
  geom_pointrange(aes(ymin=eucdisTB_mean-lCTse, ymax=eucdisTB_mean+lCTse, shape=Bgd), alpha=1,position = position_dodge(width = 0.3), fill="white") +
  geom_line(aes(x=Generation, y=.fitted_mean),size=0.5,linetype=3)+
  geom_line(aes(x=Generation, y=.fixed_mean), size=1.5, linetype=4,col=colvec[1])+
  scale_colour_manual(values=colvec_full)+
  scale_shape_manual(values=c(19,21))+
  scale_linetype_manual(values=c(1,2))+
  my_theme+
  guides(col=FALSE)+guides(linetype=FALSE)+guides(shape=FALSE)+
  xlab("Generation")+ylab("Euclidean distance in CIE LAB")+
  theme(axis.title = element_text(size = 14))  +
  theme(axis.text = element_text(size = 12)) +
  ggtitle("(ii) Colour difference") +
  theme(plot.title = element_text(face = "italic"))

#narrow colspace
CWSCREEN_NARROW$sqrteucdisTB<-sqrt(CWSCREEN_NARROW$eucdisTB)
nc2c<-lmer(sqrteucdisTB~poly(Generation,2)+(1|Folder), data=CWSCREEN_NARROW)
ncpred<-augment(nc2c)#make predictions
for (i in 1: length(ncpred$Folder)){
  ncpred$Bgd[i]<-str_split( ncpred$Folder[i], "_")[[1]][2]
}
ncpred$Generation<-CWSCREEN_NARROW$Generation

nc.sum <- ncpred %>%
  group_by(Generation,Bgd,Folder)%>%
  select(sqrteucdisTB, .fitted, .fixed) %>% 
  summarise_each(funs(mean = mean, 
                      sd = sd))
nc.sum$lCTse<-nc.sum$sqrteucdisTB_sd/sqrt(24)
nc.sum$Bgd<-as.factor(nc.sum$Bgd)
nc.sum$Bgd<-factor(nc.sum$Bgd,levels=c("Sm","Fr"))

indpred_nc<-ggplot(nc.sum, aes(x=Generation, y=sqrteucdisTB_mean, group=Folder, col=Folder))+  
  geom_pointrange(aes(ymin=sqrteucdisTB_mean-lCTse, ymax=sqrteucdisTB_mean+lCTse, shape=Bgd), alpha=1,position = position_dodge(width = 0.3), fill="white") +
  geom_line(aes(x=Generation, y=.fitted_mean),size=0.5,linetype=3)+
  geom_line(aes(x=Generation, y=.fixed_mean), size=1.5, linetype=4, col=colvec[2])+
  scale_colour_manual(values=colvec_narrow)+
  scale_shape_manual(values=c(19,21))+
  scale_linetype_manual(values=c(1,2))+
  my_theme+
  guides(col=FALSE)+guides(linetype=FALSE)+guides(shape=FALSE)+
  xlab("Generation")+ylab("sqrt(Euclidean distance in CIE LAB)")+
  theme(axis.title = element_text(size = 14))  +
  theme(axis.text = element_text(size = 12)) +
  ggtitle("(ii) Colour difference") +
  theme(plot.title = element_text(face = "italic"))

#Colour difference (WA)

#full colour space
CWSCREEN_COLDIST_FULL$Folder<-factor(CWSCREEN_COLDIST_FULL$Folder, levels=c("Full_Sm_1", "Full_Sm_2", "Full_Sm_3","Full_Fr_1", "Full_Fr_2" ,"Full_Fr_3","Nar_Sm_1",  "Nar_Sm_2" , "Nar_Sm_3" ,"Nar_Fr_1" , "Nar_Fr_2",  "Nar_Fr_3"))
fcc1<-lmer(NewColMap_DeltaS~poly(Generation,2)*Bgd+(Generation|Folder), data=CWSCREEN_COLDIST_FULL)
fccpred<-augment(fcc1)#
fcc.sum <- fccpred %>%
  group_by(Generation,Bgd,Folder)%>%
  select(NewColMap_DeltaS, .fitted, .fixed) %>% 
  summarise_each(funs(mean = mean, 
                      sd = sd))
fcc.sum$lCTse<-fcc.sum$NewColMap_DeltaS_sd/sqrt(24)
fcc.sum$Bgd<-as.factor(fcc.sum$Bgd)
fcc.sum$Bgd<-factor(fcc.sum$Bgd,levels=c("Smooth","Furrowed"))

indpred_fcc<-ggplot(fcc.sum, aes(x=Generation, y=NewColMap_DeltaS_mean, group=Folder, col=Folder))+  
  geom_pointrange(aes(ymin=NewColMap_DeltaS_mean-lCTse, ymax=NewColMap_DeltaS_mean+lCTse, shape=Bgd), alpha=1,position = position_dodge(width = 0.3), fill="white") +
  geom_line(aes(x=Generation, y=.fitted_mean),size=0.5,linetype=3)+
  geom_line(aes(x=Generation, y=.fixed_mean, linetype=Bgd), size=1.5, col=colvec[1])+
  scale_colour_manual(values=colvec_full)+
  scale_shape_manual(values=c(19,21))+
  scale_linetype_manual(values=c(1,2))+
  my_theme+
  guides(col=FALSE)+guides(linetype=FALSE)+guides(shape=FALSE)+
  xlab("Generation")+ylab("Colour difference (Delta-S)")+
  theme(axis.title = element_text(size = 14))  +
  theme(axis.text = element_text(size = 12)) +
  ggtitle("(iii) Weighted average colour difference") +
  theme(plot.title = element_text(face = "italic"))

#narrow colspace - requires diffferent technique to mkae preictions from this type of model
CWSCREEN_COLDIST_NARROW$Folder<-factor(CWSCREEN_COLDIST_NARROW$Folder, levels=c("Full_Sm_1", "Full_Sm_2", "Full_Sm_3","Full_Fr_1", "Full_Fr_2" ,"Full_Fr_3","Nar_Sm_1",  "Nar_Sm_2" , "Nar_Sm_3" ,"Nar_Fr_1" , "Nar_Fr_2",  "Nar_Fr_3"))

ncc1<-glmmTMB(ColDistRound ~ poly(Generation,2)*Bgd+(Generation|Folder), data=CWSCREEN_COLDIST_NARROW, family=nbinom1 )

nccpred1 <- as.data.frame(ggpredict(ncc1, terms = c("Generation[all]","Bgd[Smooth]","Folder[Nar_Sm_1]"), type="random"))
nccpred2 <- as.data.frame(ggpredict(ncc1, terms = c("Generation[all]","Bgd[Smooth]","Folder[Nar_Sm_2]"), type="random"))
nccpred3 <- as.data.frame(ggpredict(ncc1, terms = c("Generation[all]","Bgd[Smooth]","Folder[Nar_Sm_3]"), type="random"))
nccpred4 <- as.data.frame(ggpredict(ncc1, terms = c("Generation[all]","Bgd[Furrowed]","Folder[Nar_Fr_1]"), type="random"))
nccpred5 <- as.data.frame(ggpredict(ncc1, terms = c("Generation[all]","Bgd[Furrowed]","Folder[Nar_Fr_2]"), type="random"))
nccpred6 <- as.data.frame(ggpredict(ncc1, terms = c("Generation[all]","Bgd[Furrowed]","Folder[Nar_Fr_3]"), type="random"))

#and predictions for fixed effects only
nccpred1_fonly <- as.data.frame(ggpredict(ncc1, terms = c("Generation[all]","Bgd[Smooth]"), type="fixed"))
nccpred2_fonly <- as.data.frame(ggpredict(ncc1, terms = c("Generation[all]","Bgd[Furrowed]"), type="fixed"))


coldist_sum<-CWSCREEN_COLDIST_NARROW%>%
  group_by(Bgd, Folder,Generation)%>%
  summarise(mean=mean(ColDistRound),sd=sd(ColDistRound))
coldist_sum<-as.data.frame(coldist_sum)
coldist_sum$Folder<-factor(coldist_sum$Folder, levels=c("Full_Sm_1", "Full_Sm_2", "Full_Sm_3","Full_Fr_1", "Full_Fr_2" ,"Full_Fr_3","Nar_Sm_1",  "Nar_Sm_2" , "Nar_Sm_3" ,"Nar_Fr_1" , "Nar_Fr_2",  "Nar_Fr_3"))

for(i in 1:length(coldist_sum$Generation)){
  targetgen<-coldist_sum$Generation[i]
  
  if(coldist_sum$Folder[i]=="Nar_Sm_1"){
    predict_val<-nccpred1

    for(j in 1:length(predict_val$predicted)){
      if(predict_val$x[j]==targetgen){
        coldist_sum$predicted[i]<-predict_val$predicted[j]
      }
      
    }
  }
  if(coldist_sum$Folder[i]=="Nar_Sm_2"){
    predict_val<-nccpred2
    
    for(j in 1:length(predict_val$predicted)){
      if(predict_val$x[j]==targetgen){
        coldist_sum$predicted[i]<-predict_val$predicted[j]
      }
      
    }
  }
  if(coldist_sum$Folder[i]=="Nar_Sm_3"){
    predict_val<-nccpred3
    
    for(j in 1:length(predict_val$predicted)){
      if(predict_val$x[j]==targetgen){
        coldist_sum$predicted[i]<-predict_val$predicted[j]
      }
      
    }
  }
  if(coldist_sum$Folder[i]=="Nar_Fr_1"){
    predict_val<-nccpred4
    
    for(j in 1:length(predict_val$predicted)){
      if(predict_val$x[j]==targetgen){
        coldist_sum$predicted[i]<-predict_val$predicted[j]
      }
      
    }
  }
  if(coldist_sum$Folder[i]=="Nar_Fr_2"){
    predict_val<-nccpred5
    
    for(j in 1:length(predict_val$predicted)){
      if(predict_val$x[j]==targetgen){
        coldist_sum$predicted[i]<-predict_val$predicted[j]
      }
      
    }
  }
  if(coldist_sum$Folder[i]=="Nar_Fr_3"){
    predict_val<-nccpred6
    
    for(j in 1:length(predict_val$predicted)){
      if(predict_val$x[j]==targetgen){
        coldist_sum$predicted[i]<-predict_val$predicted[j]
      }
      
    }
  }
  
  if(coldist_sum$Bgd[i]=="Smooth"){
    predict_fixed<-nccpred1_fonly
    for(k in 1: length(predict_fixed$predicted)){
      if(predict_fixed$x[k]==targetgen){
        coldist_sum$fixed[i]<-predict_fixed$predicted[k]
      }
    }
  }
  else {
    predict_fixed<-nccpred2_fonly
    for(k in 1: length(predict_fixed$predicted)){
      if(predict_fixed$x[k]==targetgen){
        coldist_sum$fixed[i]<-predict_fixed$predicted[k]
      }
    }
  }
  
}
coldist_sum$se<-coldist_sum$sd/sqrt(24)
coldist_sum$Bgd<-factor(coldist_sum$Bgd, levels=c("Smooth", "Furrowed"))
indpred_ncc<-ggplot(coldist_sum, aes(x=Generation, y=mean, group=Folder, col=Folder))+  
  geom_pointrange(aes(ymin=mean-se, ymax=mean+se, shape=Bgd), alpha=1,position = position_dodge(width = 0.3), fill="white") +
  geom_line(aes(x=Generation, y=predicted),size=0.5,linetype=3)+
  geom_line(aes(x=Generation, y=fixed, linetype=Bgd), size=1.5,  col=colvec[2])+
  scale_colour_manual(values=colvec_narrow)+
  scale_shape_manual(values=c(19,21))+
  scale_linetype_manual(values=c(1,2))+
  my_theme+
  guides(col=FALSE)+guides(linetype=FALSE)+guides(shape=FALSE)+
  xlab("Generation")+ylab("Colour difference (Delta-S)")+
  theme(axis.title = element_text(size = 14))  +
  theme(axis.text = element_text(size = 12)) +
  ggtitle("(iii) Weighted average colour difference") +
  theme(plot.title = element_text(face = "italic"))

#Pattern difference
#full colour space
fp1d<-lmer(pattern_energy_diff~(Generation|Folder), data=CWSCREEN_PATTERN_FULL)
fppred<-augment(fp1d)#make predictions
for (i in 1: length(fppred$Folder)){
  fppred$Bgd[i]<-str_split( fppred$Folder[i], "_")[[1]][2]
}

fp.sum <- fppred %>%
  group_by(Generation,Bgd,Folder)%>%
  select(pattern_energy_diff, .fitted, .fixed) %>% 
  summarise_each(funs(mean = mean, 
                      sd = sd))
fp.sum$lCTse<-fp.sum$pattern_energy_diff_sd/sqrt(24)
fp.sum$Bgd<-as.factor(fp.sum$Bgd)
fp.sum$Bgd<-factor(fp.sum$Bgd,levels=c("Sm","Fr"))

fp.sum$Folder<-factor(fp.sum$Folder, levels=c("Full_Sm_1", "Full_Sm_2", "Full_Sm_3","Full_Fr_1", "Full_Fr_2" ,"Full_Fr_3","Nar_Sm_1",  "Nar_Sm_2" , "Nar_Sm_3" ,"Nar_Fr_1" , "Nar_Fr_2",  "Nar_Fr_3"))

indpred_fp<-ggplot(fp.sum, aes(x=Generation, y=pattern_energy_diff_mean, group=Folder, col=Folder))+  
  geom_pointrange(aes(ymin=pattern_energy_diff_mean-lCTse, ymax=pattern_energy_diff_mean+lCTse, shape=Bgd), alpha=1,position = position_dodge(width = 0.3), fill="white") +
  geom_line(aes(x=Generation, y=.fitted_mean),size=0.5,linetype=3)+
  scale_colour_manual(values=colvec_full)+
  scale_shape_manual(values=c(19,21))+
  scale_linetype_manual(values=c(1,2))+
  my_theme+
  guides(col=FALSE)+guides(linetype=FALSE)+guides(shape=FALSE)+
  xlab("Generation")+ylab("Energy difference")+
  theme(axis.title = element_text(size = 14))  +
  theme(axis.text = element_text(size = 12)) +
  ggtitle("(iv) Pattern difference") +
  theme(plot.title = element_text(face = "italic"))

#narrow colour space
np1c_2<-lmer(pattern_energy_diff~Generation+(Generation|Folder), data=CWSCREEN_PATTERN_NARROW)
nppred<-augment(np1c_2)#make predictions
for (i in 1: length(nppred$Folder)){
  nppred$Bgd[i]<-str_split( nppred$Folder[i], "_")[[1]][2]
}

np.sum <- nppred %>%
  group_by(Generation,Bgd,Folder)%>%
  select(pattern_energy_diff, .fitted, .fixed) %>% 
  summarise_each(funs(mean = mean, 
                      sd = sd))
np.sum$lCTse<-np.sum$pattern_energy_diff_sd/sqrt(24)
np.sum$Bgd<-as.factor(np.sum$Bgd)
np.sum$Bgd<-factor(np.sum$Bgd,levels=c("Sm","Fr"))
np.sum$Folder<-factor(np.sum$Folder, levels=c("Full_Sm_1", "Full_Sm_2", "Full_Sm_3","Full_Fr_1", "Full_Fr_2" ,"Full_Fr_3","Nar_Sm_1",  "Nar_Sm_2" , "Nar_Sm_3" ,"Nar_Fr_1" , "Nar_Fr_2",  "Nar_Fr_3"))

indpred_np<-ggplot(np.sum, aes(x=Generation, y=pattern_energy_diff_mean, group=Folder, col=Folder))+  
  geom_pointrange(aes(ymin=pattern_energy_diff_mean-lCTse, ymax=pattern_energy_diff_mean+lCTse, shape=Bgd), alpha=1,position = position_dodge(width = 0.3), fill="white") +
  geom_line(aes(x=Generation, y=.fitted_mean),size=0.5,linetype=3)+
  geom_line(aes(x=Generation, y=.fixed_mean), size=1.5, linetype=4, col=colvec[2])+
  scale_colour_manual(values=colvec_narrow)+
  scale_shape_manual(values=c(19,21))+
  scale_linetype_manual(values=c(1,2))+
  my_theme+
  guides(col=FALSE)+guides(linetype=FALSE)+guides(shape=FALSE)+
  xlab("Generation")+ylab("Energy difference")+
  theme(axis.title = element_text(size = 14))  +
  theme(axis.text = element_text(size = 12)) +
  ggtitle("(iv) Pattern difference") +
  theme(plot.title = element_text(face = "italic"))

#Edge disruption
#gabrat
fgr1d<-lmer(Target_L_GabRat~(Generation|Folder), data=CWSCREEN_FULL)
fgpred<-augment(fgr1d)#make predictions
for (i in 1: length(fgpred$Folder)){
  fgpred$Bgd[i]<-str_split( fgpred$Folder[i], "_")[[1]][2]
}

fg.sum <- fgpred %>%
  group_by(Generation,Bgd,Folder)%>%
  select(Target_L_GabRat, .fitted, .fixed) %>% 
  summarise_each(funs(mean = mean, 
                      sd = sd))
fg.sum$lCTse<-fg.sum$Target_L_GabRat_sd/sqrt(24)
fg.sum$Bgd<-as.factor(fg.sum$Bgd)
fg.sum$Bgd<-factor(fg.sum$Bgd,levels=c("Sm","Fr"))

indpred_fg<-ggplot(fg.sum, aes(x=Generation, y=Target_L_GabRat_mean, group=Folder, col=Folder))+  
  geom_pointrange(aes(ymin=Target_L_GabRat_mean-lCTse, ymax=Target_L_GabRat_mean+lCTse, shape=Bgd), alpha=1,position = position_dodge(width = 0.3), fill="white") +
  geom_line(aes(x=Generation, y=.fitted_mean),size=0.5,linetype=3)+
  scale_colour_manual(values=colvec_full)+
  scale_shape_manual(values=c(19,21))+
  scale_linetype_manual(values=c(1,2))+
  my_theme+
  guides(col=FALSE)+guides(linetype=FALSE)+guides(shape=FALSE)+
  xlab("Generation")+ylab("GabRat in L")+
  theme(axis.title = element_text(size = 14))  +
  theme(axis.text = element_text(size = 12)) +
  ggtitle("(v) Edge disruption") +
  theme(plot.title = element_text(face = "italic"))

#narrow colour space
ngr2c_2<-lmer(Target_L_GabRat~Generation+(1|Folder), data=CWSCREEN_NARROW)
ngpred<-augment(ngr2c_2)#make predictions
for (i in 1: length(ngpred$Folder)){
  ngpred$Bgd[i]<-str_split( ngpred$Folder[i], "_")[[1]][2]
}

ng.sum <- ngpred %>%
  group_by(Generation,Bgd,Folder)%>%
  select(Target_L_GabRat, .fitted, .fixed) %>% 
  summarise_each(funs(mean = mean, 
                      sd = sd))
ng.sum$lCTse<-ng.sum$Target_L_GabRat_sd/sqrt(24)
ng.sum$Bgd<-as.factor(ng.sum$Bgd)
ng.sum$Bgd<-factor(ng.sum$Bgd,levels=c("Sm","Fr"))

indpred_ng<-ggplot(ng.sum, aes(x=Generation, y=Target_L_GabRat_mean, group=Folder, col=Folder))+  
  geom_pointrange(aes(ymin=Target_L_GabRat_mean-lCTse, ymax=Target_L_GabRat_mean+lCTse, shape=Bgd), alpha=1,position = position_dodge(width = 0.3), fill="white") +
  geom_line(aes(x=Generation, y=.fitted_mean),size=0.5,linetype=3)+
  geom_line(aes(x=Generation, y=.fixed_mean), size=1.5, linetype=4, col=colvec[2])+
  scale_colour_manual(values=colvec_narrow)+
  scale_shape_manual(values=c(19,21))+
  scale_linetype_manual(values=c(1,2))+
  my_theme+
  guides(col=FALSE)+guides(linetype=FALSE)+guides(shape=FALSE)+
  xlab("Generation")+ylab("GabRat in L")+
  theme(axis.title = element_text(size = 14))  +
  theme(axis.text = element_text(size = 12)) +
  ggtitle("(v) Edge disruption") +
  theme(plot.title = element_text(face = "italic"))


#Capture time
CWSCREEN15_FULL$logCaptureTime<-log(CWSCREEN15_FULL$Capture_Time)
CWSCREEN15_NARROW$logCaptureTime<-log(CWSCREEN15_NARROW$Capture_Time)

#re-order factor levels
levels(CWSCREEN15_FULL$Bgd)<-levels(CWSCREEN15_NARROW$Bgd)<-c("Smooth", "Furrowed")
levels(CWSCREEN15_FULL$Folder)<-c("Full_Sm_1", "Full_Sm_2", "Full_Sm_3","Full_Fr_1", "Full_Fr_2" ,"Full_Fr_3","Nar_Sm_1",  "Nar_Sm_2" , "Nar_Sm_3" ,"Nar_Fr_1" , "Nar_Fr_2",  "Nar_Fr_3")
levels(CWSCREEN15_NARROW$Folder)<-c("Full_Sm_1", "Full_Sm_2", "Full_Sm_3","Full_Fr_1", "Full_Fr_2" ,"Full_Fr_3","Nar_Sm_1",  "Nar_Sm_2" , "Nar_Sm_3" ,"Nar_Fr_1" , "Nar_Fr_2",  "Nar_Fr_3")

#best model for full colour space
fct1<-lmer(logCaptureTime~poly(Generation,2)*Bgd+(Generation|Folder), data=CWSCREEN15_FULL) 
fctpred<-augment(fct1)#make predictions
fct.sum <- fctpred %>%
  group_by(Generation,Bgd,Folder)%>%
  select(logCaptureTime, .fitted, .fixed) %>% 
  summarise_each(funs(mean = mean, 
                      sd = sd))
fct.sum$lCTse<-fct.sum$logCaptureTime_sd/sqrt(24)

indpred_fct<-ggplot(fct.sum, aes(x=Generation, y=logCaptureTime_mean, group=Folder, col=Folder))+  
  geom_pointrange(aes(ymin=logCaptureTime_mean-lCTse, ymax=logCaptureTime_mean+lCTse, shape=Bgd), alpha=1,position = position_dodge(width = 0.3), fill="white") +
  geom_line(aes(x=Generation, y=.fitted_mean),size=0.5,linetype=3)+
  geom_line(aes(x=Generation, y=.fixed_mean, linetype=Bgd), size=1.5, col=colvec[1])+
  scale_colour_manual(values=colvec_full)+
  scale_shape_manual(values=c(19,21))+
  scale_linetype_manual(values=c(1,2))+
  my_theme+
  guides(col=FALSE)+guides(linetype=FALSE)+guides(shape=FALSE)+
  xlab("Generation")+ylab("log(Capture time (ms))")+
  theme(axis.title = element_text(size = 14))  +
  theme(axis.text = element_text(size = 12)) +
  ggtitle("(vi) Capture time") +
  theme(plot.title = element_text(face = "italic"))

#now for narrow colour space
nct2c_2<-lmer(logCaptureTime~Generation+(1|Folder), data=CWSCREEN15_NARROW)
nctpred<-augment(nct2c_2)#make predictions
nct.sum <- nctpred %>%
  group_by(Generation,Folder)%>%
  select(logCaptureTime, .fitted, .fixed) %>% 
  summarise_each(funs(mean = mean, 
                      sd = sd))
nct.sum$lCTse<-nct.sum$logCaptureTime_sd/sqrt(24)

indpred_nct<-ggplot(nct.sum, aes(x=Generation, y=logCaptureTime_mean, group=Folder, col=Folder))+  
  geom_pointrange(aes(ymin=logCaptureTime_mean-lCTse, ymax=logCaptureTime_mean+lCTse, shape=Folder), alpha=1,position = position_dodge(width = 0.3), fill="white") +
  geom_line(aes(x=Generation, y=.fitted_mean),size=0.5, linetype=3)+
  geom_line(aes(x=Generation, y=.fixed_mean), size=1.5, col=colvec[2], linetype=4)+
  scale_colour_manual(values=colvec_narrow)+
  scale_shape_manual(values=c(19,19,19,21,21,21))+
  my_theme+
  guides(col=FALSE)+guides(linetype=FALSE)+guides(shape=FALSE)+
  xlab("Generation")+ylab("log(Capture time (ms))")+
  theme(axis.title = element_text(size = 14))  +
  theme(axis.text = element_text(size = 12)) +
  ggtitle("(vi) Capture time") +
  theme(plot.title = element_text(face = "italic"))


```

```{r figure5, message=FALSE, results = 'hide', echo=FALSE, warning=FALSE, fig.dim=c(20,30), fig.cap="Figure 5: Changes in camouflage metrics in the screen-based experiments"}
(indpred_nlt+indpred_flt)/
  (indpred_nc+indpred_fc)/
  (indpred_ncc+indpred_fcc)/
  (indpred_np+indpred_fp)/
  (indpred_ng+indpred_fg)/
  (indpred_nct+indpred_fct)
```

### Experimental versus control populations

Do the trajectories of experimental (selected) populations and control (neutral drift) replicates diverge across generations?
Key variables are as above; SelectionType represents the scenario described in SI (control or experimental)

```{r prep_data_control_1, message=FALSE, warning=FALSE, echo=FALSE}
#add selection type column to screen datasets
CWSCREEN15_FULL$SelectionType<-rep("Human",length(CWSCREEN15_FULL$ID))
CWSCREEN_PATTERN_FULL$SelectionType<-rep("Human",length(CWSCREEN_PATTERN_FULL$PreyID))
CWSCREEN_COLDIST_FULL$SelectionType<-rep("Human",length(CWSCREEN_COLDIST_FULL$ID))
CWSCREEN15_NARROW$SelectionType<-rep("Human",length(CWSCREEN15_NARROW$ID))
CWSCREEN_PATTERN_NARROW$SelectionType<-rep("Human",length(CWSCREEN_PATTERN_NARROW$PreyID))
CWSCREEN_COLDIST_NARROW$SelectionType<-rep("Human",length(CWSCREEN_COLDIST_NARROW$ID))
```
```{r prep_data_control_2, message=FALSE, warning=FALSE}

#load control population data
CWSCREEN_CONTROL<-read.csv("CamoWild_ScreenControl.csv", header=TRUE)
CWSCREEN_COLDIST_CONTROL<-read.csv("CamoWild_ScreenControl_ColDist.csv", header=TRUE)

#add useful variables
#colour difference between means - background to target (euclidean distance) 
CWSCREEN_CONTROL$eucdisTB<-sqrt((CWSCREEN_CONTROL$Target_L_Mean-CWSCREEN_CONTROL$BG_L_Mean)^2+
                                  (CWSCREEN_CONTROL$Target_A_Mean-CWSCREEN_CONTROL$BG_A_Mean)^2+
                                 (CWSCREEN_CONTROL$ Target_B_Mean-CWSCREEN_CONTROL$BG_B_Mean)^2)
#luminance difference (difference in L values)
CWSCREEN_CONTROL$LdiffTB<-abs(CWSCREEN_CONTROL$Target_L_Mean-CWSCREEN_CONTROL$BG_L_Mean)

#separate into full and narrow colour spaces
CWSCREEN_CONTROL_FULL<-CWSCREEN_CONTROL%>%
  filter(ColSpace=="Full")
CWSCREEN_CONTROL_NARROW<-CWSCREEN_CONTROL%>%
  filter(ColSpace=="Narrow")
CWSCREEN_COLDIST_CONTROL_FULL<-CWSCREEN_COLDIST_CONTROL%>%
  filter(ColSpace=="Full")
CWSCREEN_COLDIST_CONTROL_NARROW<-CWSCREEN_COLDIST_CONTROL%>%
  filter(ColSpace=="Narrow")

#combine experimental and control datasets

CWSCREEN15_FULL2<-CWSCREEN15_FULL%>%
  select(Generation,ID, ParticipantID,Folder,ColSpace,Bgd,SelectionType,eucdisTB,LdiffTB,Target_L_GabRat)
CWSCREEN15_NARROW2<-CWSCREEN15_NARROW%>%
  select(Generation,ID, ParticipantID,Folder,ColSpace,Bgd,SelectionType,eucdisTB,LdiffTB,Target_L_GabRat)
CWSCREEN_CONTROL_FULL2<-CWSCREEN_CONTROL_FULL%>%
  select(Generation,ID, ParticipantID,Folder,ColSpace,Bgd,SelectionType,eucdisTB,LdiffTB,Target_L_GabRat)
CWSCREEN_CONTROL_NARROW2<-CWSCREEN_CONTROL_NARROW%>%
  select(Generation,ID, ParticipantID,Folder,ColSpace,Bgd,SelectionType,eucdisTB,LdiffTB,Target_L_GabRat)

COMBINED_FULL<-rbind(CWSCREEN15_FULL2,CWSCREEN_CONTROL_FULL2)
COMBINED_NARROW<-rbind(CWSCREEN15_NARROW2,CWSCREEN_CONTROL_NARROW2)

#colour distance dataset

#restrict experimental datasets to 15 generations only
CWSCREEN15_COLDIST_FULL<-CWSCREEN_COLDIST_FULL%>%
  filter(Generation!=16)
CWSCREEN15_COLDIST_NARROW<-CWSCREEN_COLDIST_NARROW%>%
  filter(Generation!=16)

#add rounded colour distance and restrict to useful variaibles only
CWSCREEN_COLDIST_CONTROL_FULL$ColDistRound<-round(CWSCREEN_COLDIST_CONTROL_FULL$NewColMap_DeltaS)
CWSCREEN_COLDIST_CONTROL_NARROW$ColDistRound<-round(CWSCREEN_COLDIST_CONTROL_NARROW$NewColMap_DeltaS)
CWSCREEN_COLDIST_CONTROL_FULL<-CWSCREEN_COLDIST_CONTROL_FULL%>%
  select(ColSpace, Bgd, Folder, Generation, ID, NewColMap_DeltaS, ColDistRound, SelectionType)
CWSCREEN_COLDIST_CONTROL_NARROW<-CWSCREEN_COLDIST_CONTROL_NARROW%>%
  select(ColSpace, Bgd, Folder, Generation, ID, NewColMap_DeltaS, ColDistRound, SelectionType)
#combine datasets
COMBINED_COLDIST_FULL<-rbind(CWSCREEN15_COLDIST_FULL,CWSCREEN_COLDIST_CONTROL_FULL)
COMBINED_COLDIST_NARROW<-rbind(CWSCREEN15_COLDIST_NARROW,CWSCREEN_COLDIST_CONTROL_NARROW)
```

#### Trends in luminance difference

Full colour space:

```{r control_lumdiff1, message=FALSE, warning=FALSE, fig.show='hide'}
combimodl1<-lmer(sqrt(LdiffTB)~poly(Generation,2)*SelectionType+(Generation|Folder), data=COMBINED_FULL)
plot(combimodl1)
hist(resid(combimodl1))#sqrt transform is best

combimodl1b<-lmer(sqrt(LdiffTB)~poly(Generation,2)+SelectionType+(Generation|Folder), data=COMBINED_FULL)
anova(combimodl1, combimodl1b)#remove interaction - no difference between exptl and control runs
combimodl1c<-lmer(sqrt(LdiffTB)~poly(Generation,2)+(Generation|Folder), data=COMBINED_FULL)
anova(combimodl1c, combimodl1b)#remove selection type
combimodl1d<-lmer(sqrt(LdiffTB)~(Generation|Folder), data=COMBINED_FULL)
anova(combimodl1c, combimodl1d)#significant effect of generation 
#is the polynomial useful?
combimodl1e<-lmer(sqrt(LdiffTB)~Generation+(Generation|Folder), data=COMBINED_FULL)
anova(combimodl1c, combimodl1e)#yes
#and random effects?
rand(combimodl1c)#yes
```
```{r control_lumdiff2, message=FALSE, warning=FALSE, results='hide', fig.show='hide'}
#without random slope
combimodl1i<-lmer(sqrt(LdiffTB)~poly(Generation,2)*SelectionType+(1|Folder), data=COMBINED_FULL)
plot(combimodl1)
hist(resid(combimodl1))#ok

combimodl1ib<-lmer(sqrt(LdiffTB)~poly(Generation,2)+SelectionType+(1|Folder), data=COMBINED_FULL)
anova(combimodl1i, combimodl1ib)#remove interaction - no difference between exptl and random run
combimodl1ic<-lmer(sqrt(LdiffTB)~poly(Generation,2)+(1|Folder), data=COMBINED_FULL)
anova(combimodl1ic, combimodl1ib)#remove selection type
combimodl1id<-lmer(sqrt(LdiffTB)~(1|Folder), data=COMBINED_FULL)
anova(combimodl1ic, combimodl1id)#significant effect of generation
#polynomial useful?
combimodl1ie<-lmer(sqrt(LdiffTB)~Generation+(1|Folder), data=COMBINED_FULL)
anova(combimodl1ic, combimodl1ie)#yes
```
```{r control_lumdiff3, message=FALSE, warning=FALSE}
AIC(combimodl1c,combimodl1ic)#use model with random slope
```
Narrow colour space:

```{r control_lumdiff4, message=FALSE, warning=FALSE, fig.show='hide'}
combimodl2<-lmer(LdiffTB~poly(Generation,2)*SelectionType+(Generation|Folder), data=COMBINED_NARROW)
plot(combimodl2)
hist(resid(combimodl2))
combimodl2<-lmer(sqrt(LdiffTB)~poly(Generation,2)*SelectionType+(Generation|Folder), data=COMBINED_NARROW)
plot(combimodl2)
hist(resid(combimodl2))#better

combimodl2b<-lmer(sqrt(LdiffTB)~poly(Generation,2)+SelectionType+(Generation|Folder), data=COMBINED_NARROW)
anova(combimodl2, combimodl2b)#sig interaction - difference between exptl and random runs
combimodl2c<-lmer(sqrt(LdiffTB)~Generation*SelectionType+(Generation|Folder), data=COMBINED_NARROW)
anova(combimodl2c, combimodl2)#polynomial not needed
combimodl2d<-lmer(sqrt(LdiffTB)~Generation+SelectionType+(Generation|Folder), data=COMBINED_NARROW)
anova(combimodl2c, combimodl2d)#interaction still needed
rand(combimodl2c)#random effects useful
```
```{r control_lumdiff5, message=FALSE, warning=FALSE, results='hide', fig.show='hide'}
#without random slope
combimodl2c_simple<-lmer(sqrt(LdiffTB)~poly(Generation,2)*SelectionType+(1|Folder), data=COMBINED_NARROW)
combimodl2d_simple<-lmer(sqrt(LdiffTB)~poly(Generation,2)+SelectionType+(1|Folder), data=COMBINED_NARROW)
anova(combimodl2c_simple,combimodl2d_simple)#needs interaction
combimodl2e_simple<-lmer(sqrt(LdiffTB)~Generation*SelectionType+(1|Folder), data=COMBINED_NARROW)
anova(combimodl2c_simple,combimodl2e_simple)#don't need poly
combimodl2f_simple<-lmer(sqrt(LdiffTB)~Generation+SelectionType+(1|Folder), data=COMBINED_NARROW)
anova(combimodl2e_simple,combimodl2f_simple)#still a significant interaction
```
```{r control_lumdiff6, message=FALSE, warning=FALSE}
AIC(combimodl2e_simple,combimodl2c)#random slope best
```

#### Trends in colour difference

Full colour space:
```{r control_coldiff1, message=FALSE, warning=FALSE, fig.show='hide'}
combimodc1<-lmer(sqrt(eucdisTB)~poly(Generation,2)*SelectionType+(Generation|Folder), data=COMBINED_FULL)
plot(combimodc1)
hist(resid(combimodc1))#ok

combimodc1b<-lmer(sqrt(eucdisTB)~poly(Generation,2)+SelectionType+(Generation|Folder), data=COMBINED_FULL)
anova(combimodc1, combimodc1b)#can remove interaction  
combimodc1c<-lmer(sqrt(eucdisTB)~poly(Generation,2)+(Generation|Folder), data=COMBINED_FULL)
anova(combimodc1c, combimodc1b)#remove scenario
combimodc1d<-lmer(sqrt(eucdisTB)~1+(Generation|Folder), data=COMBINED_FULL)
anova(combimodc1c, combimodc1d)#need generation
combimodc1e<-lmer(sqrt(eucdisTB)~Generation+(Generation|Folder), data=COMBINED_FULL)
anova(combimodc1e, combimodc1c)#polynomial useful 
#and random effects?
rand(combimodc1c)#yes
#summary(combimodc1c)
```
```{r control_coldiff2, message=FALSE, warning=FALSE, results='hide', fig.show='hide'}
#without random slope
combimodc1i<-lmer(sqrt(eucdisTB)~poly(Generation,2)*SelectionType+(1|Folder), data=COMBINED_FULL)
plot(combimodc1)
hist(resid(combimodc1))#ok

combimodc1bi<-lmer(sqrt(eucdisTB)~poly(Generation,2)+SelectionType+(1|Folder), data=COMBINED_FULL)
anova(combimodc1i, combimodc1bi)#can't remove interaction  
combimodc1ci<-lmer(sqrt(eucdisTB)~Generation*SelectionType+(1|Folder), data=COMBINED_FULL)
anova(combimodc1ci, combimodc1i)#polynomial useful

```
```{r control_coldiff3, message=FALSE, warning=FALSE}
AIC(combimodc1c,combimodc1i)#use model with random slope
```
Narrow colour space:

```{r control_coldiff4, message=FALSE, warning=FALSE, fig.show='hide'}
combimodc2<-lmer(eucdisTB~poly(Generation,2)*SelectionType+(Generation|Folder), data=COMBINED_NARROW)
plot(combimodc2)
hist(resid(combimodc2))
combimodc2<-lmer(sqrt(eucdisTB)~poly(Generation,2)*SelectionType+(Generation|Folder), data=COMBINED_NARROW)
plot(combimodc2)
hist(resid(combimodc2))#better

combimodc2b<-lmer(sqrt(eucdisTB)~poly(Generation,2)+SelectionType+(Generation|Folder), data=COMBINED_NARROW)
anova(combimodc2, combimodc2b)#sig interaction - difference between exptl and random runs
combimodc2c<-lmer(sqrt(eucdisTB)~Generation*SelectionType+(Generation|Folder), data=COMBINED_NARROW)
anova(combimodc2c, combimodc2)#polynomial not needed
combimodc2d<-lmer(sqrt(eucdisTB)~Generation+SelectionType+(Generation|Folder), data=COMBINED_NARROW)
anova(combimodc2c, combimodc2d)#interaction still needed
rand(combimodc2c)#random effects useful
```
```{r control_coldiff5, message=FALSE, warning=FALSE, results='hide', fig.show='hide'}
#without random slope
combimodc2i<-lmer(sqrt(eucdisTB)~poly(Generation,2)*SelectionType+(1|Folder), data=COMBINED_NARROW)
combimodc2ib<-lmer(sqrt(eucdisTB)~poly(Generation,2)+SelectionType+(1|Folder), data=COMBINED_NARROW)
anova(combimodc2i, combimodc2ib)#sig interaction - difference between exptl and random runs
combimodc2ic<-lmer(sqrt(eucdisTB)~Generation*SelectionType+(1|Folder), data=COMBINED_NARROW)
anova(combimodc2ic, combimodc2i)#polynomial not needed
combimodc2id<-lmer(sqrt(eucdisTB)~Generation+SelectionType+(1|Folder), data=COMBINED_NARROW)
anova(combimodc2ic, combimodc2id)
```
```{r control_coldiff6, message=FALSE, warning=FALSE}
AIC(combimodc2c,combimodc2ic)#random slope best
```
#### Trends in colour difference (weighted average method)
Full colour space:
```{r control_coldiff7, message=FALSE, warning=FALSE, fig.show='hide'}
#use regular model for full, negative binomial for narrow colour space
combimodc3<-lmer(NewColMap_DeltaS~poly(Generation,2)*SelectionType+(Generation|Folder), data=COMBINED_COLDIST_FULL)
plot(combimodc3)
hist(resid(combimodc3))#ok

combimodc3b<-lmer(NewColMap_DeltaS~poly(Generation,2)+SelectionType+(Generation|Folder), data=COMBINED_COLDIST_FULL)
anova(combimodc3, combimodc3b)#remove interaction - no difference between exptl and control run
combimodc3c<-lmer(NewColMap_DeltaS~poly(Generation,2)+(Generation|Folder), data=COMBINED_COLDIST_FULL)
anova(combimodc3c, combimodc3b)#remove selection type
combimodc3d<-lmer(NewColMap_DeltaS~(Generation|Folder), data=COMBINED_COLDIST_FULL)
anova(combimodc3c, combimodc3d)#significant effect of generation 
#polynomial useful?
combimodc3e<-lmer(NewColMap_DeltaS~Generation+(Generation|Folder), data=COMBINED_COLDIST_FULL)
anova(combimodc3c, combimodc3e)#no
combimodc3f<-lmer(NewColMap_DeltaS~(Generation|Folder), data=COMBINED_COLDIST_FULL)
anova(combimodc3f, combimodc3e)#generation still significant
#and random effects?
rand(combimodc3e)#yes
#summary(combimodc3e)
```
```{r control_coldiff8, message=FALSE, warning=FALSE, results='hide', fig.show='hide'}
#without random slope
combimodc3_2<-lmer(NewColMap_DeltaS~poly(Generation,2)*SelectionType+(1|Folder), data=COMBINED_COLDIST_FULL)
combimodc3b_2<-lmer(NewColMap_DeltaS~poly(Generation,2)+SelectionType+(1|Folder), data=COMBINED_COLDIST_FULL)
anova(combimodc3_2, combimodc3b_2)#cannot remove interaction - difference between exptl and control run
combimodc3c_2<-lmer(NewColMap_DeltaS~Generation*SelectionType+(1|Folder), data=COMBINED_COLDIST_FULL)
anova(combimodc3_2, combimodc3c_2)#polynomial significant
```
```{r control_coldiff9, message=FALSE, warning=FALSE}
AIC(combimodc3e,combimodc3_2)#use model with random slope
```
Narrow colour space:

```{r control_coldiff10, message=FALSE, warning=FALSE, fig.show='hide'}
combimodc4<-glmmTMB(ColDistRound ~ poly(Generation,2)*SelectionType+(Generation|Folder), data=COMBINED_COLDIST_NARROW, family=nbinom1 )
#check assumptions
res<-simulateResiduals(combimodc4, plot=T)
#check dispersion
par(mfrow = c(1,2))
testDispersion(res)
testZeroInflation(res)#no dispersion problems, no zero-inflation

#model simplification
combimodc4b<-glmmTMB(ColDistRound ~ poly(Generation,2)+SelectionType+(Generation|Folder), data=COMBINED_COLDIST_NARROW, family=nbinom1 )
anova(combimodc4, combimodc4b)#interaction not significant 
combimodc4c<-glmmTMB(ColDistRound ~ poly(Generation,2)+(Generation|Folder), data=COMBINED_COLDIST_NARROW, family=nbinom1 )
anova(combimodc4c, combimodc4b)#need selection type
combimodc4d<-glmmTMB(ColDistRound ~ SelectionType+(Generation|Folder), data=COMBINED_COLDIST_NARROW, family=nbinom1 )
anova(combimodc4d, combimodc4b)#can remove generation
combimodc4e<-glmmTMB(ColDistRound ~ (Generation|Folder), data=COMBINED_COLDIST_NARROW, family=nbinom1 )
anova(combimodc4d, combimodc4e)#need selection type still
```
```{r control_coldiff11, message=FALSE, warning=FALSE, results='hide', fig.show='hide'}
#without random slope
combimodc4_2<-glmmTMB(ColDistRound ~ poly(Generation,2)*SelectionType+(1|Folder), data=COMBINED_COLDIST_NARROW, family=nbinom1 )
combimodc4b_2<-glmmTMB(ColDistRound ~ poly(Generation,2)+SelectionType+(1|Folder), data=COMBINED_COLDIST_NARROW, family=nbinom1 )
anova(combimodc4_2, combimodc4b_2)#interaction not significant 
combimodc4c_2<-glmmTMB(ColDistRound ~ poly(Generation,2)+(1|Folder), data=COMBINED_COLDIST_NARROW, family=nbinom1 )
anova(combimodc4c_2, combimodc4b_2)#scenario not needed
combimodc4d_2<-glmmTMB(ColDistRound ~ 1+(1|Folder), data=COMBINED_COLDIST_NARROW, family=nbinom1 )
anova(combimodc4d_2, combimodc4c_2)#need generation
combimodc4e_2<-glmmTMB(ColDistRound ~ Generation+(1|Folder), data=COMBINED_COLDIST_NARROW, family=nbinom1 )
anova(combimodc4c_2, combimodc4e_2)#don't need polynomial
combimodc4f_2<-glmmTMB(ColDistRound ~ 1+(1|Folder), data=COMBINED_COLDIST_NARROW, family=nbinom1 )
anova(combimodc4f_2, combimodc4e_2)#generation still sig
```
```{r control_coldiff12, message=FALSE, warning=FALSE}
AIC(combimodc4e_2,combimodc4d)#random slope best
```

#### Edge disruption
Full colour space:
```{r control_ed1, message=FALSE, warning=FALSE, results='hide', fig.show='hide'}
combimodg1<-lmer(Target_L_GabRat~poly(Generation,2)*SelectionType+(Generation|Folder), data=COMBINED_FULL)
#model convergence issues
```
```{r control_ed2, message=FALSE, warning=FALSE, fig.show='hide'}
#without random slope
combimodg1i<-lmer(Target_L_GabRat~poly(Generation,2)*SelectionType+(1|Folder), data=COMBINED_FULL)
combimodg1ib<-lmer(Target_L_GabRat~poly(Generation,2)+SelectionType+(1|Folder), data=COMBINED_FULL)
anova(combimodg1i, combimodg1ib)#remove interaction - no difference between exptl and control run
combimodg1ic<-lmer(Target_L_GabRat~poly(Generation,2)+(1|Folder), data=COMBINED_FULL)
anova(combimodg1ic, combimodg1ib)#remove selection type
combimodg1id<-lmer(Target_L_GabRat~(1|Folder), data=COMBINED_FULL)
anova(combimodg1ic, combimodg1id)# no effect of generation
```
Narrow colour space:

```{r control_ed3, message=FALSE, warning=FALSE, results='hide', fig.show='hide'}
combimodg2<-lmer(Target_L_GabRat~poly(Generation,2)*SelectionType+(Generation|Folder), data=COMBINED_NARROW)
plot(combimodg2)
hist(resid(combimodg2))
#model convergence issues during simplification, so use model without random slope
```
```{r control_ed4, message=FALSE, warning=FALSE, fig.show='hide'}
#without random slope
combimodg2_2<-lmer(Target_L_GabRat~poly(Generation,2)*SelectionType+(1|Folder), data=COMBINED_NARROW)
plot(combimodg2_2)
hist(resid(combimodg2_2))
combimodg2b_2<-lmer(Target_L_GabRat~poly(Generation,2)+SelectionType+(1|Folder), data=COMBINED_NARROW)
anova(combimodg2_2, combimodg2b_2)#no significant  interaction - no difference between exptl and control runs
combimodg2c_2<-lmer(Target_L_GabRat~poly(Generation,2)+(1|Folder), data=COMBINED_NARROW)
anova(combimodg2c_2, combimodg2b_2)#significant effect of selection type, but not interacting
combimodg2d_2<-lmer(Target_L_GabRat~SelectionType+(1|Folder), data=COMBINED_NARROW)
anova(combimodg2b_2, combimodg2d_2)#generation significant
combimodg2e_2<-lmer(Target_L_GabRat~Generation+SelectionType+(1|Folder), data=COMBINED_NARROW)
anova(combimodg2e_2, combimodg2b_2)#polynomial not needed
#summary(combimodg2e_2)
combimodg2f_2<-lmer(Target_L_GabRat~SelectionType+(1|Folder), data=COMBINED_NARROW)
anova(combimodg2e_2, combimodg2f_2)#generation still significant
combimodg2g_2<-lmer(Target_L_GabRat~Generation+(1|Folder), data=COMBINED_NARROW)
anova(combimodg2e_2, combimodg2g_2)#selection type still significant
```


#### Plot results

```{r suppfig6_prep, message=FALSE, results = 'hide', echo=FALSE, warning=FALSE}
#Luminance - full colour space
COMBINED_FULL$sqrtLdiffTB<-sqrt(COMBINED_FULL$LdiffTB)
combimodl1c<-lmer(sqrtLdiffTB~poly(Generation,2)+(Generation|Folder), data=COMBINED_FULL)
cl1pred<-augment(combimodl1c)#make predictions
#add selection type
for (i in 1: length(cl1pred$Folder)){
  temp<-str_split( cl1pred$Folder[i], "_")[[1]][1]
  if(temp=="Random"){
    cl1pred$SelectionType[i]<-"Random"
  }
  else{cl1pred$SelectionType[i]<-"Human"}
  
}

cl1.sum <- cl1pred %>%
  group_by(Generation,SelectionType,Folder)%>%
  select(sqrtLdiffTB, .fitted, .fixed) %>% 
  summarise_each(funs(mean = mean, 
                      sd = sd))
cl1.sum$lCTse<-cl1.sum$sqrtLdiffTB_sd/sqrt(24)

tempcol<-c(colvec_full,"grey0","grey20","grey30","grey45","grey65", "grey85")
cl1.sum$Folder<-factor(cl1.sum$Folder, levels=c("Full_Sm_1", "Full_Sm_2", "Full_Sm_3","Full_Fr_1", "Full_Fr_2" ,"Full_Fr_3","Nar_Sm_1",  "Nar_Sm_2" , "Nar_Sm_3" ,"Nar_Fr_1" , "Nar_Fr_2",  "Nar_Fr_3",
                                                "Random_Full_Sm_1", "Random_Full_Sm_2", "Random_Full_Sm_3","Random_Full_Fr_1" ,"Random_Full_Fr_2" ,"Random_Full_Fr_3",
                                                "Random_Nar_Sm_1" , "Random_Nar_Sm_2",  "Random_Nar_Sm_3","Random_Nar_Fr_1" , "Random_Nar_Fr_2" , "Random_Nar_Fr_3" ))

indpred_cl1<-ggplot(cl1.sum, aes(x=Generation, y=sqrtLdiffTB_mean, group=Folder, col=Folder, shape=SelectionType))+  
  geom_pointrange(aes(ymin=sqrtLdiffTB_mean-lCTse, ymax=sqrtLdiffTB_mean+lCTse), alpha=0.7) +
  geom_line(aes(x=Generation, y=.fitted_mean, col=Folder),size=0.6, alpha=0.8, linetype=3)+
  geom_line(aes(x=Generation, y=.fixed_mean),size=1.5, alpha=0.5, linetype=4, col="#BD0026")+
  scale_colour_manual(values=tempcol)+
  scale_shape_manual(values=c(19,17))+
  my_theme+
  guides(col=FALSE)+
  guides(linetype=FALSE)+
  guides(shape=FALSE)+
  xlab("Generation")+ylab("sqrt(L difference)")+
  theme(axis.title = element_text(size = 14))  +
  theme(axis.text = element_text(size = 12))  +
  ggtitle("(i) Luminance difference")+
  theme(plot.title = element_text(face = "italic"))
  
#Luminance - narrow colour space
COMBINED_NARROW$sqrtLdiffTB<-sqrt(COMBINED_NARROW$LdiffTB)
combimodl2c<-lmer(sqrtLdiffTB~Generation*SelectionType+(Generation|Folder), data=COMBINED_NARROW)
cl2pred<-augment(combimodl2c)#make predictions

cl2.sum <- cl2pred %>%
  group_by(Generation,SelectionType,Folder)%>%
  select(sqrtLdiffTB, .fitted, .fixed) %>% 
  summarise_each(funs(mean = mean, 
                      sd = sd))
cl2.sum$lCTse<-cl2.sum$sqrtLdiffTB_sd/sqrt(24)

tempcol2<-c(colvec_narrow,"grey0","grey20","grey30","grey45","grey65", "grey85")
cl2.sum$Folder<-factor(cl2.sum$Folder, levels=c("Full_Sm_1", "Full_Sm_2", "Full_Sm_3","Full_Fr_1", "Full_Fr_2" ,"Full_Fr_3","Nar_Sm_1",  "Nar_Sm_2" , "Nar_Sm_3" ,"Nar_Fr_1" , "Nar_Fr_2",  "Nar_Fr_3",
                                                "Random_Full_Sm_1", "Random_Full_Sm_2", "Random_Full_Sm_3","Random_Full_Fr_1" ,"Random_Full_Fr_2" ,"Random_Full_Fr_3",
                                                "Random_Nar_Sm_1" , "Random_Nar_Sm_2",  "Random_Nar_Sm_3","Random_Nar_Fr_1" , "Random_Nar_Fr_2" , "Random_Nar_Fr_3" ))

indpred_cl2<-ggplot(cl2.sum, aes(x=Generation, y=sqrtLdiffTB_mean, group=Folder, col=Folder, shape=SelectionType))+  
  geom_pointrange(aes(ymin=sqrtLdiffTB_mean-lCTse, ymax=sqrtLdiffTB_mean+lCTse), alpha=0.7) +
  geom_line(aes(x=Generation, y=.fitted_mean, col=Folder),size=0.6, alpha=0.8, linetype=3)+
  geom_line(aes(x=Generation, y=.fixed_mean, linetype=SelectionType),size=1.5, alpha=0.5,  col="#245D99")+
  scale_colour_manual(values=tempcol2)+
  scale_shape_manual(values=c(19,17))+
  scale_linetype_manual(values=c(1,2))+
  my_theme+
  guides(col=FALSE)+
  guides(linetype=FALSE)+
  guides(shape=FALSE)+
  xlab("Generation")+ylab("sqrt(L difference)")+
  theme(axis.title = element_text(size = 14))  +
  theme(axis.text = element_text(size = 12)) +
  ggtitle("(i) Luminance difference")+
  theme(plot.title = element_text(face = "italic"))

#Colour - full colour space
COMBINED_FULL$sqrteucdisTB<-sqrt(COMBINED_FULL$eucdisTB)
combimodc1c<-lmer(sqrteucdisTB~poly(Generation,2)+(Generation|Folder), data=COMBINED_FULL)
cc1pred<-augment(combimodc1c)#make predictions
#add selection type
for (i in 1: length(cc1pred$Folder)){
  temp<-str_split( cc1pred$Folder[i], "_")[[1]][1]
  if(temp=="Random"){
    cc1pred$SelectionType[i]<-"Random"
  }
  else{cc1pred$SelectionType[i]<-"Human"}
  
}

cc1.sum <- cc1pred %>%
  group_by(Generation,SelectionType,Folder)%>%
  select(sqrteucdisTB, .fitted, .fixed) %>% 
  summarise_each(funs(mean = mean, 
                      sd = sd))
cc1.sum$lCTse<-cc1.sum$sqrteucdisTB_sd/sqrt(24)

tempcol<-c(colvec_full,"grey0","grey20","grey30","grey45","grey65", "grey85")
cc1.sum$Folder<-factor(cc1.sum$Folder, levels=c("Full_Sm_1", "Full_Sm_2", "Full_Sm_3","Full_Fr_1", "Full_Fr_2" ,"Full_Fr_3","Nar_Sm_1",  "Nar_Sm_2" , "Nar_Sm_3" ,"Nar_Fr_1" , "Nar_Fr_2",  "Nar_Fr_3",
                                                "Random_Full_Sm_1", "Random_Full_Sm_2", "Random_Full_Sm_3","Random_Full_Fr_1" ,"Random_Full_Fr_2" ,"Random_Full_Fr_3",
                                                "Random_Nar_Sm_1" , "Random_Nar_Sm_2",  "Random_Nar_Sm_3","Random_Nar_Fr_1" , "Random_Nar_Fr_2" , "Random_Nar_Fr_3" ))

indpred_cc1<-ggplot(cc1.sum, aes(x=Generation, y=sqrteucdisTB_mean, group=Folder, col=Folder, shape=SelectionType))+  
  geom_pointrange(aes(ymin=sqrteucdisTB_mean-lCTse, ymax=sqrteucdisTB_mean+lCTse), alpha=0.7) +
  geom_line(aes(x=Generation, y=.fitted_mean, col=Folder),size=0.6, alpha=0.8, linetype=3)+
  geom_line(aes(x=Generation, y=.fixed_mean),size=1.5, alpha=0.5, linetype=4, col="#BD0026")+
  scale_colour_manual(values=tempcol)+
  scale_shape_manual(values=c(19,17))+
  my_theme+
  guides(col=FALSE)+
  guides(linetype=FALSE)+
  guides(shape=FALSE)+
  xlab("Generation")+ylab("sqrt(Euclidean distance in CIELAB)")+
  theme(axis.title = element_text(size = 14))  +
  theme(axis.text = element_text(size = 12))  +
  ggtitle("(ii) Colour difference")+
  theme(plot.title = element_text(face = "italic"))

#Colour - narrow colour space
COMBINED_NARROW$sqrteucdisTB<-sqrt(COMBINED_NARROW$eucdisTB)
combimodc2c<-lmer(sqrteucdisTB~Generation*SelectionType+(Generation|Folder), data=COMBINED_NARROW)
cc2pred<-augment(combimodc2c)#make predictions

cc2.sum <- cc2pred %>%
  group_by(Generation,SelectionType,Folder)%>%
  select(sqrteucdisTB, .fitted, .fixed) %>% 
  summarise_each(funs(mean = mean, 
                      sd = sd))
cc2.sum$lCTse<-cc2.sum$sqrteucdisTB_sd/sqrt(24)

tempcol2<-c(colvec_narrow,"grey0","grey20","grey30","grey45","grey65", "grey85")

cc2.sum$Folder<-factor(cc2.sum$Folder, levels=c("Full_Sm_1", "Full_Sm_2", "Full_Sm_3","Full_Fr_1", "Full_Fr_2" ,"Full_Fr_3","Nar_Sm_1",  "Nar_Sm_2" , "Nar_Sm_3" ,"Nar_Fr_1" , "Nar_Fr_2",  "Nar_Fr_3",
                                                "Random_Full_Sm_1", "Random_Full_Sm_2", "Random_Full_Sm_3","Random_Full_Fr_1" ,"Random_Full_Fr_2" ,"Random_Full_Fr_3",
                                                "Random_Nar_Sm_1" , "Random_Nar_Sm_2",  "Random_Nar_Sm_3","Random_Nar_Fr_1" , "Random_Nar_Fr_2" , "Random_Nar_Fr_3" ))

indpred_cc2<-ggplot(cc2.sum, aes(x=Generation, y=sqrteucdisTB_mean, group=Folder, col=Folder, shape=SelectionType))+  
  geom_pointrange(aes(ymin=sqrteucdisTB_mean-lCTse, ymax=sqrteucdisTB_mean+lCTse), alpha=0.7) +
  geom_line(aes(x=Generation, y=.fitted_mean, col=Folder),size=0.6, alpha=0.8, linetype=3)+
  geom_line(aes(x=Generation, y=.fixed_mean, linetype=SelectionType),size=1.5, alpha=0.5,  col="#245D99")+
  scale_colour_manual(values=tempcol2)+
  scale_shape_manual(values=c(19,17))+
  scale_linetype_manual(values=c(1,2))+
  my_theme+
  guides(col=FALSE)+
  guides(linetype=FALSE)+
  guides(shape=FALSE)+
  xlab("Generation")+ylab("sqrt(Euclidean distance in CIELAB)")+
  theme(axis.title = element_text(size = 14))  +
  theme(axis.text = element_text(size = 12)) +
  ggtitle("(ii) Colour difference")+
  theme(plot.title = element_text(face = "italic"))

#Colour, weighted average - full colour space
combimodc3e<-lmer(NewColMap_DeltaS~Generation+(Generation|Folder), data=COMBINED_COLDIST_FULL)
cc3pred<-augment(combimodc3e)#make predictions
#add selection type
for (i in 1: length(cc3pred$Folder)){
  temp<-str_split( cc3pred$Folder[i], "_")[[1]][1]
  if(temp=="Random"){
    cc3pred$SelectionType[i]<-"Random"
  }
  else{cc3pred$SelectionType[i]<-"Human"}
  
}

cc3.sum <- cc3pred %>%
  group_by(Generation,SelectionType,Folder)%>%
  select(NewColMap_DeltaS, .fitted, .fixed) %>% 
  summarise_each(funs(mean = mean, 
                      sd = sd))
cc3.sum$lCTse<-cc3.sum$NewColMap_DeltaS_sd/sqrt(24)

tempcol<-c(colvec_full,"grey0","grey20","grey30","grey45","grey65", "grey85")
cc3.sum$Folder<-factor(cc3.sum$Folder, levels=c("Full_Sm_1", "Full_Sm_2", "Full_Sm_3","Full_Fr_1", "Full_Fr_2" ,"Full_Fr_3","Nar_Sm_1",  "Nar_Sm_2" , "Nar_Sm_3" ,"Nar_Fr_1" , "Nar_Fr_2",  "Nar_Fr_3",
                                                "Random_Full_Sm_1", "Random_Full_Sm_2", "Random_Full_Sm_3","Random_Full_Fr_1" ,"Random_Full_Fr_2" ,"Random_Full_Fr_3",
                                                "Random_Nar_Sm_1" , "Random_Nar_Sm_2",  "Random_Nar_Sm_3","Random_Nar_Fr_1" , "Random_Nar_Fr_2" , "Random_Nar_Fr_3" ))

indpred_cc3<-ggplot(cc3.sum, aes(x=Generation, y=NewColMap_DeltaS_mean, group=Folder, col=Folder, shape=SelectionType))+  
  geom_pointrange(aes(ymin=NewColMap_DeltaS_mean-lCTse, ymax=NewColMap_DeltaS_mean+lCTse), alpha=0.7) +
  geom_line(aes(x=Generation, y=.fitted_mean, col=Folder),size=0.6, alpha=0.8, linetype=3)+
  geom_line(aes(x=Generation, y=.fixed_mean),size=1.5, alpha=0.5, linetype=4, col="#BD0026")+
  scale_colour_manual(values=tempcol)+
  scale_shape_manual(values=c(19,17))+
  my_theme+
  guides(col=FALSE)+
  guides(linetype=FALSE)+
  guides(shape=FALSE)+
  xlab("Generation")+ylab("Colour difference (Delta-S)")+
  theme(axis.title = element_text(size = 14))  +
  theme(axis.text = element_text(size = 12))  +
  ggtitle("(iii) Weighted average colour difference")+
  theme(plot.title = element_text(face = "italic"))

#Colour, weighted average - narrow colour space
combimodc4d<-glmmTMB(ColDistRound ~ SelectionType+(Generation|Folder), data=COMBINED_COLDIST_NARROW, family=nbinom1 )
cnccpred1 <- as.data.frame(ggpredict(combimodc4d, terms = c("Generation[all]","SelectionType[Human]","Folder[Nar_Sm_1]"), type="random"))
cnccpred2 <- as.data.frame(ggpredict(combimodc4d, terms = c("Generation[all]","SelectionType[Human]","Folder[Nar_Sm_2]"), type="random"))
cnccpred3 <- as.data.frame(ggpredict(combimodc4d, terms = c("Generation[all]","SelectionType[Human]","Folder[Nar_Sm_3]"), type="random"))
cnccpred4 <- as.data.frame(ggpredict(combimodc4d, terms = c("Generation[all]","SelectionType[Human]","Folder[Nar_Fr_1]"), type="random"))
cnccpred5 <- as.data.frame(ggpredict(combimodc4d, terms = c("Generation[all]","SelectionType[Human]","Folder[Nar_Fr_2]"), type="random"))
cnccpred6 <- as.data.frame(ggpredict(combimodc4d, terms = c("Generation[all]","SelectionType[Human]","Folder[Nar_Fr_3]"), type="random"))

cnccpred7 <- as.data.frame(ggpredict(combimodc4d, terms = c("Generation[all]","SelectionType[Random]","Folder[Random_Nar_Sm_1]"), type="random"))
cnccpred8 <- as.data.frame(ggpredict(combimodc4d, terms = c("Generation[all]","SelectionType[Random]","Folder[Random_Nar_Sm_2]"), type="random"))
cnccpred9 <- as.data.frame(ggpredict(combimodc4d, terms = c("Generation[all]","SelectionType[Random]","Folder[Random_Nar_Sm_3]"), type="random"))
cnccpred10 <- as.data.frame(ggpredict(combimodc4d, terms = c("Generation[all]","SelectionType[Random]","Folder[Random_Nar_Fr_1]"), type="random"))
cnccpred11 <- as.data.frame(ggpredict(combimodc4d, terms = c("Generation[all]","SelectionType[Random]","Folder[Random_Nar_Fr_2]"), type="random"))
cnccpred12 <- as.data.frame(ggpredict(combimodc4d, terms = c("Generation[all]","SelectionType[Random]","Folder[Random_Nar_Fr_3]"), type="random"))

#and predictions for fixed effects only
cnccpred1_fonly <- as.data.frame(ggpredict(combimodc4d, terms = c("Generation[all]","SelectionType[Human]"), type="fixed"))
cnccpred2_fonly <- as.data.frame(ggpredict(combimodc4d, terms = c("Generation[all]","SelectionType[Random]"), type="fixed"))


combicoldist_sum<-COMBINED_COLDIST_NARROW%>%
  group_by(SelectionType, Folder,Generation)%>%
  summarise(mean=mean(ColDistRound),sd=sd(ColDistRound))
combicoldist_sum<-as.data.frame(combicoldist_sum)
combicoldist_sum$Folder<-factor(combicoldist_sum$Folder, levels=c("Full_Sm_1", "Full_Sm_2", "Full_Sm_3","Full_Fr_1", "Full_Fr_2" ,"Full_Fr_3","Nar_Sm_1",  "Nar_Sm_2" , "Nar_Sm_3" ,"Nar_Fr_1" , "Nar_Fr_2",  "Nar_Fr_3",
                                                                  "Random_Full_Sm_1", "Random_Full_Sm_2", "Random_Full_Sm_3","Random_Full_Fr_1" ,"Random_Full_Fr_2" ,"Random_Full_Fr_3",
                                                                  "Random_Nar_Sm_1" , "Random_Nar_Sm_2",  "Random_Nar_Sm_3","Random_Nar_Fr_1" , "Random_Nar_Fr_2" , "Random_Nar_Fr_3" ))

for(i in 1:length(combicoldist_sum$Generation)){
  targetgen<-combicoldist_sum$Generation[i]
  
  if(combicoldist_sum$Folder[i]=="Nar_Sm_1"){
    predict_val<-cnccpred1
    
    for(j in 1:length(predict_val$predicted)){
      if(predict_val$x[j]==targetgen){
        combicoldist_sum$predicted[i]<-predict_val$predicted[j]
      }
      
    }
  }
  if(combicoldist_sum$Folder[i]=="Nar_Sm_2"){
    predict_val<-cnccpred2
    
    for(j in 1:length(predict_val$predicted)){
      if(predict_val$x[j]==targetgen){
        combicoldist_sum$predicted[i]<-predict_val$predicted[j]
      }
      
    }
  }
  if(combicoldist_sum$Folder[i]=="Nar_Sm_3"){
    predict_val<-cnccpred3
    
    for(j in 1:length(predict_val$predicted)){
      if(predict_val$x[j]==targetgen){
        combicoldist_sum$predicted[i]<-predict_val$predicted[j]
      }
      
    }
  }
  if(combicoldist_sum$Folder[i]=="Nar_Fr_1"){
    predict_val<-cnccpred4
    
    for(j in 1:length(predict_val$predicted)){
      if(predict_val$x[j]==targetgen){
        combicoldist_sum$predicted[i]<-predict_val$predicted[j]
      }
      
    }
  }
  if(combicoldist_sum$Folder[i]=="Nar_Fr_2"){
    predict_val<-cnccpred5
    
    for(j in 1:length(predict_val$predicted)){
      if(predict_val$x[j]==targetgen){
        combicoldist_sum$predicted[i]<-predict_val$predicted[j]
      }
      
    }
  }
  if(combicoldist_sum$Folder[i]=="Nar_Fr_3"){
    predict_val<-cnccpred6
    
    for(j in 1:length(predict_val$predicted)){
      if(predict_val$x[j]==targetgen){
        combicoldist_sum$predicted[i]<-predict_val$predicted[j]
      }
      
    }
  }
  
  if(combicoldist_sum$Folder[i]=="Random_Nar_Sm_1"){
    predict_val<-cnccpred7
    
    for(j in 1:length(predict_val$predicted)){
      if(predict_val$x[j]==targetgen){
        combicoldist_sum$predicted[i]<-predict_val$predicted[j]
      }
      
    }
  }
  if(combicoldist_sum$Folder[i]=="Random_Nar_Sm_2"){
    predict_val<-cnccpred8
    
    for(j in 1:length(predict_val$predicted)){
      if(predict_val$x[j]==targetgen){
        combicoldist_sum$predicted[i]<-predict_val$predicted[j]
      }
      
    }
  }
  if(combicoldist_sum$Folder[i]=="Random_Nar_Sm_3"){
    predict_val<-cnccpred9
    
    for(j in 1:length(predict_val$predicted)){
      if(predict_val$x[j]==targetgen){
        combicoldist_sum$predicted[i]<-predict_val$predicted[j]
      }
      
    }
  }
  if(combicoldist_sum$Folder[i]=="Random_Nar_Fr_1"){
    predict_val<-cnccpred10
    
    for(j in 1:length(predict_val$predicted)){
      if(predict_val$x[j]==targetgen){
        combicoldist_sum$predicted[i]<-predict_val$predicted[j]
      }
      
    }
  }
  if(combicoldist_sum$Folder[i]=="Random_Nar_Fr_2"){
    predict_val<-cnccpred11
    
    for(j in 1:length(predict_val$predicted)){
      if(predict_val$x[j]==targetgen){
        combicoldist_sum$predicted[i]<-predict_val$predicted[j]
      }
      
    }
  }
  if(combicoldist_sum$Folder[i]=="Random_Nar_Fr_3"){
    predict_val<-cnccpred12
    
    for(j in 1:length(predict_val$predicted)){
      if(predict_val$x[j]==targetgen){
        combicoldist_sum$predicted[i]<-predict_val$predicted[j]
      }
      
    }
  }
  
  if(combicoldist_sum$SelectionType[i]=="Human"){
    predict_fixed<-cnccpred1_fonly
    for(k in 1: length(predict_fixed$predicted)){
      if(predict_fixed$x[k]==targetgen){
        combicoldist_sum$fixed[i]<-predict_fixed$predicted[k]
      }
    }
  }
  else {
    predict_fixed<-cnccpred2_fonly
    for(k in 1: length(predict_fixed$predicted)){
      if(predict_fixed$x[k]==targetgen){
        combicoldist_sum$fixed[i]<-predict_fixed$predicted[k]
      }
    }
  }
  
}
combicoldist_sum$se<-combicoldist_sum$sd/sqrt(24)
tempcol2<-c(colvec_narrow,"grey0","grey20","grey30","grey45","grey65", "grey85")

indpred_cc4<-ggplot(combicoldist_sum, aes(x=Generation, y=mean, group=Folder, col=Folder))+  
  geom_pointrange(aes(ymin=mean-se, ymax=mean+se, shape=SelectionType), alpha=1,position = position_dodge(width = 0.3), fill="white") +
  geom_line(aes(x=Generation, y=predicted),size=0.6, alpha=0.8, linetype=3)+
  geom_line(aes(x=Generation, y=fixed, linetype=SelectionType), size=1.5,  col="#245D99")+
  scale_colour_manual(values=tempcol2)+
  scale_shape_manual(values=c(19,17))+
  scale_linetype_manual(values=c(1,2))+
  my_theme+
  guides(col=FALSE)+guides(linetype=FALSE)+guides(shape=FALSE)+
  xlab("Generation")+ylab("Colour difference (Delta-S)")+
  theme(axis.title = element_text(size = 14))  +
  theme(axis.text = element_text(size = 12)) +
  ggtitle("(iii) Weighted average colour difference") +
  theme(plot.title = element_text(face = "italic"))

#Edge disruption - full colour space
combimodg1d<-lmer(Target_L_GabRat~(1|Folder), data=COMBINED_FULL)
cg1pred<-augment(combimodg1d)#make predictions
#add selection type
for (i in 1: length(cg1pred$Folder)){
  temp<-str_split( cg1pred$Folder[i], "_")[[1]][1]
  if(temp=="Random"){
    cg1pred$SelectionType[i]<-"Random"
  }
  else{cg1pred$SelectionType[i]<-"Human"}
  
}
cg1pred$Generation<-COMBINED_FULL$Generation
cg1.sum <- cg1pred %>%
  group_by(Generation,SelectionType,Folder)%>%
  select(Target_L_GabRat, .fitted, .fixed) %>% 
  summarise_each(funs(mean = mean, 
                      sd = sd))
cg1.sum$lCTse<-cg1.sum$Target_L_GabRat_sd/sqrt(24)

tempcol<-c(colvec_full,"grey0","grey20","grey30","grey45","grey65", "grey85")
cg1.sum$Folder<-factor(cg1.sum$Folder, levels=c("Full_Sm_1", "Full_Sm_2", "Full_Sm_3","Full_Fr_1", "Full_Fr_2" ,"Full_Fr_3","Nar_Sm_1",  "Nar_Sm_2" , "Nar_Sm_3" ,"Nar_Fr_1" , "Nar_Fr_2",  "Nar_Fr_3",
                                                "Random_Full_Sm_1", "Random_Full_Sm_2", "Random_Full_Sm_3","Random_Full_Fr_1" ,"Random_Full_Fr_2" ,"Random_Full_Fr_3",
                                                "Random_Nar_Sm_1" , "Random_Nar_Sm_2",  "Random_Nar_Sm_3","Random_Nar_Fr_1" , "Random_Nar_Fr_2" , "Random_Nar_Fr_3" ))

indpred_cg1<-ggplot(cg1.sum, aes(x=Generation, y=Target_L_GabRat_mean, group=Folder, col=Folder, shape=SelectionType))+  
  geom_pointrange(aes(ymin=Target_L_GabRat_mean-lCTse, ymax=Target_L_GabRat_mean+lCTse), alpha=0.7) +
  geom_line(aes(x=Generation, y=.fitted_mean, col=Folder),size=0.6, alpha=0.8, linetype=3)+
 # geom_line(aes(x=Generation, y=.fixed_mean),size=1.5, alpha=0.5, linetype=4, col="#BD0026")+
  scale_colour_manual(values=tempcol)+
  scale_shape_manual(values=c(19,17))+
  my_theme+
  guides(col=FALSE)+
  guides(linetype=FALSE)+
  guides(shape=FALSE)+
  xlab("Generation")+ylab("GabRat in L")+
  theme(axis.title = element_text(size = 14))  +
  theme(axis.text = element_text(size = 12))  +
  ggtitle("(iv) Edge disruption")+
  theme(plot.title = element_text(face = "italic"))

#Edge disruption - narrow colour space
combimodg2d<-lmer(Target_L_GabRat~Generation + SelectionType+(1|Folder), data=COMBINED_NARROW)
cg2pred<-augment(combimodg2d)#make predictions

cg2.sum <- cg2pred %>%
  group_by(Generation,SelectionType,Folder)%>%
  select(Target_L_GabRat, .fitted, .fixed) %>% 
  summarise_each(funs(mean = mean, 
                      sd = sd))
cg2.sum$lCTse<-cg2.sum$Target_L_GabRat_sd/sqrt(24)

tempcol2<-c(colvec_narrow,"grey0","grey20","grey30","grey45","grey65", "grey85")
cg2.sum$Folder<-factor(cg2.sum$Folder, levels=c("Full_Sm_1", "Full_Sm_2", "Full_Sm_3","Full_Fr_1", "Full_Fr_2" ,"Full_Fr_3","Nar_Sm_1",  "Nar_Sm_2" , "Nar_Sm_3" ,"Nar_Fr_1" , "Nar_Fr_2",  "Nar_Fr_3",
                                                "Random_Full_Sm_1", "Random_Full_Sm_2", "Random_Full_Sm_3","Random_Full_Fr_1" ,"Random_Full_Fr_2" ,"Random_Full_Fr_3",
                                                "Random_Nar_Sm_1" , "Random_Nar_Sm_2",  "Random_Nar_Sm_3","Random_Nar_Fr_1" , "Random_Nar_Fr_2" , "Random_Nar_Fr_3" ))

indpred_cg2<-ggplot(cg2.sum, aes(x=Generation, y=Target_L_GabRat_mean, group=Folder, col=Folder, shape=SelectionType))+  
  geom_pointrange(aes(ymin=Target_L_GabRat_mean-lCTse, ymax=Target_L_GabRat_mean+lCTse), alpha=0.7, position=position_dodge(width=0.3)) +
  geom_line(aes(x=Generation, y=.fitted_mean, col=Folder),size=0.6, alpha=0.8, linetype=3)+
  geom_line(aes(x=Generation, y=.fixed_mean, linetype=SelectionType),size=1.5, alpha=0.5, col="#245D99")+
  scale_colour_manual(values=tempcol2)+
  scale_shape_manual(values=c(19,17))+
  scale_linetype_manual(values=c(1,2))+
  my_theme+
  guides(col=FALSE)+
  guides(linetype=FALSE)+
  guides(shape=FALSE)+
  xlab("Generation")+ylab("GabRat in L")+
  theme(axis.title = element_text(size = 14))  +
  theme(axis.text = element_text(size = 12))  +
  ggtitle("(iv) Edge disruption")+
  theme(plot.title = element_text(face = "italic"))

```

```{r suppfig6, message=FALSE, results = 'hide', echo=FALSE, warning=FALSE, fig.dim=c(20,30), fig.cap="Supplementary Figure 6: Changes in camouflage metrics in the screen-based experiments, experimental and control populations"}
(indpred_cl2+indpred_cl1)/
  (indpred_cc2+indpred_cc1)/
  (indpred_cc4+indpred_cc3)/
  (indpred_cg2+indpred_cg1)

```
````
